# The genetic basis of tail-loss evolution in humans and apes

**DOI:** 10.1101/2021.09.14.460388

**Authors:** Bo Xia, Weimin Zhang, Aleksandra Wudzinska, Emily Huang, Ran Brosh, Maayan Pour, Alexander Miller, Jeremy S. Dasen, Matthew T. Maurano, Sang Y. Kim, Jef D. Boeke, Itai Yanai

**Affiliations:** Institute for Computational Medicine, NYU Langone Health, New York, NY 10016, USA; Institute for Systems Genetics, NYU Langone Health, New York, NY 10016, USA; Department of Biochemistry and Molecular Pharmacology, NYU Langone Health, New York, NY 10016, USA; Department of Neuroscience and Physiology, NYU Langone Health, New York, NY 10016, USA; Department of Pathology, NYU Langone Health, New York, NY 10016, USA; Department of Biomedical Engineering, NYU Tandon School of Engineering, Brooklyn, NY, 11201, USA

## Abstract

The loss of the tail is one of the main anatomical evolutionary changes to have occurred along the lineage leading to humans and to the “anthropomorphous apes”^1,2^. This morphological reprogramming in the ancestral hominoids has been long considered to have accommodated a characteristic style of locomotion and contributed to the evolution of bipedalism in humans^3–5^. Yet, the precise genetic mechanism that facilitated tail-loss evolution in hominoids remains unknown. Primate genome sequencing projects have made possible the identification of causal links between genotypic and phenotypic changes^6–8^, and enable the search for hominoid-specific genetic elements controlling tail development^9^. Here, we present evidence that tail-loss evolution was mediated by the insertion of an individual *Alu* element into the genome of the hominoid ancestor. We demonstrate that this *Alu* element – inserted into an intron of the *TBXT* gene (also called *T* or *Brachyury*^10–12^) – pairs with a neighboring ancestral *Alu* element encoded in the reverse genomic orientation and leads to a hominoid-specific alternative splicing event. To study the effect of this splicing event, we generated a mouse model that mimics the expression of human *TBXT* products by expressing both full-length and exon-skipped isoforms of the mouse *TBXT* ortholog. We found that mice with this genotype exhibit the complete absence of a tail or a shortened tail, supporting the notion that the exon-skipped transcript is sufficient to induce a tail-loss phenotype, albeit with incomplete penetrance. We further noted that mice homozygous for the exon-skipped isoforms exhibited embryonic spinal cord malformations, resembling a neural tube defect condition, which affects ∼1/1000 human neonates^13^. We propose that selection for the loss of the tail along the hominoid lineage was associated with an adaptive cost of potential neural tube defects and that this ancient evolutionary trade-off may thus continue to affect human health today.

---

The tail appendage varies widely in its morphology and function across vertebrate species^4,5^. For primates in particular, the tail is adapted to a range of environments, with implications for the animal’s style of locomotion^14,15^. The New World howler monkeys, for example, evolved a prehensile tail that helps the animal to grasp or hold objects while occupying arboreal habitats^16^. Hominoids – which include humans and the apes – however, are distinct among the primates in their loss of an external tail (**Fig. 1a**). The loss of the tail is inferred to have occurred ∼25 million years ago when the hominoid lineage diverged from the ancient Old World monkeys (**Fig. 1a**), leaving only 3-4 caudal vertebrae to form the coccyx, or tailbone, in modern humans^17^. It has long been speculated that tail loss in hominoids has contributed to bipedal locomotion, whose evolutionary occurrence coincided with the loss of tail^18–20^. Recent progress in developmental biology has led to the elucidation of the gene regulatory networks that underlie tail development^9,21^. Specifically, the absence of the tail phenotype in the Mouse Genome Informatics has so far recorded 31 genes from the study of mutants and naturally occurring variants^21,22^ (**Supp. Table 1**). Expression of these genes is enriched in the development of the primitive streak and posterior body formation, including the core gene regulation network for inducing the mesoderm and definitive endoderm such as *Tbxt, Wnt3a*, and *Msgn1*. While these genes and their relationships have been studied, the exact genetic changes that drove the evolution of tail-loss in hominoids remain unknown, preventing an understanding of how tail loss affected other human evolutionary events, such as bipedalism.

**Fig. 1.**
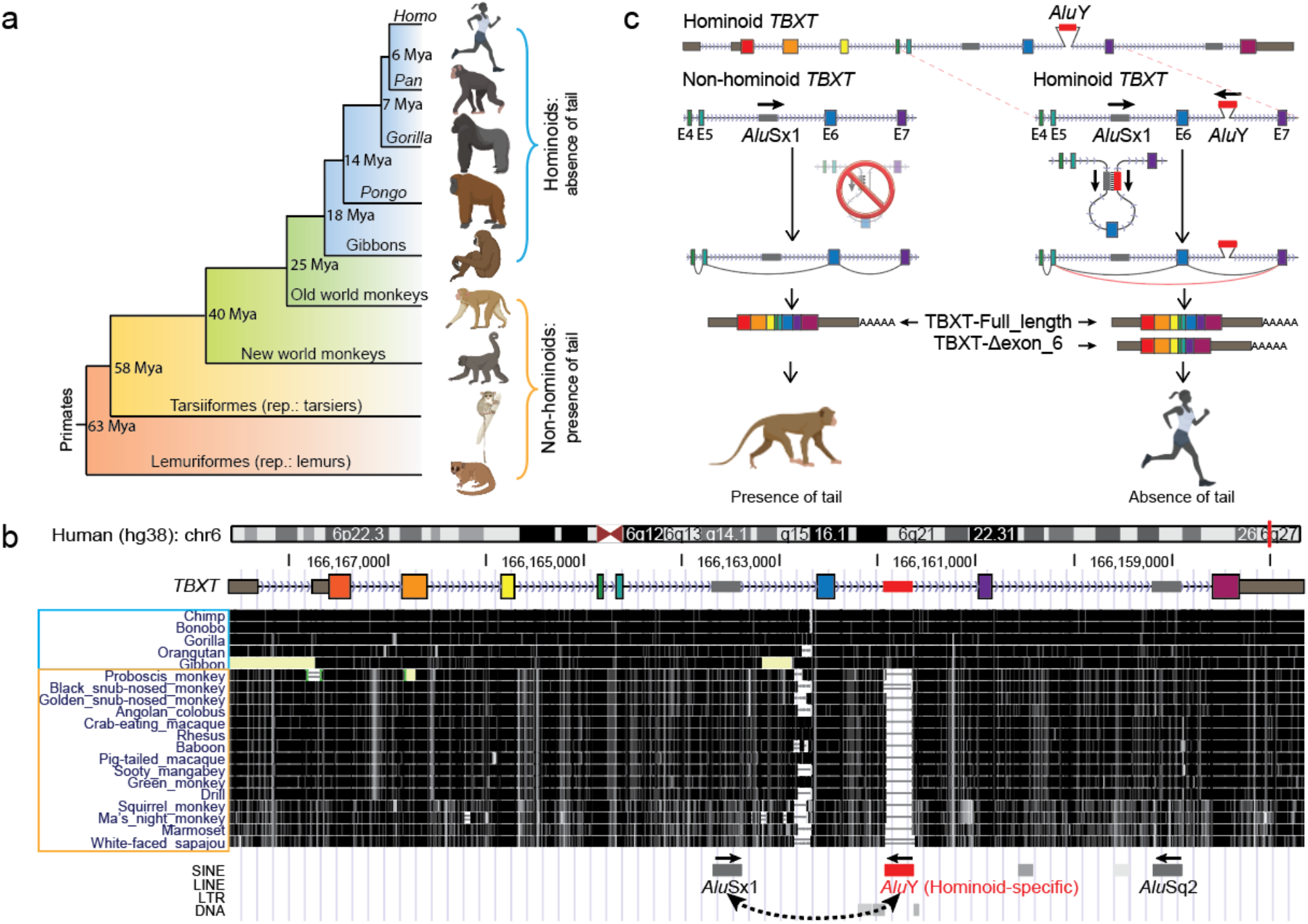
Evolution of tail loss in hominoids. **a**, Tail phenotypes across the primate phylogenetic tree. **b**, UCSC Genome browser view of the conservation score through multi-species alignment at the *TBXT* locus across primate genomes^36^. The hominoid-specific *Alu*Y element is labelled in red. **c**, Schematic of the hypothesized mechanism of tail-loss evolution in hominoids.

## Results

### A hominoid-specific intronic *AluY* element in *TBXT*

We screened through the 31 human genes – and their primate orthologs – involved in tail development, with the goal of identifying a genetic variation associated with the loss of the tail in hominoids (**Supp. Table 1**). We first examined protein sequence conservation between the hominoid genomes and its closest sister lineage, the Old World monkeys (Cercopithecidae). However, we failed to detect candidate variants in hominoid coding sequences that might provide a genetic mechanism for tail-loss evolution (**Supp. File 1**). We next queried for hominoid-specific genomic rearrangements in the non-coding regions of genes related to tail development. Surprisingly, we found a hominoid-specific *Alu* element inserted in intron 6 of *TBXT* (**Fig. 1b and Supp. File 1**)^10,11^. *TBXT* codes a highly-conserved transcription factor critical for mesoderm and definitive endoderm formation during embryonic development^12,23–25^. Heterozygous mutations in the coding regions of the *TBXT* orthologs in tailed animals, such as mouse^10^, Manx cat^26^, dog^27^ and zebrafish^28^, lead to the absence or reduced form of the tail, and homozygous mutants are typically not viable. Moreover, this particular *Alu* insertion is from the *Alu*Y subfamily, a relatively ‘young’ but not human-specific subfamily shared between the genomes of hominoids and Old World monkeys, the activity of which coincides with the evolutionary time when early hominoids lost their tails^29^.

The *Alu*Y element in *TBXT* is not inserted in the vicinity of a splice site; rather, it is >500 bp from exon 6 of *TBXT*, the nearest coding exon. As such, it would not be expected to lead to an alternative splicing event, as found for other intronic *Alu* elements affecting splicing^30–32^. However, we noted the presence of another *Alu* element (*Alu*Sx1) in the reverse orientation in intron 5 of *TBXT*, that is conserved in all simians. Together, the *Alu*Y and *Alu*Sx1 elements form an inverted repeat pair (**Fig. 1b**). We thus posited that upon transcription, the simian-specific *Alu*Sx1 element pairs with the hominoid-specific *Alu*Y element, forming a stem-loop structure in the *TBXT* pre-mRNA and trapping exon 6 in the loop (**Fig. 1c**). An inferred RNA secondary structure model supported an interaction between these two *Alu* elements^33^ (**Fig. S1**). The secondary structure of the transcript may thus conjoin the splice donor and receptor of exons 5 and 7, respectively, and promote the skipping of exon 6, leading to a hominoid-specific and in-frame alternative splicing isoform, *TBXT-Δexon6* (**Fig. 1c**). Indeed, we validated the existence of *TBXT-Δexon6* transcripts in human and its corresponding absence in mouse, which lacks the *Alu* elements, using an embryonic stem cells (ESCs) *in vitro* differentiation system that induces *TBXT* expression similar to that present in the primitive streak of the embryo (**Fig. S2**)^34,35^. Considering the high conservation of *TBXT* exon 6 and its potential transcriptional regulation function (**Fig. S3**), we thus hypothesized that in humans and apes, the TBXT-Δexon6 isoform protein disrupts tail elongation during embryonic development, leading to the reduction or loss of an external tail (**Fig. 1c**).

### *Alu*Y insertion in *TBXT* induces alternative splicing, and requires interaction with *Alu*Sx1

To test whether both *Alu*Y and *Alu*Sx1 are required to induce the hominoid-specific alternatively spliced isoform of *TBXT*, we used CRISPR/Cas9 in human ESCs to individually delete the hominoid-specific *Alu*Y element and – in a separate line – its potentially interacting counterpart *Alu*Sx1 (**Fig. 2a, S4a**). Again, we adapted the hESC *in vitro* differentiation system to mimic the *TBXT* expression in the embryo (**Fig. S2**)^34^. We found that deleting the *Alu*Y almost completely eliminated the generation of the *TBXT-Δexon6* isoform transcript (**Fig. 2b**, middle). Similarly, deleting the interacting partner, *Alu*Sx1, sufficed to repress this alternatively spliced isoform (**Fig. 2b**, right). These results support the notion that the hominoid-specific *Alu*Y insertion induces a novel *TBXT-Δexon6* AS isoform, through an interaction with the neighboring *Alu*Sx1 element (**Fig. 2c**, top).

**Fig. 2.**
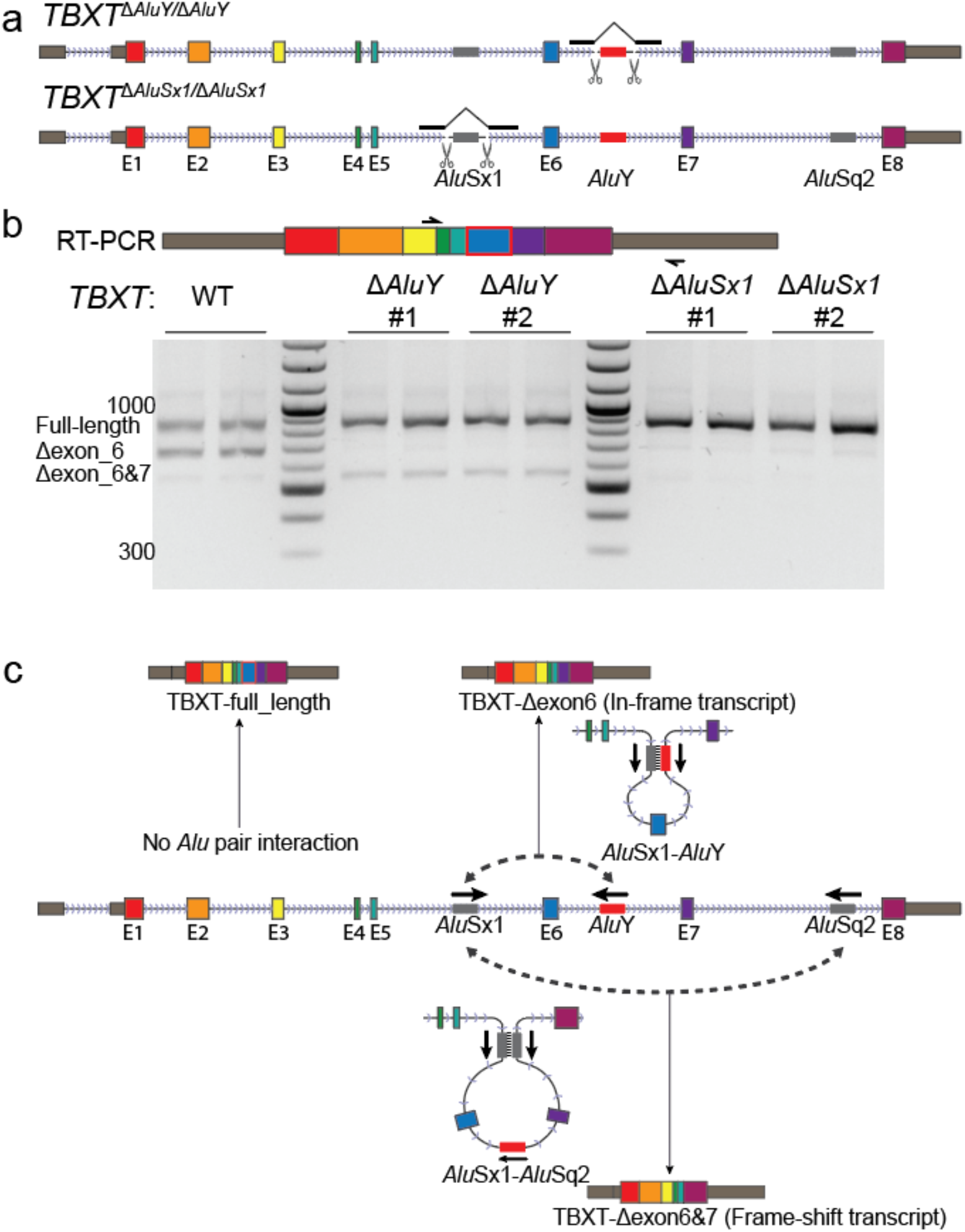
Both *Alu*Y and *Alu*Sx1 are required for *TBXT* alternative splicing. **a**, CRISPR-generated homozygous knock-outs of the *Alu*Y element in intron 6 and (in a separate line) *Alu*Sx1 element in intron 5 of *TBXT*. **b**, RT-PCR results of *TBXT* transcripts isolated from differentiated hESCs of wild-type, *ΔAluY*, and *ΔAluSx1* genotypes. Each genotype (*ΔAluY* and *ΔAluSx1*) was analyzed by two independent replicate clones. **c**, A schematic of *Alu* interactions and the corresponding *TBXT* transcripts in human, indicating that an *Alu*Y-*Alu*Sx1 interaction leads to the *TBXT-Δexon6* transcript. The *TBXT-Δexon6&7* transcript may stem from an *Alu*Sx1-*Alu*Sq2 interaction.

Interestingly, we found that wild-type differentiated hESCs also express a minor, previously un-annotated transcript that excludes both exon 6 and exon 7, leading to a frameshift and early truncation at the protein level (**Fig. 2b**, left, and **S4b**). Whereas deleting *Alu*Y slightly enhanced the abundance of this *TBXT-Δexon6&7* transcript, deleting *Alu*Sx1 in intron 5 completely eliminated this transcript (**Fig. 2b**). This may be best explained by a secondary interaction of the *Alu*Sx1 element with a distal *Alu*Sq2 element in intron 7. In this scenario, the secondary interaction would occur at a lower probability than the *Alu*Y-*Alu*Sx1 interaction pair (**Fig. 2c**, bottom). These results further support an interaction among intronic transposable elements affecting splicing of the conserved *TBXT* regulator (**Fig. 2c, S4b**).

### *Tbxt-Δexon6* is sufficient to induce tail loss in mice

To test whether the *TBXT-Δexon6* isoform is sufficient to induce tail loss, we generated a heterozygous mouse *Tbxt*^*Δexon6/+*^ model (**Fig. 3a**). *TBXT* is highly conserved in vertebrates and human and mouse protein sequences share 91% identity with a similar exon/intron architecture^11^. We could thus simulate a *TBXT-Δexon6* isoform by deleting exon 6 in mice, forcing splicing of exons 5 with exon 7. The *Tbxt*^*Δexon6/+*^ heterozygous mouse thus mimics the *TBXT* gene in human, which expresses both full-length and Δexon6 splice isoforms (**Fig. 2b and 3b-c**).

**Fig. 3.**
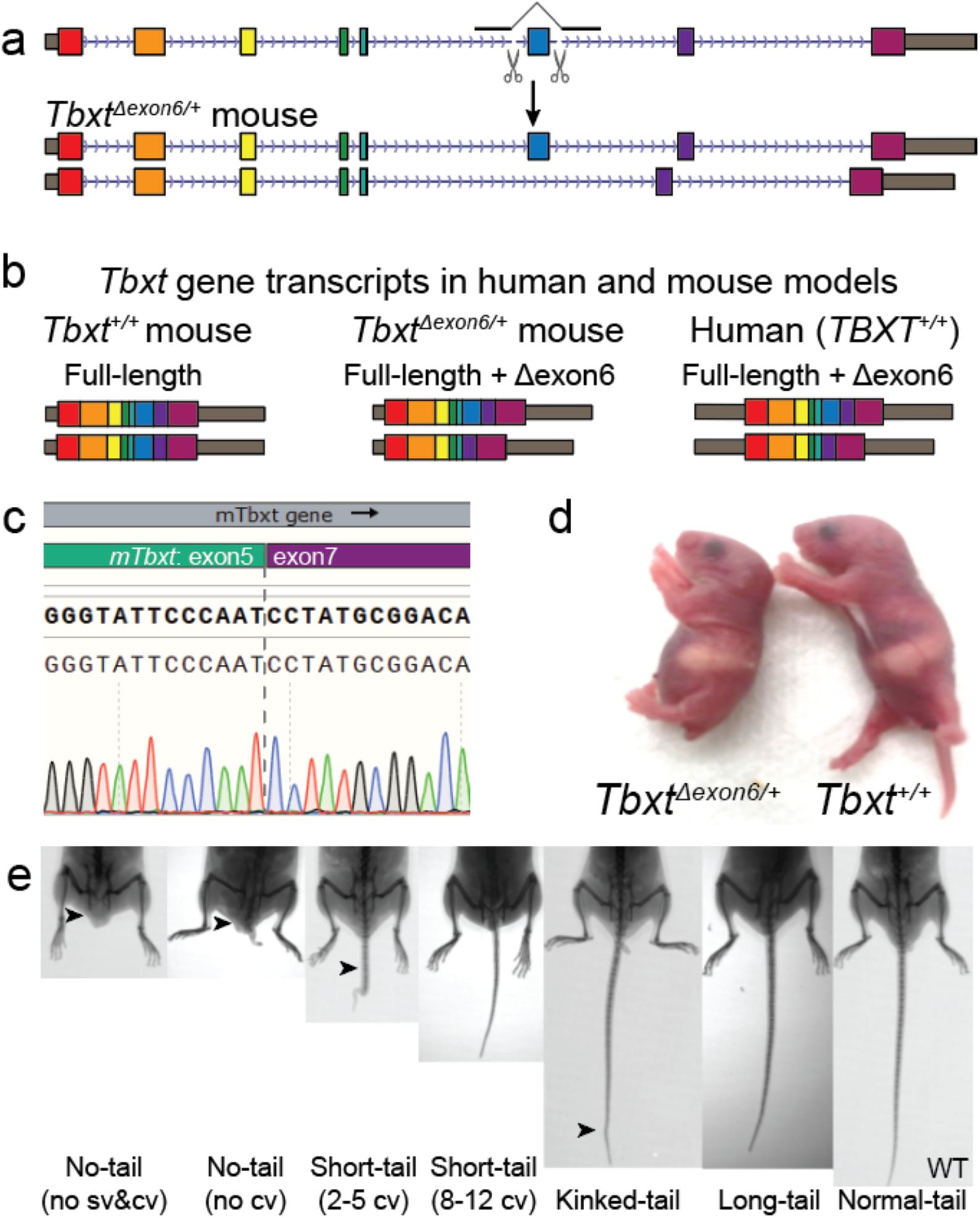
*TBXT-Δexon6* isoform is sufficient to induce tail-loss phenotype. **a**, CRISPR design for generating a *Tbxt*^*Δexon6/+*^ heterozygous mouse model. **b**, *Tbxt*^*Δexon6/+*^ mouse mimics *TBXT* gene expression products in human. **c**, Sanger-sequencing of the *Tbxt* RT-PCR product shows that deleting exon 6 in *Tbxt* leads to correct splicing by fusing exon 5 and 7. **d**, A representative *Tbxt*^*Δexon6/+*^ founder mouse (day 1) showing an absence of the tail. Two additional founder mice are shown in Figure S5. **e**, *Tbxt*^*Δexon6/+*^ heterozygous mice display heterogeneous tail phenotypes varied from absolute no-tail to long-tails. sv, sacral vertebrae; cv, caudal vertebrae.

Studying the phenotypes of the *Tbxt*^*Δexon6/+*^ mice, we found that simultaneous expression of both isoforms led to strong but heterogeneous tail morphologies, including no-tail and short-tail phenotypes (**Fig. 3d-e, S5**). Specifically, 21 of the 63 heterozygous mice showed tail phenotypes, while none of their 35 wild-type littermates showed phenotypes (**Table 1**). The incomplete penetration of phenotypes among the heterozygotes was stable across generations and founder lines: no-/short-tailed (*Tbxt*^*Δexon6/+*^) parent can give birth to long-tailed *Tbxt*^*Δexon6/+*^ mice, whereas long-tailed (*Tbxt*^*Δexon6/+*^) parents can give birth to pups with varied tail phenotypes (**Table 1, Fig. S5**), providing further evidence that the presence of *TBXT-Δexon6* suffices to induce tail loss.

**Table 1.** Genotype and phenotype analyses of the *Tbxt*^*Δexon6/+*^ intercrossed F2 pups.

| Genotypes | Total F2 pups | Pups with tail phenotypes | Tail phenotypes |  |  | Intercross (type 1)* | Intercross (type 2) |
| --- | --- | --- | --- | --- | --- | --- | --- |
|  |  |  | No-tail | Short-tail | Kinked-tail |  |  |
| *Tbxt*<sup>Δexon6/Δexon6</sup> | 0 | 0 | 0 | 0 | 0 | 0 | 0 |
| *Tbxt*<sup>Δexon6/+</sup> | 63 | 21 | 4 | 9 | 8 | 17(7)** | 46(14) |
| *Tbxt*<sup>+/+</sup> | 35 | 0 | 0 | 0 | 0 | 7(0) | 28(0) |

Note: \*: Type 1 intercrossing: at least one of the parent mice is no-/short-tailed. Type 2 intercrossing: both parent mice are long-tailed.
\*\* : Numbers in parentheses indicate the number of pups with tail phenotypes.

To control for the possibility that zygotic CRISPR targeting induced off-targeting DNA changes at the *Tbxt* locus, we performed Capture-seq covering the *Tbxt* locus and ∼200kb of both upstream and downstream flanking regions^37^ (**Fig. S6**). Capture-seq did not detect any off-targeting at the *Tbxt* locus across three independent founder mice, supporting our conclusion that the observed tail phenotype from the *Tbxt*^*Δexon6/+*^ mice derived from the *Tbxt-Δexon6* isoform.

### Homozygous removal of *Tbxt-Δexon6* is lethal

The human *TBXT* gene expresses a mixture of *TBXT-Δexon6* and *TBXT-full length* transcripts – induced as we inferred by the *Alu*Y insertion and interaction with *Alu*Sx1 – while mouse *Tbxt* only expresses the full length *Tbxt*. Thus, we next inquired into the mode by which homozygous *TBXT-Δexon6* mutation (*Tbxt*^*Δexon6*/*Δexon6*^) affects development. Intercrossing the *Tbxt*^*Δexon6/+*^ mice across multiple litters and replicated in different founders, we failed to produce viable homozygotes (**Table 1**). Dissecting intercrossed embryos at E11.5 showed that homozygotes either arrested development at ∼E9 or developed with spinal cord malformations that consequently led to death at birth (**Fig. S7**). We noted that the *Tbxt*^*Δexon6*/*Δexon6*^ embryos showed malformations of the spinal cord similar to spina bifida in humans. Together, while the *Tbxt*^*Δexon6/+*^ mice present incomplete penetrance of the tail phenotypes requires further investigation, these results indicate that the *TBXT-Δexon6* isoform, which in human is induced by the intronic *Alu*Y-*Alu*Sx1 interaction, may indeed be the key driver of tail-loss evolution in hominoids.

## Discussion

We have presented evidence that tail-loss evolution in hominoids was driven by the intronic insertion of an *Alu*Y element. As opposed to disrupting a splice site, we inferred that this element interacts with a neighboring (simian-shared) *Alu*Sx1 element in the neighboring intron, leading to an alternatively spliced isoform which skips an intervening exon (**Fig. 1c**). Experimental deletion of *Alu*Y or its interacting counterpart eliminates such *TBXT* alternative splicing in differentiated hESCs in the primitive streak state (**Fig. 2**).

Alternative splicing mediated by *Alu*-pairing in the *TBXT* gene demonstrates how an interaction between intronic transposable elements can dramatically modulate gene function to affect a complex trait. The human genome contains ∼1.8 million copies of SINE transposons – including ∼1 million *Alu*s – of which more than 60% are intronic^38^. Systematically searching for such interactions may lead to the identification of additional functional roles by which these elements impact human development and disease. Interestingly, inverted *Alu* pairs have been found to contribute to the biogenesis of exonic circular RNAs (circRNA)^39^ through ‘backsplicing’. Thus, it is an interesting possibility that the interactions between paired transposable elements may create functional splice variants and circRNA isoforms from the same genetic locus.

We found that expressing the *Tbxt-Δexon6* transcript – along with the full-length transcript – in mice was sufficient to induce no-tail phenotypes, though with incomplete penetrance (**Fig. 3** and **Table 1**). It is possible that a heterogeneity of tail phenotypes also existed in the ancestral hominoids upon the initial *Alu*Y insertion. Thus, while tail-loss evolution in hominoids may have been initiated by the *Alu*Y insertion, additional genetic changes may have then acted to stabilize the no-tail phenotype in early hominoids (**Fig. S8**). Such a set of genetic events would explain how a change to the *Alu*Y in modern hominoids would not result in the re-appearance of the tail.

The specific evolutionary advantage for the loss of the tail is not clear, though it likely involved enhanced locomotion in a non-arboreal lifestyle. We can assume however that the selective advantage must have been very strong since the loss of the tail may have included an evolutionary trade-off of neural tube defects, as demonstrated by the presence of spinal cord malformations in the *Tbxt*^*Δexon6*/*Δexon6*^ mutant at E11.5 (**Fig. S7**). Interestingly, mutations leading to neural tube defect and/or sacral agenesis have been detected in the coding and noncoding regions of the *TBXT* gene^40–43^. We thus speculate that the evolutionary tradeoff involving the loss of the tail – made ∼25 million years ago – continues to influence health today. This evolutionary insight into a complex human disease may in the future lead to the design of therapeutic strategies.

## Acknowledgments

We thank Naoya Yamaguchi, Eric Wang, John Shin, Susan Liao, Huiyuan Zhang, and the members of the Yanai and Boeke labs for constructive comments and suggestions. We thank Megan Hogan and Raven Luther for sequencing assistance, and Michael Ceriello and Ahmad Naimi for assistance with the mice work. This work was supported in part by the NHGRI RM1 HG009491 to J.B., and by the NYU Grossman School of Medicine with funding to I.Y. MTM is partially funded by NIH grant R35GM119703. B.X. was partially supported by the NYSTEM pre-doctoral fellowship (C322560GG).

## Author contributions

B.X. conceived the project. B.X., J.D.B. and I.Y designed the experiments with contribution from W.Z. and S.Y.K. B.X. led and conducted most of the experimental and analysis components, with contribution from W.Z., A.W., R.B. and M.P. E.H., R.B. and M.T.M. contributed to the capture-seq validation. A.M. and J.S.D. helped with embryo analysis work. J.D.B. and I.Y. supervised the study. B.X. drafted the manuscript. B.X., J.D.B. and I.Y. edited the manuscript with contribution from all authors.

## Methods

### Gene/protein sequence analysis

Protein sequences were downloaded from NCBI, and analyzed by the MUSCLE algorithm using MEGA X software with default settings^44^. Multiple species gene sequence alignment and analysis were done through the Ensembl Comparative Genomics (release 104) module^45^, using available species of hominoids (human, chimpanzee, bonobo, gorilla, orangutan and gibbon) and Old World monkeys (macaque, crab-eating macaque, pig-tailed macaque, olive baboon, drill, black snub-nosed monkey, golden snub-nosed monkey, Ma’s night monkey). The identified candidate gene (*TBXT*) was then visualized through the UCSC genome browser^36^ (Figure 1), highlighting the multiple sequence alignment mapping to the human genome (hg38) sequences.

### RNA secondary structure prediction

RNA secondary structure prediction of the *TBXT* intron5-exon6-intron6 sequence was performed using the ViennaRNA RNAfold Web Services (http://rna.tbi.univie.ac.at/). The algorithm calculates the folding probability using minimum free energy (MFE) matrix with default parameters. In addition, the calculation included the partition function and base pairing probability matrix.

### Human ESCs culture and differentiation

Human ESCs (WA01, also called H1, from WiCell Research Institute) were cultured with StemFlex Medium (Gibco, Cat. No. A3349401) in a feeder cell-free condition. Cells were grown on tissue culture-grade plates coated with hESC-qualified Geltrex (Gibco, Cat. No: A1413302). Geltrex was 1:100 diluted in DMEM/F-12 (Gibco, Cat. No. 11320033) supplemented with 1X GlutaMax (100X, Gibco, Cat. No. 35050061) and 1% Penicillin-Streptomycin (Gibco, Cat. No. 15070063). Before seeding hESCs, the plate was treated with Geltrex working solution in a tissue culture incubator (37°C and 5% CO_2_) for at least 1h.

Human ESC maintenance and culturing condition were performed according to the manufacturer’s protocol of StemFlex Medium. Briefly, StemFlex complete medium was made by combining StemFlex basal medium (450mL) with 50mL of StemFlex supplement (10X), plus 1% Penicillin-Streptomycin. Each Geltrex-coated well on a 6-well plate was seeded with 200K cells for ∼80% confluence in 3-4 days. Human ESCs were cryopreserved in PSC Cryomedium (Gibco, Cat. No. A2644601). The culturing medium was supplemented with 1X RevitaCell (100X, Gibco, Cat. No. A2644501, which is also included in the PSC Cryomedium kit) when cells had gone through stressed condition, such as freezing-and-thawing or nucleofection. RevitaCell supplemented medium was replaced with regular StemFlex complete medium on the second day. hESCs grown under RevitaCell condition might become stretched, but would recover after replacing to the normal StemFlex complete medium.

The human ESC differentiation assay to induce a gene expression pattern of primitive streak was adapted from Xi *et al*^34^. On day -1, freshly cultured hESC colonies were dissociated into clumps (2-5 cells) with Versene buffer (with EDTA, Gibco, Cat. No. 15040066). The dissociated cells were seeded on Geltrex-coated 6-well tissue culture plates at 25,000 cells/cm^2^ (0.25M per well in the 6-well plates) in StemFlex complete medium. Differentiation to the primitive streak state was initiated on the next day (day 0) by switching StemFlex complete medium to basal differentiation medium. Basal differentiation medium (50mL) was made with 48.5mL DMEM/F-12, 1% GlutaMax (500uL), 1% ITS-G (500uL, Gibco Cat. No. 41400045), and 1% penicillin-streptomycin (500 µL), and supplemented with 3µM GSK3 inhibitor CHIR99021 (10µL of 15mM stock solution in DMSO. Tocris, Cat. No. 4423). The cells were collected at differentiation day 1 to 3 for downstream experiments, which confirmed the expression fluctuations of mesoderm genes (*TBXT* and *MIXL1*) in a 3-day differentiation period (Fig. S3)^34^.

### Mouse ESC culture and differentiation

Mouse ESCs derived from C57BL/6J strain background were cultured in a feeder cell-free condition. Cells were grown on tissue culture-grade plates coated with mESC-qualified gelatin. Before plating cells, the plastic tissue culture-treated plates were coated with 0.1% gelatin (EMD Millipore ES-006-B) at room temperature for at least 30min, followed by switching to mESC medium and warming up the medium at 37°C and 5% CO2 incubator for at least 30min.

The feeder cell-free mESC culturing medium, also called ‘80/20’ medium, comprises 80% 2i medium and 20% mESC medium by volume. 2i medium was made from a 1:1 mix of Advanced DMEM/F-12 (Gibco, Cat. No. 12634010) and Neurobasal-A (Gibco, Cat. No. 10888022), 1X N-2 supplement (Gibco, Cat. No. 17502048), 1X B-27 supplement (Gibco, Cat. No. 17504044), 1X Glutamax (Gibco, Cat. No. 35050061), 0.1 mM Beta-Mercaptoethanol (Gibco, Cat. No. 31350010), 1000 units/mL LIF (Millipore, Cat. No. ESG1107), 1 µM MEK1/2 inhibitor (Stemgent, Cat. No. PD0325901), and 3 µM GSK3 inhibitor CHIR99021 (Tocris, Cat. No. 4423). mESC medium was made from Knockout DMEM (Gibco, Cat. No. 10829018), containing 15% Fetal Bovine Serum (GeminiBio, Cat. No. 100-106), 0.1 mM Beta-Mercaptoethanol, 1X MEM Non Essential Amino Acids (Gibco, Cat. No. 11140050), 1X Glutamax, 1X Nucleosides (Millipore, Cat. No. ES-008-D) and 1000 units/mL LIF.

mESC differentiation for inducing *Tbxt* gene expression was adapted from Pour *et al* in a feeder cell-free condition^46^. Cells were first plated in 80/20 medium for 24 hours on a gelatin-coated 6-well plate, followed by switching to N2/B27 medium without LIF or 2i for another 2-day culturing. The N2/B27 medium (50mL) is made with 18mL Advanced DMEM/F-12, 18mL Neurobasal-A, 9mL Knockout DMEM, 2.5mL Knockout Serum Replacement (Gibco, Cat. No. 10828028), 0.5mL N-2 supplement, 1mL B-27 supplement, 0.5mL Glutamax (100X), 0.5mL Nucleosides (100X), and 0.1 mM Beta-Mercaptoethanol. Then the N2/B27 medium was supplemented with 3µM GSK3 inhibitor CHIR99021 for induced differentiation (day 0). The cells were collected at differentiation day 1 to 3 for downstream experiments, which showed consistent results of *Tbxt* gene expression fluctuations in a 3-day differentiation period.

### CRISPR targeting

All guide RNAs of the CRISPR experiments were designed using CRISPOR algorithm through its predicted target sites integrated in the UCSC genome browser^47^. Guide RNAs were cloned into the pX459V2.0-HypaCas9 plasmid (AddGene plasmid #62988), or its custom derivative by replacing the puromycin resistance gene to blasticidin resistance gene. Guide RNAs in this study were designed in pairs to delete the intervening sequences. The sequence and targeting sites of the guide RNAs were listed below:

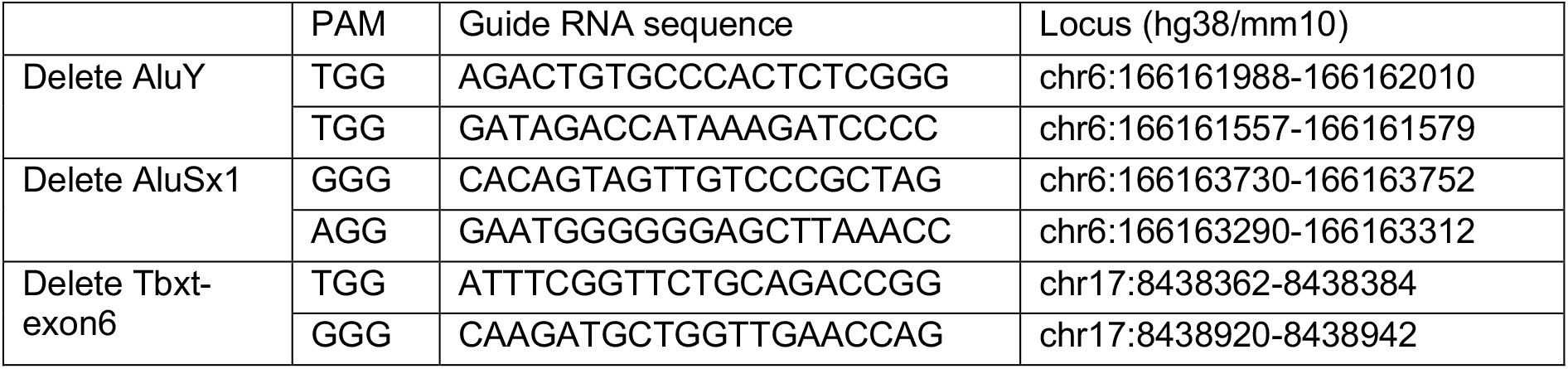

All oligos (plus Goldern-Gate assembly overhangs) were synthesized from Integrated DNA Technologies (IDT) and ligated into empty pX459V2.0 vector following standard Golden Gate Assembly protocol using BbsI restriction enzyme (NEB, Cat. No. R3539). The constructed plasmids were purified from 3mL *E. coli* cultures using ZR Plasmid MiniPrep Purification Kit (Zymo Research, Cat. No. D4015) for sequence verification. Plasmids delivered into ESCs were purified from 250mL *E. coli* cultures using PureLink HiPure Plasmid Midiprep Kit (Invitrogen, Cat. No. K210005). To facilitate DNA delivery to ESCs through nucleofection, the purified plasmids were resolved in Tris-EDTA buffer (pH 7.5) for a concentration of at least 1 µg/µL in a sterile hood.

### DNA delivery

DNA delivery into human/mouse ESCs for CRISPR/Cas9 targeting were performed using a Nucleofector 2b Device (Lonza, Cat. No. BioAAB-1001). Human Stem Cell Nucleofector Kit 1 (Cat. No. VPH-5012) and mESC Nucleofector Kit (Lonza, Cat. No. VVPH-1001) were used for delivering DNA into human and mouse ESCs, respectively. ESCs were double-feeded the day before the nucleofection experiment to maintain a superior condition.

Before performing nucleofection on human ESCs, 6cm tissue culture plates were treated with 0.5µg/cm^2^ rLaminin-521 in a 37°C and 5% CO_2_ incubator for at least 2h. rLaminin-521-treated plates give better viability when seeding hESCs as single cells. Cultured human ESCs were then washed with PBS, and dissociated into single cells using TrypLE Select Enzyme (no phenol red. Gibco, Cat. No. 12563011). One million hES single cells were nucleofected using program A-023 according to the manufacturer’s instruction of the Nucleofector 2b device. Transfected cells were transferred on the rLaminin-521-treated 6cm plates with pre-warmed StemFlex complete medium supplementing with 1X RevitaCell but not Penicillin-Streptomycin. Antibiotic selection was performed 24h after nucleofection with puromycin (Gibco, Cat. No. A1113802).

Mouse ESCs were dissociated into single cells using StemPro Accutase (Gibco, Cat. No. A1110501) and five million cells were transfected using program A-023 according to the manufacturer’s instruction. Cells were plated onto gelatin-treated 10cm plates, followed by antibiotic selection 24h after nucleofection with blasticidin (Gibco, Cat. No. A1113903).

Together with the pX459V2.0-HypaCas9-gRNA plasmids for nucleofection, a single-strand DNA oligo were co-delivered for micro homology-induced deletion of the targeted sites^48^. These ssDNA sequences were synthesized from IDT through its Ultramer DNA Oligo service, including phosphorothioate bond modification on the three bases of each end. Detailed sequence information was listed below (“|” indicates a junction site):

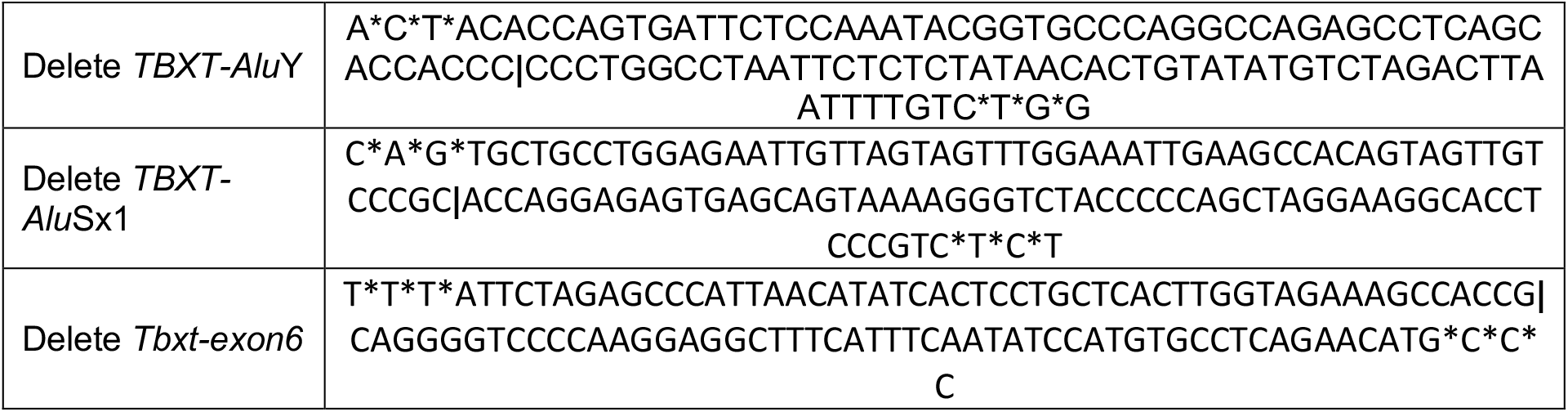

Genotyping of CRISPR/Cas9-targeted sites were performed through PCR following standard protocol. The genotyping primers were listed below:

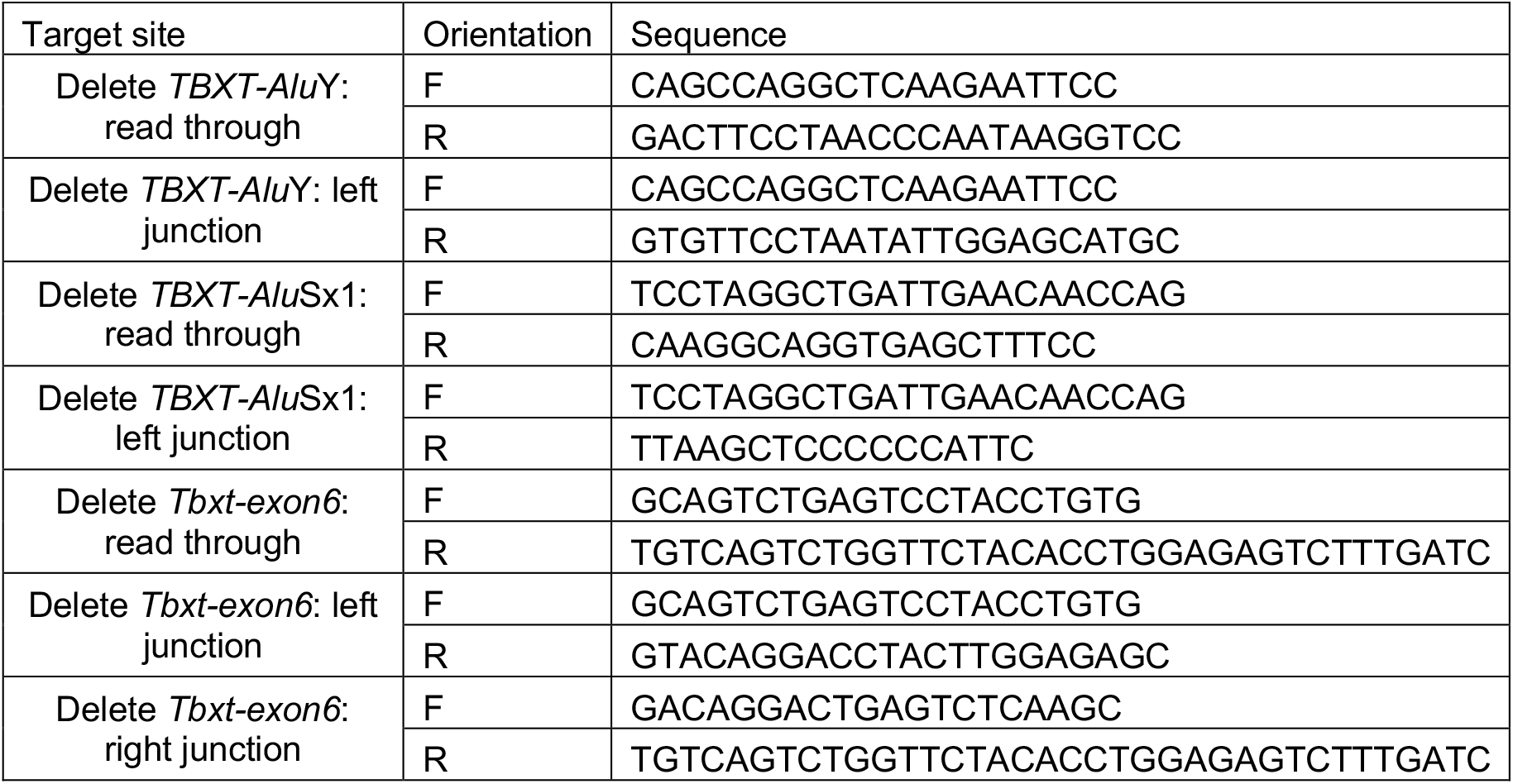

### Splicing isoforms detection

Total RNAa were collected from the undifferentiated or differentiated cells of both human and mouse ESCs, using standard column-based purification kit (QIAGEN RNeasy Kit, Cat. No. 74004). DNase treatment was applied during the purification to remove any potential DNA contamination. Following extraction, RNA quality was checked through electrophoresis based on the ribosomal RNA integrity. Reverse transcription was performed with 1µg of high-quality total RNA for each sample, using High-Capacity RNA-to-cDNA™ Kit (Applied Biosystems, Cat. No. 4387406). DNA oligos used for PCR/RT-PCR/RT-qPCR were listed below:

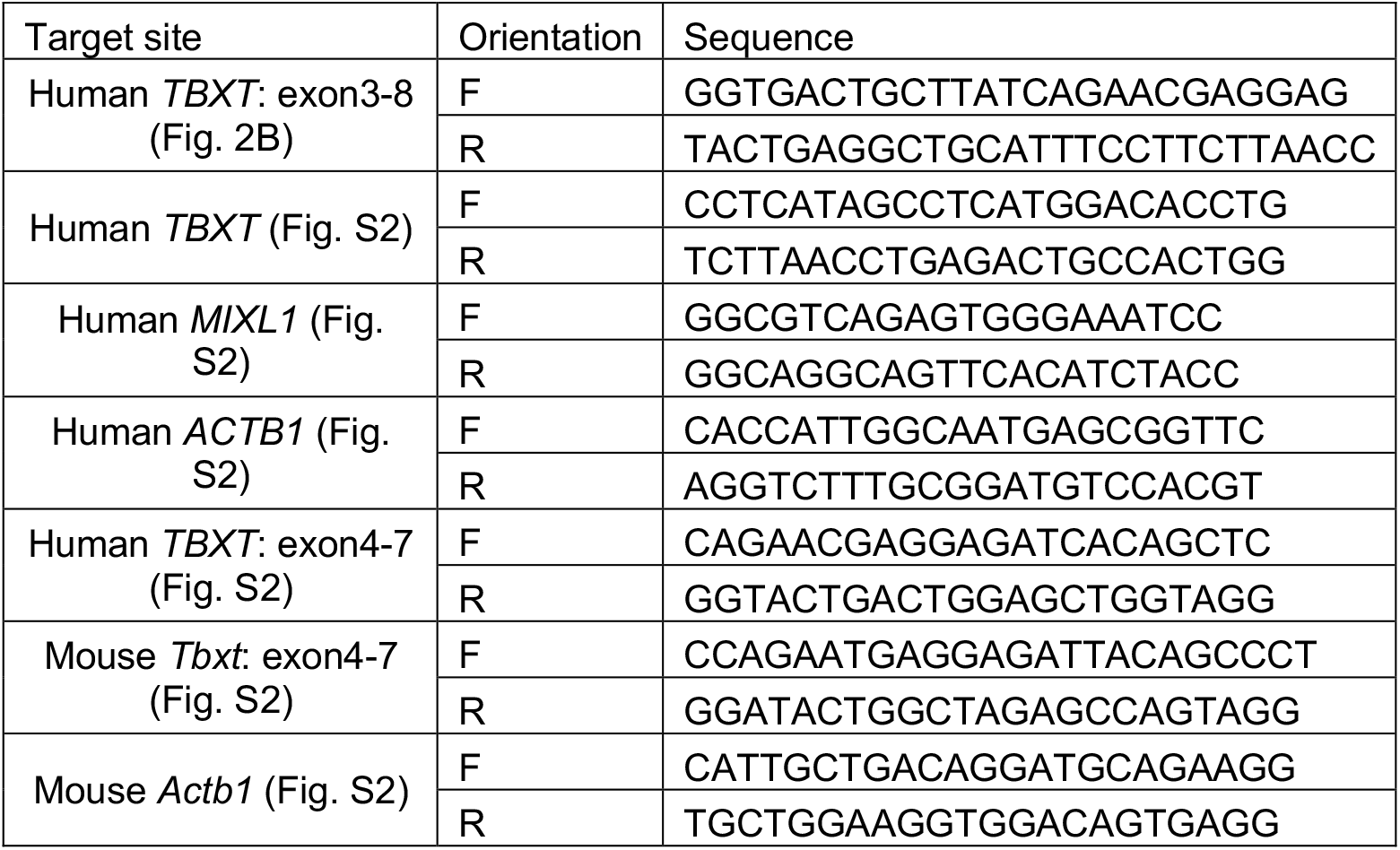

### Mouse work

All mouse work was done following NYULH’s animal protocol guidelines. The *Tbxt*^*Δexon6/+*^ heterozygous mouse model was generated through zygotic microinjection, using an experimental protocol adapted from Yang *et al*^49^. Briefly, Cas9 mRNA (MilliporeSigma, Cat. No. CAS9MRNA), synthetic guide RNAs, and single-stranded DNA oligo were co-injected into the 1-cell stage zygotes following the described procedures^49^. Synthetic guide RNAs were ordered from Synthego as their custom CRISPRevolution sgRNA EZ Kit, with the same targeting sites as used in the CRISPR deletion experiment of mouse ESCs (AUUUCGGUUCUGCAGACCGG and CAAGAUGCUGGUUGAACCAG). The co-injected single-stranded DNA oligo is the same as above mentioned as well. Processed embryos were then *in vitro* cultured to the blastomeric stage, followed by embryo transferring to the pseudopregnant foster mothers. Following zygotic microinjection and transferring, founder pups were screened based on their abnormal tail phenotypes. DNA samples were collected through ear punches at day ∼21 for genotyping.

Upon confirming the heterozygous genotype (*Tbxt*^*Δexon6/+*^), founder mice were backcrossed with wild-type C57B/6J mice for generating heterozygous F1 pups. Due to the varied tail phenotypes, intercrossing between F1 heterozygotes were performed in two categories: type1 intercrossing includes at least one parent being no-/short-tailed, whereas type 2 intercrossing were mated between two long-tailed F1 heterozygotes. Both types of intercrossing produced heterogeneous tail phenotypes in F2 *Tbxt*^*Δexon6/+*^ pups, confirming the incomplete penetrance of tail phenotypes, and the absence of homozygotes (*Tbxt*^*Δexon6/Δexon6*^), as summarized in Table 1. To confirm the embryonic phenotypes in homozygotes, embryos were dissected at E11.5 gestation stage from the timed pregnant mice through the standard protocol. Adult mice (>12 weeks) were anesthetized for X-ray imaging of vertebra using a Bruker In-Vivo Xtreme IVIS imaging system.

### Capture-seq genotyping

Capture-seq, or targeted sequencing of the loci of interest, was performed as previously described^37^. Conceptually, capture-seq uses custom biotinylated probes to pull down the genomic loci of interest from the standard whole-genome sequencing libraries, thus enabling sequencing of the specific genomic loci in a much higher depth while reducing the cost.

Genomic DNA were purified from mESCs or ear punches of founder mice using Zymo Quick-DNA Miniprep Plus Kit (Cat. No. D4068) according to manufacturer’s instruction. DNA sequencing libraries compatible for Illumina sequencers were prepared following standard protocol. Briefly, 1µg of gDNA was sheared to 500-900 base pairs in a 96-well microplate using the Covaris LE220 (450 W, 10% Duty Factor, 200 cycles per burst, and 90-s treatment time), followed by purification with a DNA Clean and Concentrate-5 Kit (Zymo Research, Cat. No. D4013). Sheared and purified DNA were then treated with end repair enzyme mix (T4 DNA polymerase, Klenow DNA polymerase, and T4 polynucleotide kinase, NEB, Cat. No. M0203, M0210 and M0201, respectively), and A-tailed using Klenow 3’-5’exo-enzyme (NEB, Cat. No. M0212). Illumina sequencing library adapters were subsequently ligated to DNA ends, followed by PCR amplification with KAPA 2X Hi-Fi Hotstart Readymix (Roche, Cat. No. KR0370).

Custom biotinylated probes were prepared as bait through nick translation, using BAC DNA and/or plasmids as the template. The probes were prepared to comprehensively cover the whole locus. We used BAC lines RP24-88H3 and RP23-159G7, purchased from BACPAC Genomics, to generate bait probes covering mouse *Tbxt* locus and ∼200kb flanking sequences in both upstream and downstream regions. The pooled whole-genome sequencing libraries were hybridized with the biotinylated baits in solution, and purified through streptavidin-coated magnetic beads. Following pull-down, DNA sequencing libraries were quantified with Qubit 3.0 Fluorometer (Invitorgen, Cat. No. Q33216) using a dsDNA HS Assay Kit (Invitorgen, Cat. No. Q32851). The sequencing libraries were subsequently sequenced on an Illumina NextSeq 500 sequencer in paired-end mode.

Sequencing results were demultiplexed with Illumina bcl2fastq v2.20 requiring a perfect match to indexing BC sequences. Low quality reads/bases and Illumina adapters were trimmed with Trimmomatic v0.39. Reads were then mapped to mouse genome (mm10) using bwa 0.7.17. The coverage and mutations in and around *Tbxt* locus were checked through visualization in a mirror version of UCSC genome browser.

## Supplementary Figures

**Fig. S1.**
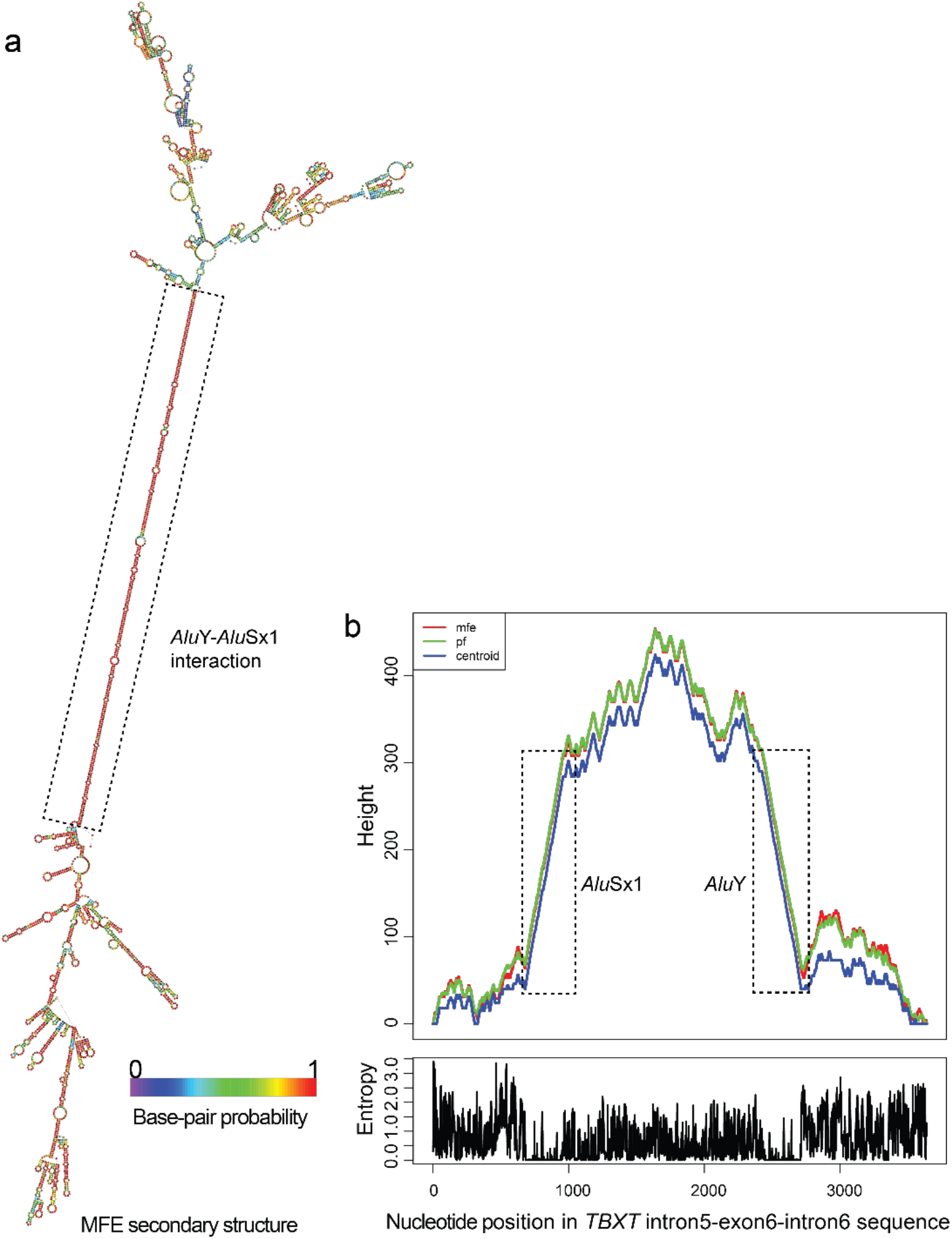
RNA structure prediction using RNAfold algorithm of the ViennaRNA package^33^. **a**, Predicted RNA secondary structure of the *TBXT* intron5-exon6-intron6 sequence. The paired *Alu*Y-*Alu*Sx1 region is highlighted. **b**, Mountain plot of the RNA secondary structure prediction, showing the ‘height’ in predicted secondary structure across the nucleotide positions. Height is computed as the number of base pairs enclosing the base at a given position. Overall, the *Alu*Sx1 and *Alu*Y regions are predicted to form helices with high probability (low entropy).

**Fig. S2.**
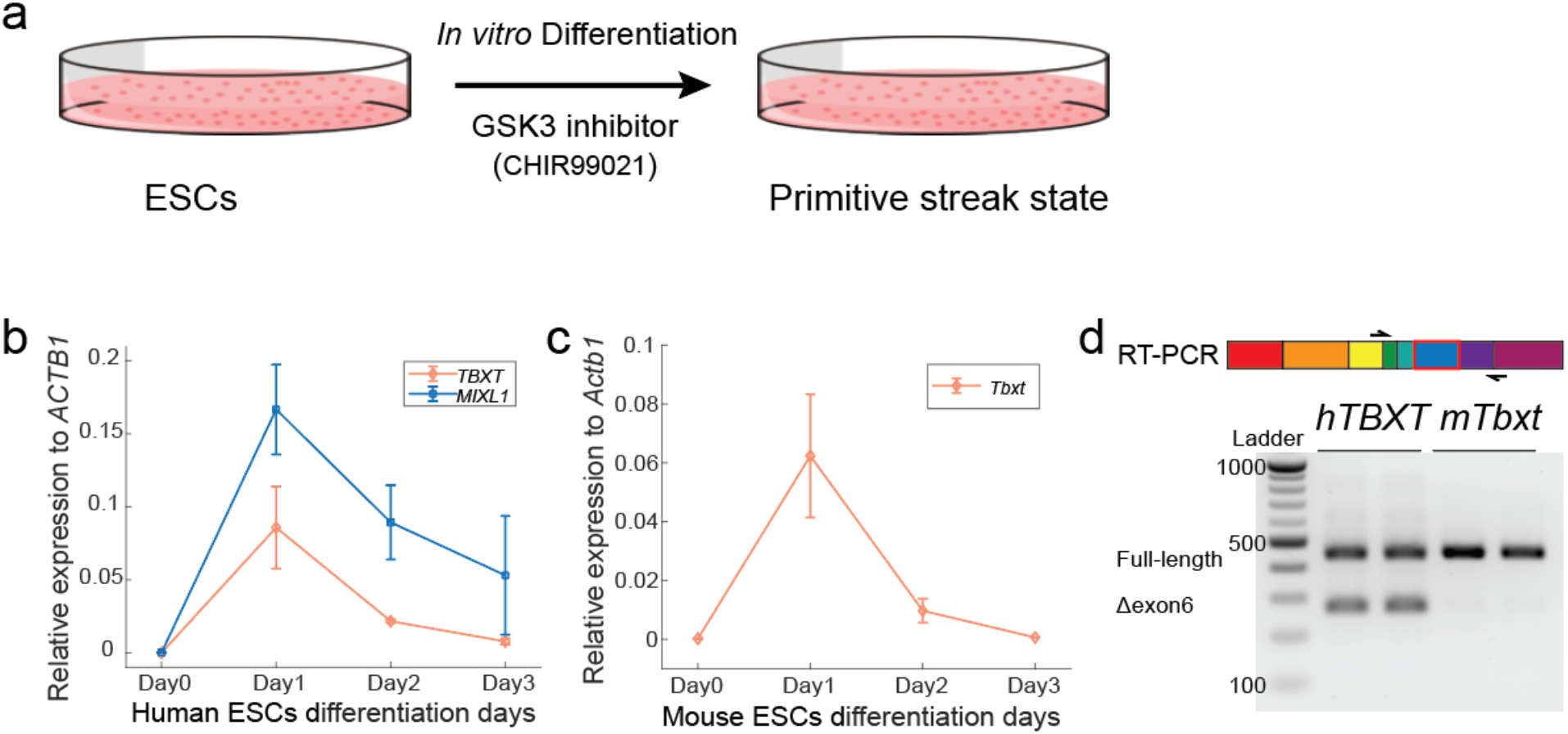
Studying *TBXT* expression isoforms using primitive streak *in vitro* differentiation. **a**, Human and mouse ESCs *in vitro* differentiation for inducing *TBXT* expression. Human and mouse ESCs differentiation assay was adapted from Xi *et al*^34^ and Pour *et al*^35^, respectively. **b**, Quantitative RT-PCR of *TBXT* and *MIXL1* expression during hESC differentiation, indicating correct induction of mesodermal gene expression program^34^. **c**, Quantitative RT-PCR of *Tbxt* expression during mESC differentiation. **d**, RT-PCR of *TBXT*/*Tbxt* transcripts in human and mouse, highlighting a unique *Δexon6* splicing isoform in human.

**Fig. S3.**
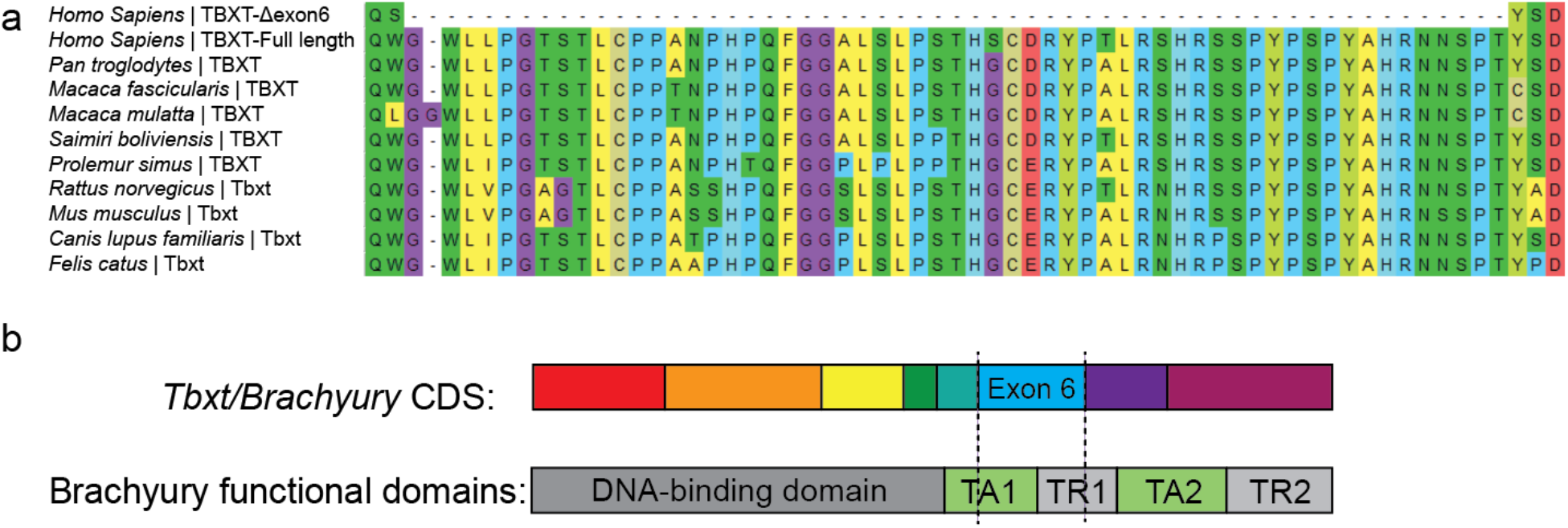
*TBXT* exon 6 conservation and functional domain analysis. **a**, Protein sequence alignment of the *TBXT* exon 6 region in representative mammals. With the exception of humans and chimpanzees, all are tailed. **b**, The exon 6-derived peptide of TBXT overlaps with large fractions of transcription regulation domains. TA, transcription activation; TR, transcriptional repression. Functional domain annotation of Brachyury was adapted from Kispert *et al*^12^.

**Fig. S4.**
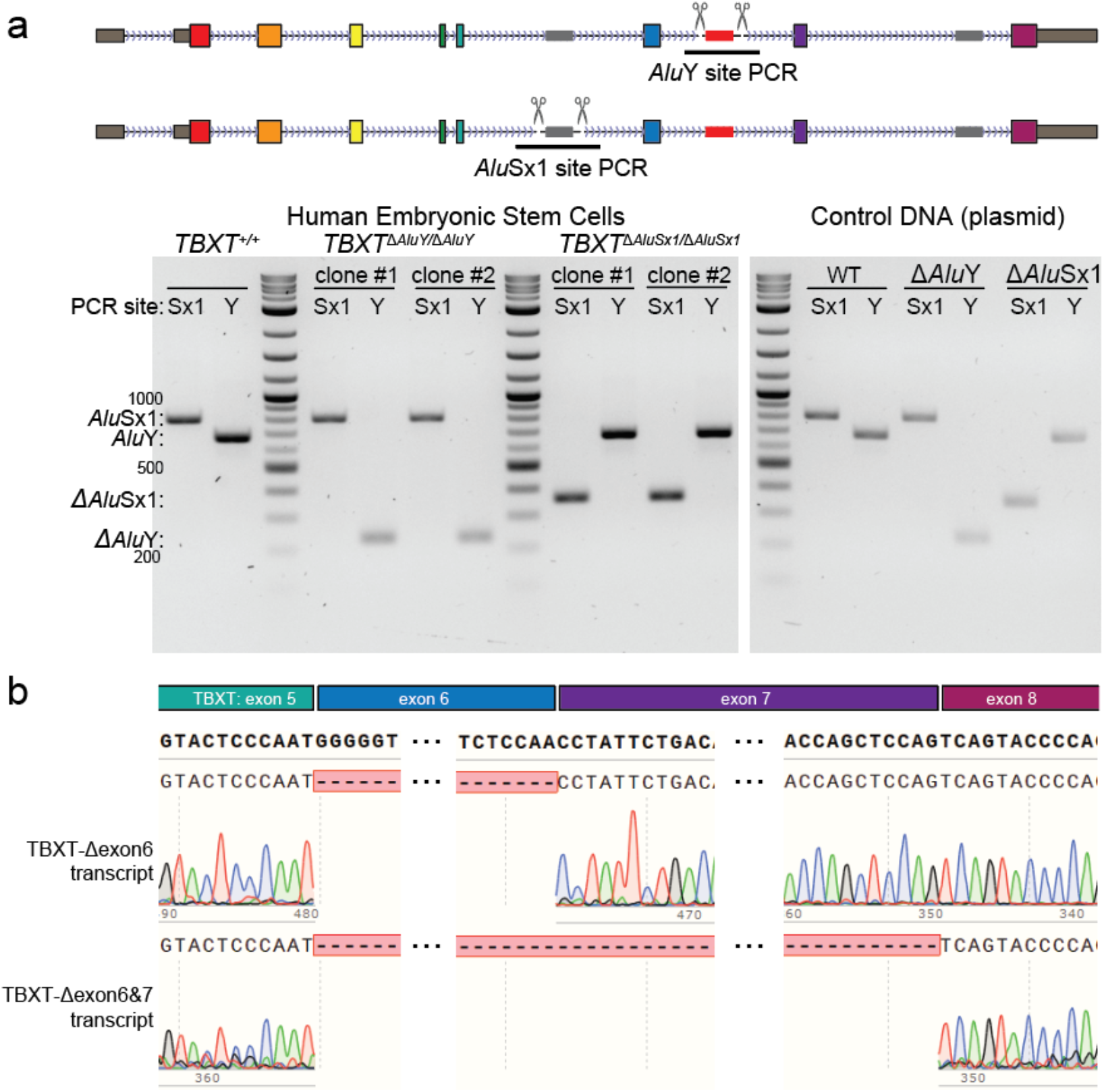
Validation of hESC CRISPR-deletion clones and the *TBXT* expression isoforms. **a**, PCR validation of the hESC clones with deletions of *Alu*Y or *Alu*Sx1 in *TBXT*. PCR validation for each clone or control samples were performed in pairs, each amplifying the *Alu*Sx1 locus (Sx1) or the *Alu*Y locus (Y), respectively, with primers that bind the two flanking sequences of the deleted region. Each genotype included two independent clones of *Alu*Y deletion or *Alu*Sx1 deletion, corresponding to the two replicates in Figure 2B. **b**, Sanger sequencing of the *TBXT*-Δexon6 and *TBXT*-Δexon6&7 transcripts detected in Figure 2B. The sequencing results were aligned to the full length *TBXT* mRNA sequence.

**Fig. S5.**
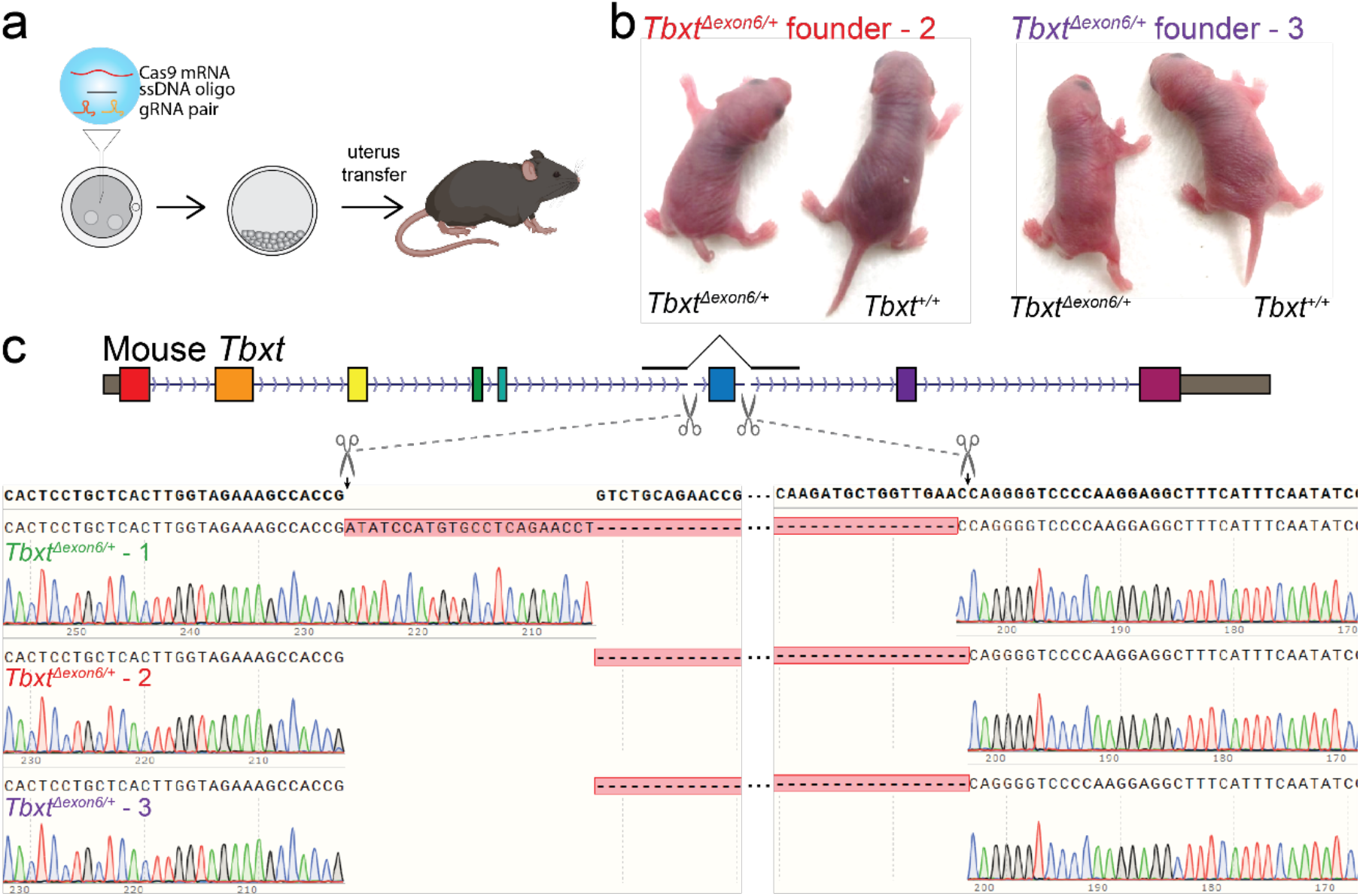
*Tbxt*^*Δexon6/+*^ founder mice generated through CRISPR/Cas9 targeting in the zygotes. **a**, Schematic of zygotic injection of CRISPR/Cas9 reactions. **b**, Two *Tbxt*^*Δexon6/+*^ founder mice (in addition to the one shown in Fig. 3) indicating an absence or reduced form of the tail (Founders 2 & 3). **c**, Sanger sequencing of the exon 6-deleted allele isolated from the genomic DNA of *Tbxt*^*Δexon6/+*^ founder mice. Founder 1 had an unexpected insertion of 23 base pairs at the CRISPR cutting site in the original intron 5 of *Tbxt*. Both founder 2 and 3 had the exact fusion between the two CRISPR cutting sites in introns 5 and 6.

**Fig. S6.**
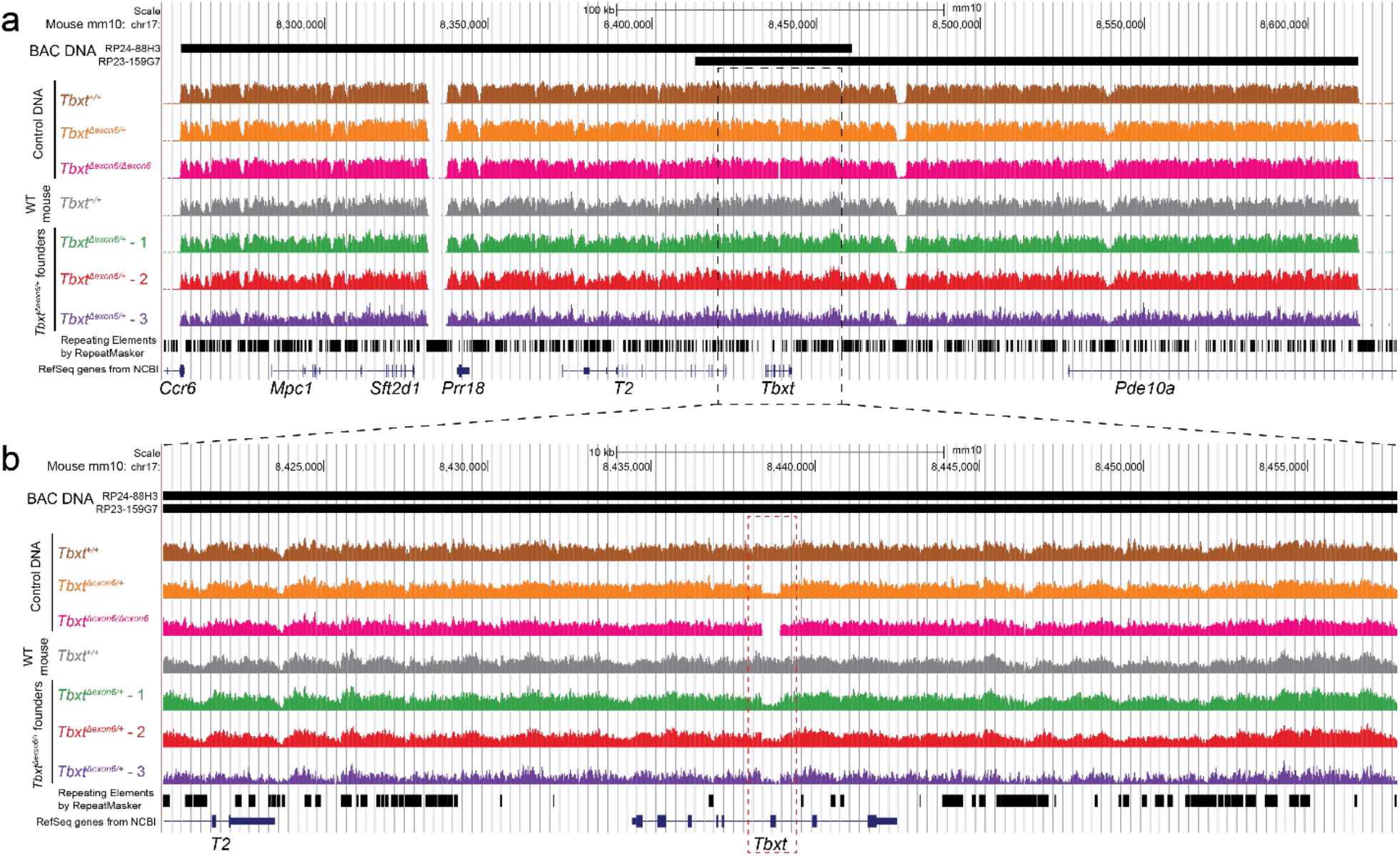
Capture-seq at the *Tbxt* locus of founder mice did not detect off target mutations. **a**, Capture-seq of the founder mice using baits generated from bacterial artificial chromosomes (RP24-88H3 and RP23-159G7). The shallow-covered regions are typically repeat sequences in the mouse genome and are consistent across samples. Control DNAs were obtained from wild-type or exon6-deleted mESCs through CRISPR targeting. **b**, A zoom-in view of the Capture-seq results at the *Tbxt* locus, highlighting the CRISPR-deleted exon 6 region.

**Fig. S7.**
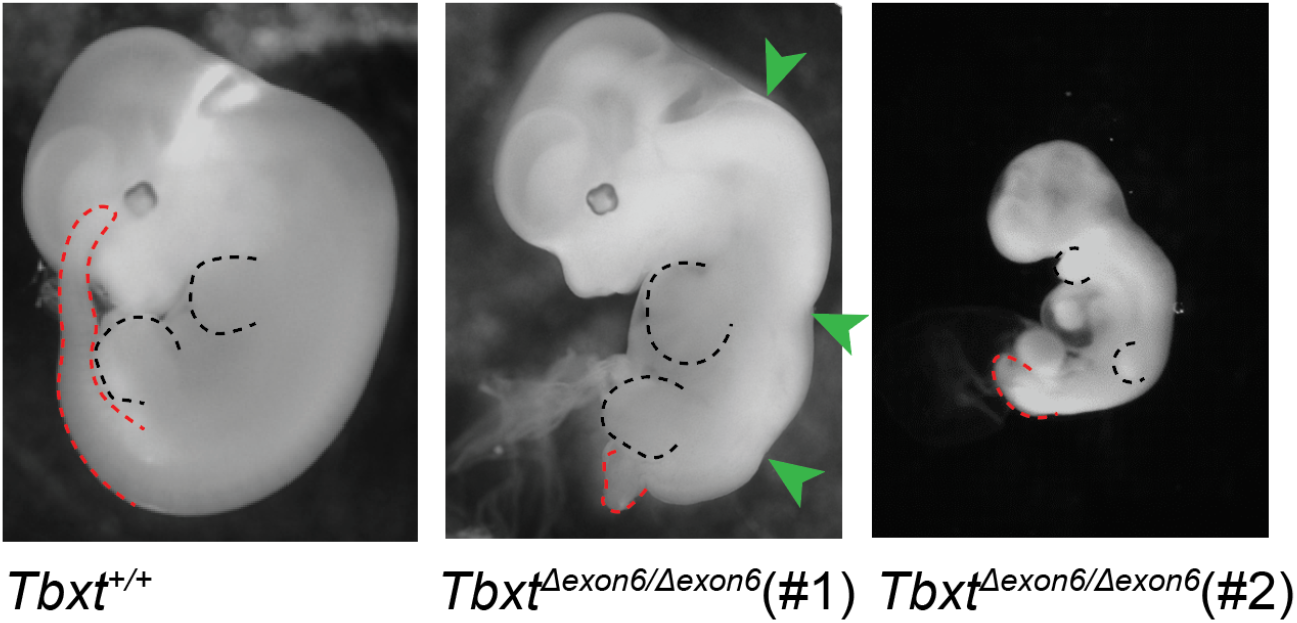
Analysis of *Tbxt*^*Δexon6*/*Δexon6*^ embryos at the E11.5 stage. *Tbxt*^*Δexon6*/*Δexon6*^ embryos either develop spinal cord defects (middle) that die at birth or arrest at approximately stage E9 of development (right). Red and black dashed lines mark the embryonic tail regions and limb buds, respectively. Green arrowheads in the middle panel indicate malformed spinal cord regions.

**Fig. S8.**
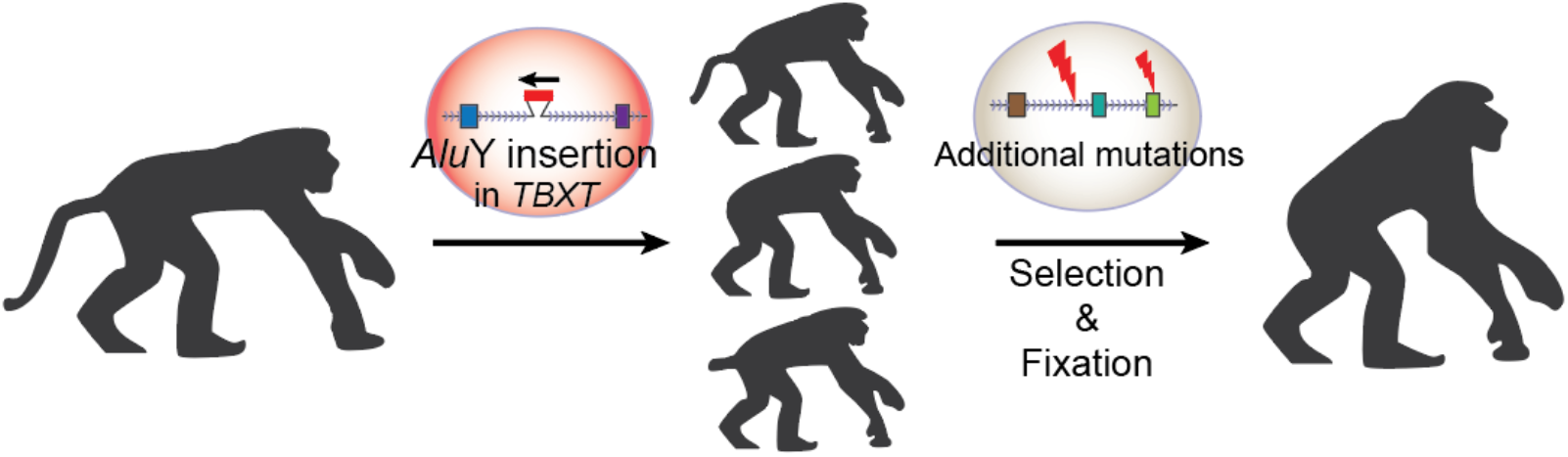
A model for tail-loss evolution in the early hominoids. The *Alu*Y insertion in *TBXT* marked an early genetic event that initiated tail-loss evolution in the hominoid common ancestor. Additional genetic changes may have then acted to stabilize the no-tail phenotype in the ancient hominoids.

