## Supplementary File for "The genetic basis of tail-loss evolution in humans and apes"

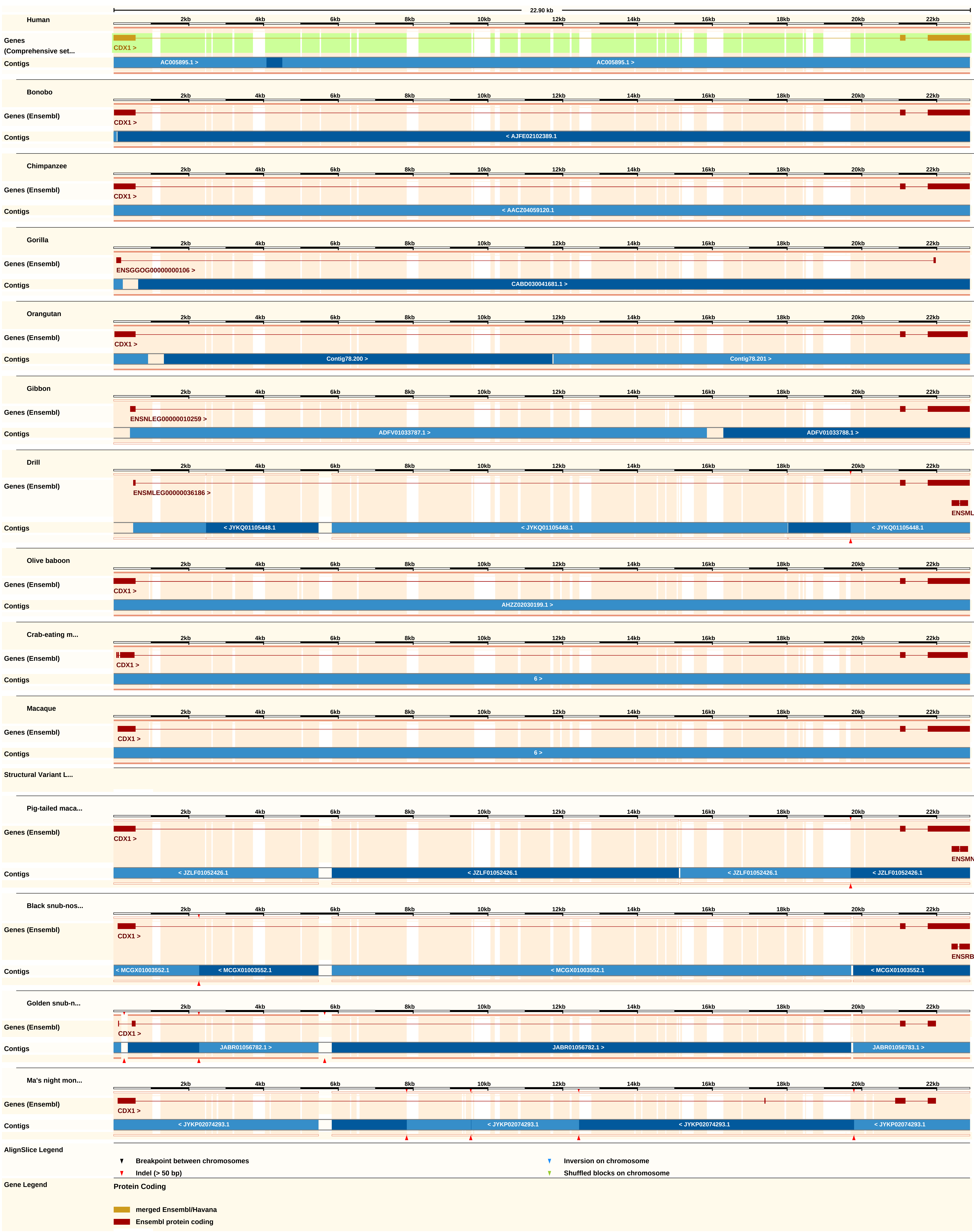
22.90 kb
Human
Genes (Comprehensive set...
CDX1 >
Contigs
AC005895.1 >
Bonobo
Genes (Ensembl)
< AJFE02102389.1
Chimpanzee
< AACZ04059120.1
Gorilla
ENSGGOG00000000106 >
CABD030041681.1 >
Orangutan
Contig78.200 >
Contig78.201 >
Gibbon
ENSNLEG00000010259 >
ADFV01033787.1 >
ADFV01033788.1 >
Drill
ENSMLEG00000036186 >
< JYKQ01105448.1
Olive baboon
AHZZ02030199.1 >
Crab-eating m...
6 >
Macaque
Structural Variant L...
Pig-tailed maca...
< JZLF01052426.1
Black snub-nos...
< MCGX01003552.1
Golden snub-n...
JABR01056782.1 >
JABR01056783.1 >
Ma's night mon...
< JYKP02074293.1
AlignSlice Legend
Breakpoint between chromosomes
Indel (> 50 bp)
Inversion on chromosome
Shuffled blocks on chromosome
Gene Legend
Protein Coding
merged Ensembl/Havana
Ensembl protein coding

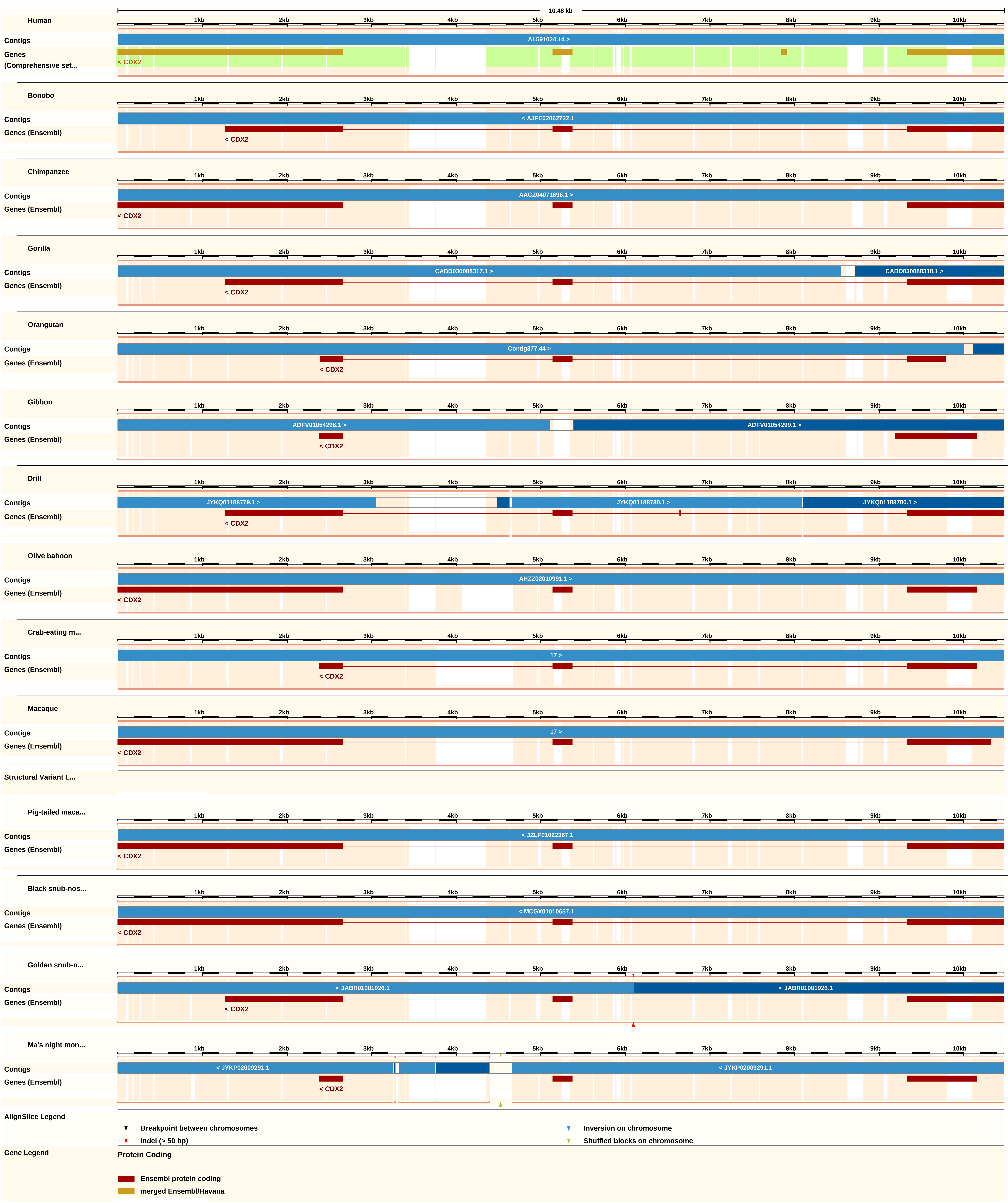
10.48 kb
1kb 2kb 3kb 4kb 5kb 6kb 7kb 8kb 9kb 10kb
Human
Contigs
AL591024.14 >
Genes (Comprehensive set...
< CDX2
Bonobo
Contigs
< AJFE02062722.1
Genes (Ensembl)
< CDX2
Chimpanzee
Contigs
AACZ04071696.1 >
Genes (Ensembl)
< CDX2
Gorilla
Contigs
CABD030088317.1 >
CABD030088318.1 >
Genes (Ensembl)
< CDX2
Orangutan
Contigs
Contig377.44 >
Genes (Ensembl)
< CDX2
Gibbon
Contigs
ADFV01054298.1 >
ADFV01054299.1 >
Genes (Ensembl)
< CDX2
Drill
Contigs
JYKQ01188779.1 >
JYKQ01188780.1 >
JYKQ01188780.1 >
Genes (Ensembl)
< CDX2
Olive baboon
Contigs
AHZZ02010991.1 >
Genes (Ensembl)
< CDX2
Crab-eating m...
Contigs
17 >
Genes (Ensembl)
< CDX2
Macaque
Contigs
17 >
Genes (Ensembl)
< CDX2
Structural Variant L...
Pig-tailed maca...
Contigs
< JZLF01022367.1
Genes (Ensembl)
< CDX2
Black snub-nos...
Contigs
< MCGX01010657.1
Genes (Ensembl)
< CDX2
Golden snub-n...
Contigs
< JABR01001926.1
< JABR01001926.1
Genes (Ensembl)
< CDX2
Ma's night mon...
Contigs
< JYKP02009291.1
< JYKP02009291.1
Genes (Ensembl)
< CDX2
AlignSlice Legend
Breakpoint between chromosomes
Inversion on chromosome
Indel (> 50 bp)
Shuffled blocks on chromosome
Gene Legend
Protein Coding
Ensembl protein coding
merged Ensembl/Havana

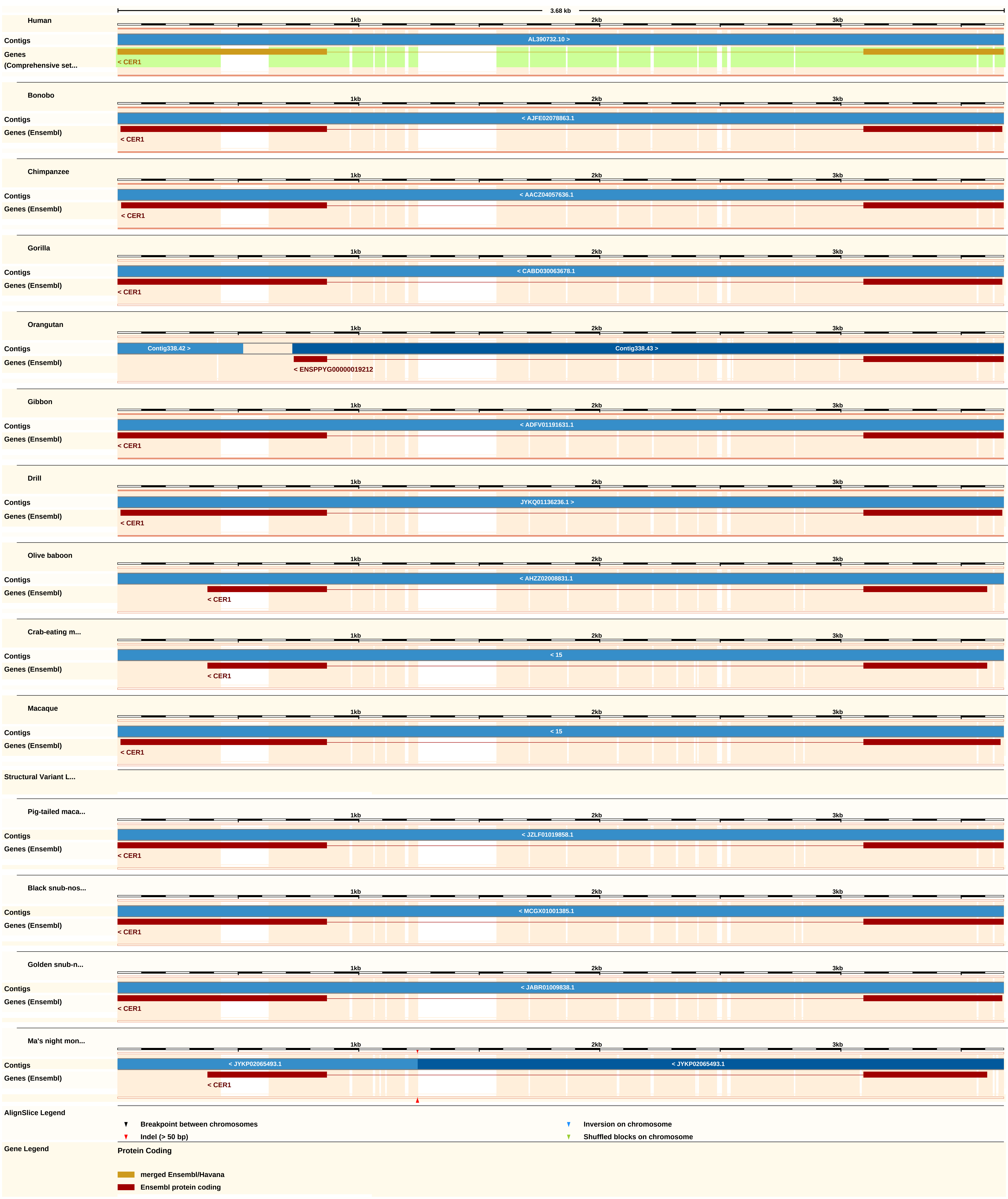
3.68 kb
1kb
2kb
3kb
Human
Contigs
AL390732.10 >
Genes
(Comprehensive set...
< CER1
Bonobo
Contigs
< AJFE02078863.1
Genes (Ensembl)
< CER1
Chimpanzee
Contigs
< AACZ04057636.1
Genes (Ensembl)
< CER1
Gorilla
Contigs
< CABD030063678.1
Genes (Ensembl)
< CER1
Orangutan
Contigs
Contig338.42 >
Contig338.43 >
Genes (Ensembl)
< ENSPPYG00000019212
Gibbon
Contigs
< ADFV01191631.1
Genes (Ensembl)
< CER1
Drill
Contigs
JYKQ01136236.1 >
Genes (Ensembl)
< CER1
Olive baboon
Contigs
< AHZZ02008831.1
Genes (Ensembl)
< CER1
Crab-eating m...
Contigs
< 15
Genes (Ensembl)
< CER1
Macaque
Contigs
< 15
Genes (Ensembl)
< CER1
Structural Variant L...
Pig-tailed maca...
Contigs
< JZLF01019858.1
Genes (Ensembl)
< CER1
Black snub-nos...
Contigs
< MCGX01001385.1
Genes (Ensembl)
< CER1
Golden snub-n...
Contigs
< JABR01009838.1
Genes (Ensembl)
< CER1
Ma's night mon...
Contigs
< JYKP02065493.1
< JYKP02065493.1
Genes (Ensembl)
< CER1
AlignSlice Legend
Breakpoint between chromosomes
Indel (> 50 bp)
Inversion on chromosome
Shuffled blocks on chromosome
Gene Legend
Protein Coding
merged Ensembl/Havana
Ensembl protein coding

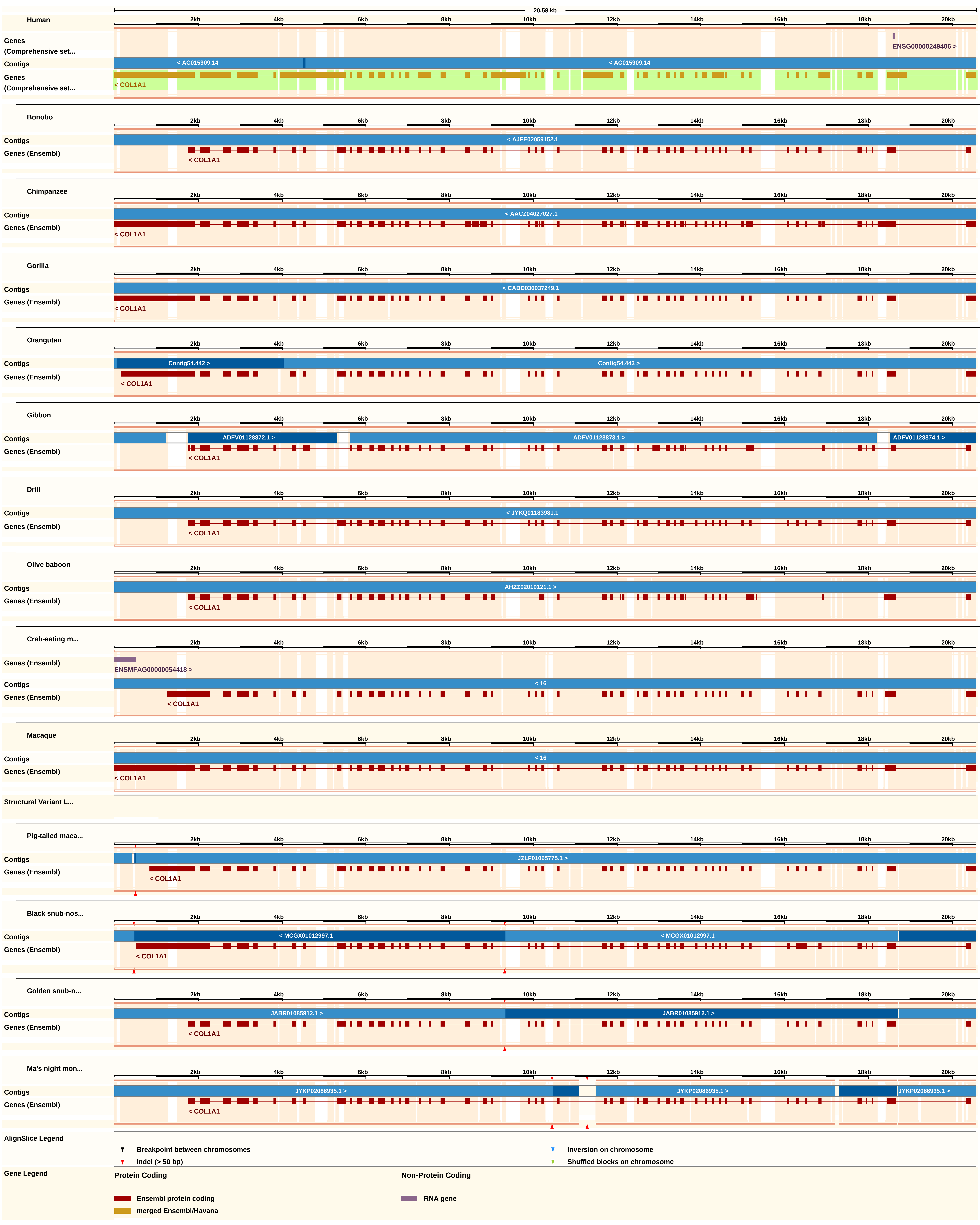

20.58 kb
Human
2kb
4kb
6kb
8kb
10kb
12kb
14kb
16kb
18kb
20kb
Genes (Comprehensive set...
ENSG00000249406 >
Contigs
< AC015909.14
< AC015909.14
Genes (Comprehensive set...
< COL1A1
Bonobo
Contigs
< AJFE02059152.1
Genes (Ensembl)
< COL1A1
Chimpanzee
Contigs
< AACZ04027027.1
Genes (Ensembl)
< COL1A1
Gorilla
Contigs
< CABD030037249.1
Genes (Ensembl)
< COL1A1
Orangutan
Contigs
Contig54.442 >
Contig54.443 >
Genes (Ensembl)
< COL1A1
Gibbon
Contigs
ADFV01128872.1 >
ADFV01128873.1 >
ADFV01128874.1 >
Genes (Ensembl)
< COL1A1
Drill
Contigs
< JYKQ01183981.1
Genes (Ensembl)
< COL1A1
Olive baboon
Contigs
AHZZ02010121.1 >
Genes (Ensembl)
< COL1A1
Crab-eating m...
Genes (Ensembl)
ENSMFAG00000054418 >
Contigs
< 16
Genes (Ensembl)
< COL1A1
Macaque
Contigs
< 16
Genes (Ensembl)
< COL1A1
Structural Variant L...
Pig-tailed maca...
Contigs
JZLF01065775.1 >
Genes (Ensembl)
< COL1A1
Black snub-nos...
Contigs
< MCGX01012997.1
< MCGX01012997.1
Genes (Ensembl)
< COL1A1
Golden snub-n...
Contigs
JABR01085912.1 >
JABR01085912.1 >
Genes (Ensembl)
< COL1A1
Ma's night mon...
Contigs
JYKP02086935.1 >
JYKP02086935.1 >
JYKP02086935.1 >
Genes (Ensembl)
< COL1A1
AlignSlice Legend
Breakpoint between chromosomes
Indel (> 50 bp)
Inversion on chromosome
Shuffled blocks on chromosome
Gene Legend
Protein Coding
Ensembl protein coding
merged Ensembl/Havana
Non-Protein Coding
RNA gene

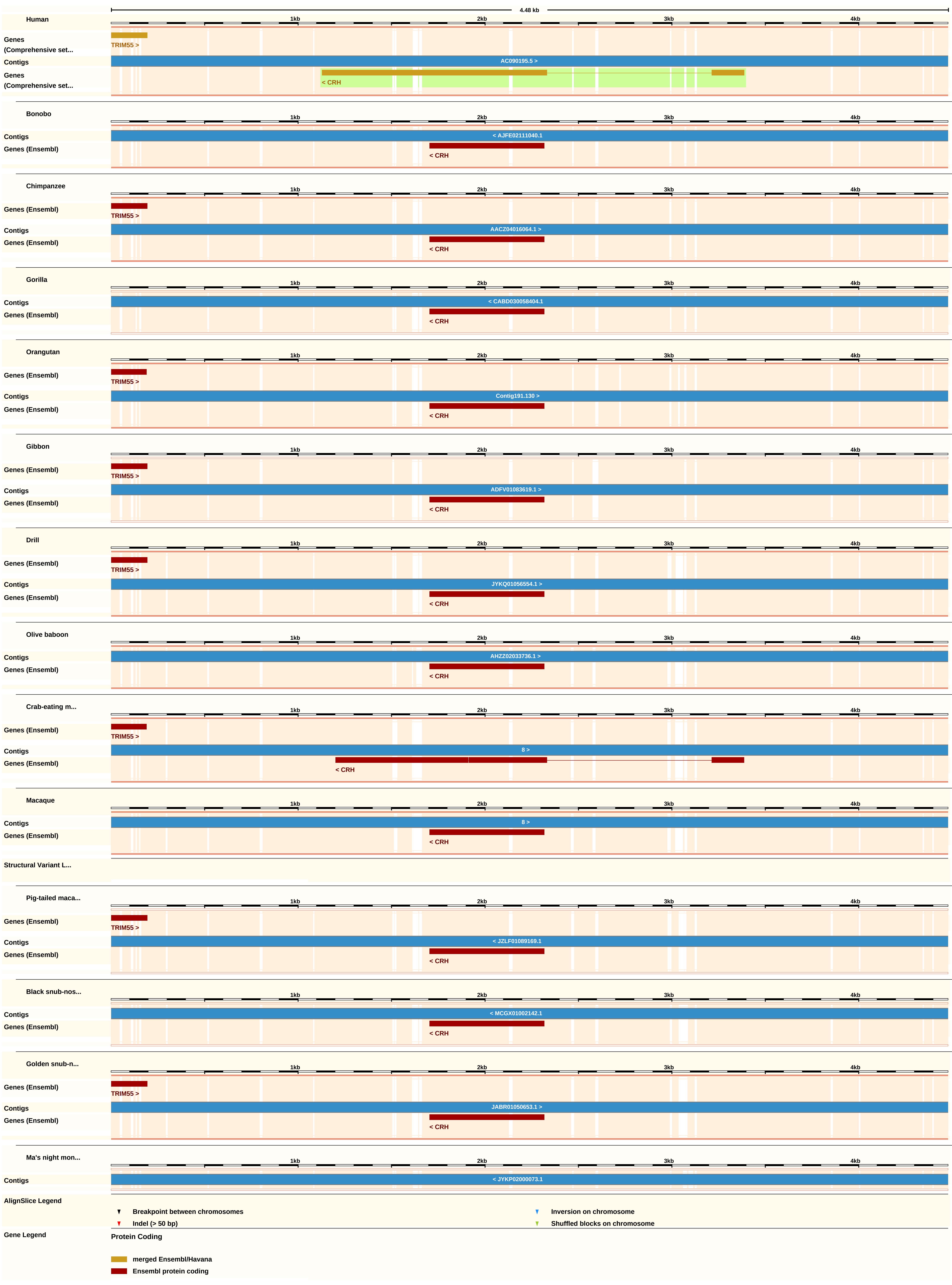
4.48 kb
1kb
2kb
3kb
4kb
Human
Genes (Comprehensive set...
TRIM55 >
Contigs
AC090195.5 >
Genes (Comprehensive set...
< CRH
Bonobo
Contigs
< AJFE02111040.1
Genes (Ensembl)
< CRH
Chimpanzee
Genes (Ensembl)
TRIM55 >
Contigs
AACZ04016064.1 >
Genes (Ensembl)
< CRH
Gorilla
Contigs
< CABD030058404.1
Genes (Ensembl)
< CRH
Orangutan
Genes (Ensembl)
TRIM55 >
Contigs
Contig191.130 >
Genes (Ensembl)
< CRH
Gibbon
Genes (Ensembl)
TRIM55 >
Contigs
ADFV01083619.1 >
Genes (Ensembl)
< CRH
Drill
Genes (Ensembl)
TRIM55 >
Contigs
JYKQ01056554.1 >
Genes (Ensembl)
< CRH
Olive baboon
Contigs
AHZZ02033736.1 >
Genes (Ensembl)
< CRH
Crab-eating m...
Genes (Ensembl)
TRIM55 >
Contigs
8 >
Genes (Ensembl)
< CRH
Macaque
Contigs
8 >
Genes (Ensembl)
< CRH
Structural Variant L...
Pig-tailed maca...
Genes (Ensembl)
TRIM55 >
Contigs
< JZLF01089169.1
Genes (Ensembl)
< CRH
Black snub-nos...
Contigs
< MCGX01002142.1
Genes (Ensembl)
< CRH
Golden snub-n...
Genes (Ensembl)
TRIM55 >
Contigs
JABR01050653.1 >
Genes (Ensembl)
< CRH
Ma's night mon...
Contigs
< JYKP02000073.1
AlignSlice Legend
Breakpoint between chromosomes
Inversion on chromosome
Indel (> 50 bp)
Shuffled blocks on chromosome
Gene Legend
Protein Coding
merged Ensembl/Havana
Ensembl protein coding

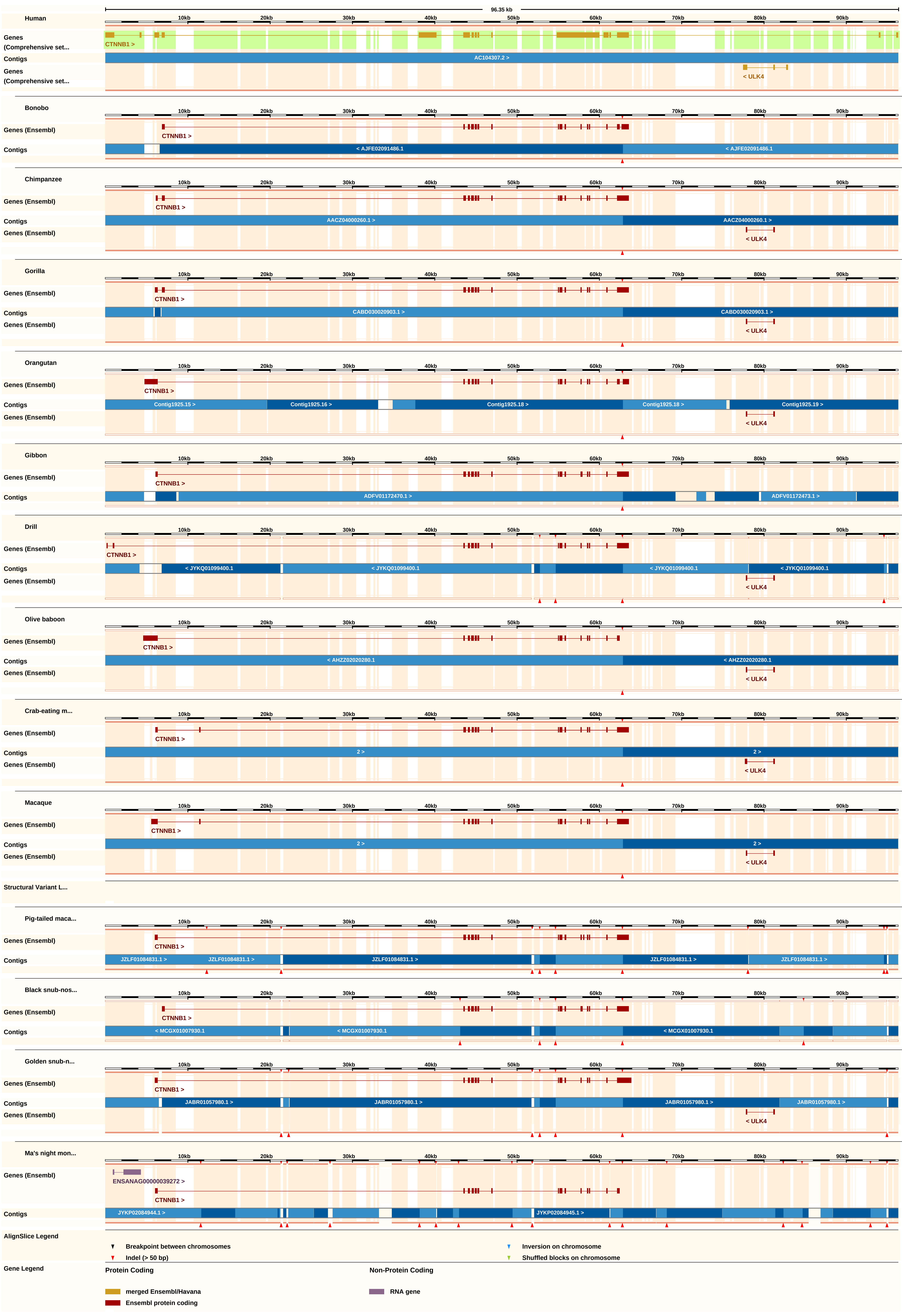

96.35 kb
Human
Genes (Comprehensive set...
CTNNB1 >
Contigs
AC104307.2 >
< ULK4
Bonobo
Genes (Ensembl)
< AJFE02091486.1
Chimpanzee
AACZ04000260.1 >
Gorilla
CABD030020903.1 >
Orangutan
Contig1925.15 >
Contig1925.16 >
Contig1925.18 >
Contig1925.19 >
Gibbon
ADFV01172470.1 >
ADFV01172473.1 >
Drill
< JYKQ01099400.1
Olive baboon
< AHZZ02020280.1
Crab-eating m...
2 >
Macaque
Structural Variant L...
Pig-tailed maca...
JZLF01084831.1 >
Black snub-nos...
< MCGX01007930.1
Golden snub-n...
JABR01057980.1 >
Ma's night mon...
ENSANAG00000039272 >
JYKP02084944.1 >
JYKP02084945.1 >
AlignSlice Legend
Breakpoint between chromosomes
Indel (> 50 bp)
Inversion on chromosome
Shuffled blocks on chromosome
Gene Legend
Protein Coding
merged Ensembl/Havana
Ensembl protein coding
Non-Protein Coding
RNA gene

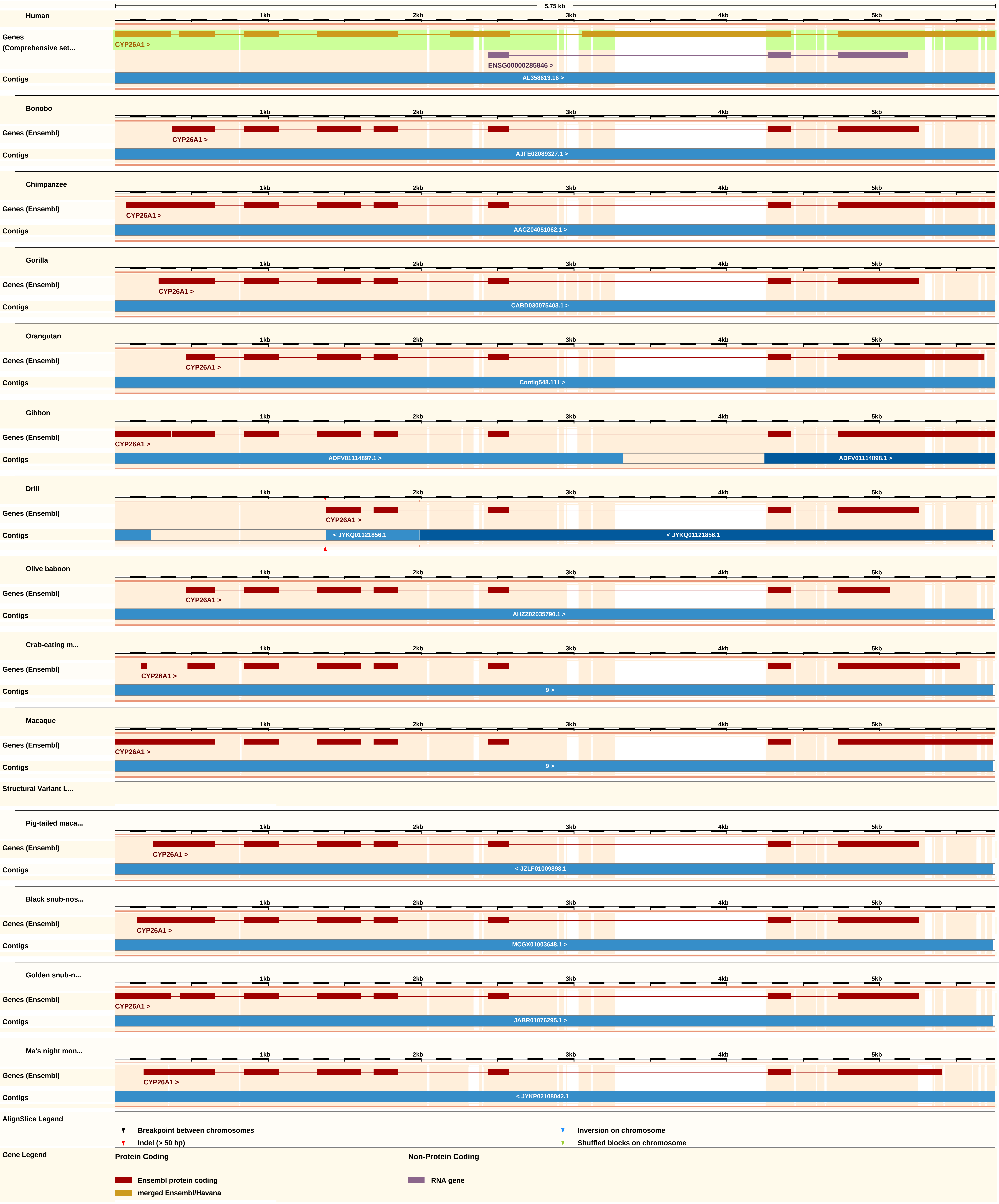
5.75 kb
Human
1kb
2kb
3kb
4kb
5kb
Genes (Comprehensive set...
CYP26A1 >
ENSG00000285846 >
Contigs
AL358613.16 >
Bonobo
Genes (Ensembl)
CYP26A1 >
AJFE02089327.1 >
Chimpanzee
AACZ04051062.1 >
Gorilla
CABD030075403.1 >
Orangutan
Contig548.111 >
Gibbon
ADFV01114897.1 >
ADFV01114898.1 >
Drill
< JYKQ01121856.1
Olive baboon
AHZZ02035790.1 >
Crab-eating m...
9 >
Macaque
Structural Variant L...
Pig-tailed maca...
< JZLF01009898.1
Black snub-nos...
MCGX01003648.1 >
Golden snub-n...
JABR01076295.1 >
Ma's night mon...
< JYKP02108042.1
AlignSlice Legend
Breakpoint between chromosomes
Indel (> 50 bp)
Inversion on chromosome
Shuffled blocks on chromosome
Gene Legend
Protein Coding
Ensembl protein coding
merged Ensembl/Havana
Non-Protein Coding
RNA gene

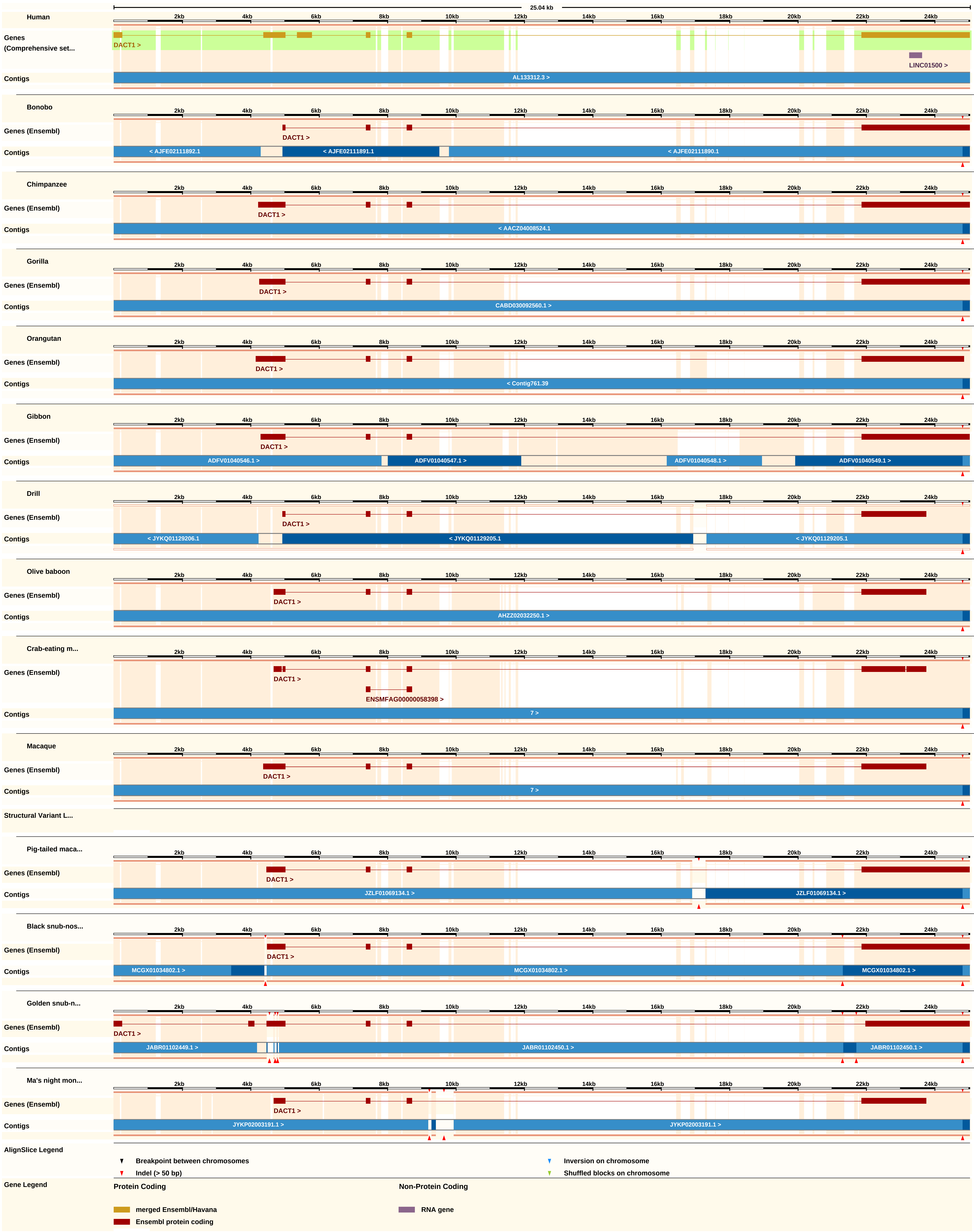
25.04 kb
2kb 4kb 6kb 8kb 10kb 12kb 14kb 16kb 18kb 20kb 22kb 24kb
Human
Genes (Comprehensive set...
DACT1 >
LINC01500 >
Contigs
AL133312.3 >
Bonobo
Genes (Ensembl)
DACT1 >
Contigs
< AJFE02111892.1
< AJFE02111891.1
< AJFE02111890.1
Chimpanzee
Genes (Ensembl)
DACT1 >
Contigs
< AACZ04008524.1
Gorilla
Genes (Ensembl)
DACT1 >
Contigs
CABD030092560.1 >
Orangutan
Genes (Ensembl)
DACT1 >
Contigs
< Contig761.39
Gibbon
Genes (Ensembl)
DACT1 >
Contigs
ADFV01040546.1 >
ADFV01040547.1 >
ADFV01040548.1 >
ADFV01040549.1 >
Drill
Genes (Ensembl)
DACT1 >
Contigs
< JYKQ01129206.1
< JYKQ01129205.1
< JYKQ01129205.1
Olive baboon
Genes (Ensembl)
DACT1 >
Contigs
AHZZ02032250.1 >
Crab-eating m...
Genes (Ensembl)
DACT1 >
ENSMFAG00000058398 >
Contigs
7 >
Macaque
Genes (Ensembl)
DACT1 >
Contigs
7 >
Structural Variant L...
Pig-tailed maca...
Genes (Ensembl)
DACT1 >
Contigs
JZLF01069134.1 >
JZLF01069134.1 >
Black snub-nos...
Genes (Ensembl)
DACT1 >
Contigs
MCGX01034802.1 >
MCGX01034802.1 >
MCGX01034802.1 >
Golden snub-n...
Genes (Ensembl)
DACT1 >
Contigs
JABR01102449.1 >
JABR01102450.1 >
JABR01102450.1 >
Ma's night mon...
Genes (Ensembl)
DACT1 >
Contigs
JYKP02003191.1 >
JYKP02003191.1 >
AlignSlice Legend
Breakpoint between chromosomes
Inversion on chromosome
Indel (> 50 bp)
Shuffled blocks on chromosome
Gene Legend
Protein Coding
Non-Protein Coding
merged Ensembl/Havana
RNA gene
Ensembl protein coding

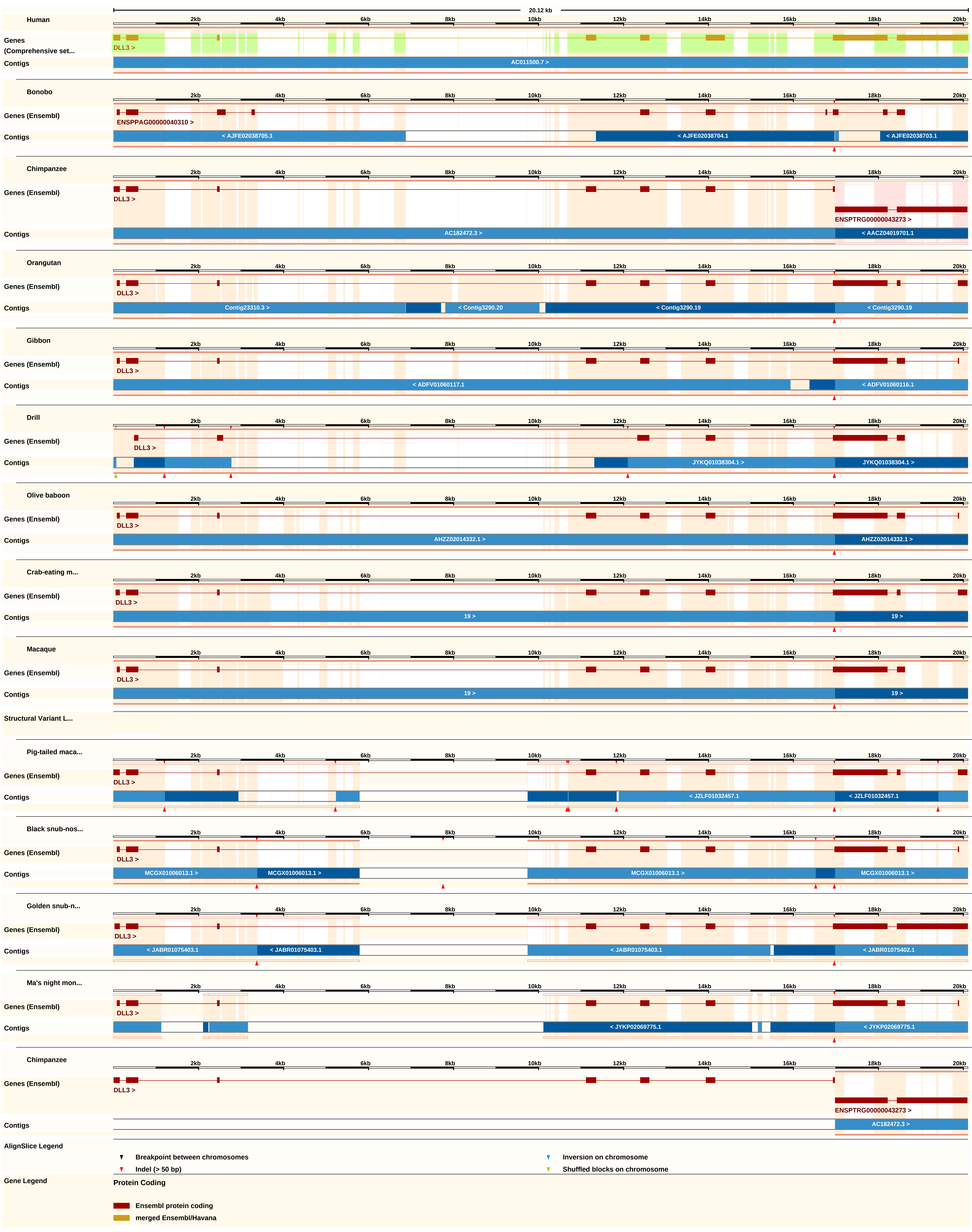
20.12 kb
Human
Genes (Comprehensive set...
DLL3 >
Contigs
AC011500.7 >
Bonobo
Genes (Ensembl)
ENSPPAG00000040310 >
< AJFE02038705.1
< AJFE02038704.1
< AJFE02038703.1
Chimpanzee
DLL3 >
ENSPTRG00000043273 >
AC182472.3 >
< AACZ04019701.1
Orangutan
Contig23310.3 >
< Contig3290.20
< Contig3290.19
Gibbon
< ADFV01060117.1
< ADFV01060116.1
Drill
JYKQ01038304.1 >
Olive baboon
AHZZ02014332.1 >
Crab-eating m...
19 >
Macaque
Structural Variant L...
Pig-tailed maca...
< JZLF01032457.1
Black snub-nos...
MCGX01006013.1 >
Golden snub-n...
< JABR01075403.1
< JABR01075402.1
Ma's night mon...
< JYKP02069775.1
AlignSlice Legend
Breakpoint between chromosomes
Indel (> 50 bp)
Inversion on chromosome
Shuffled blocks on chromosome
Gene Legend
Protein Coding
Ensembl protein coding
merged Ensembl/Havana

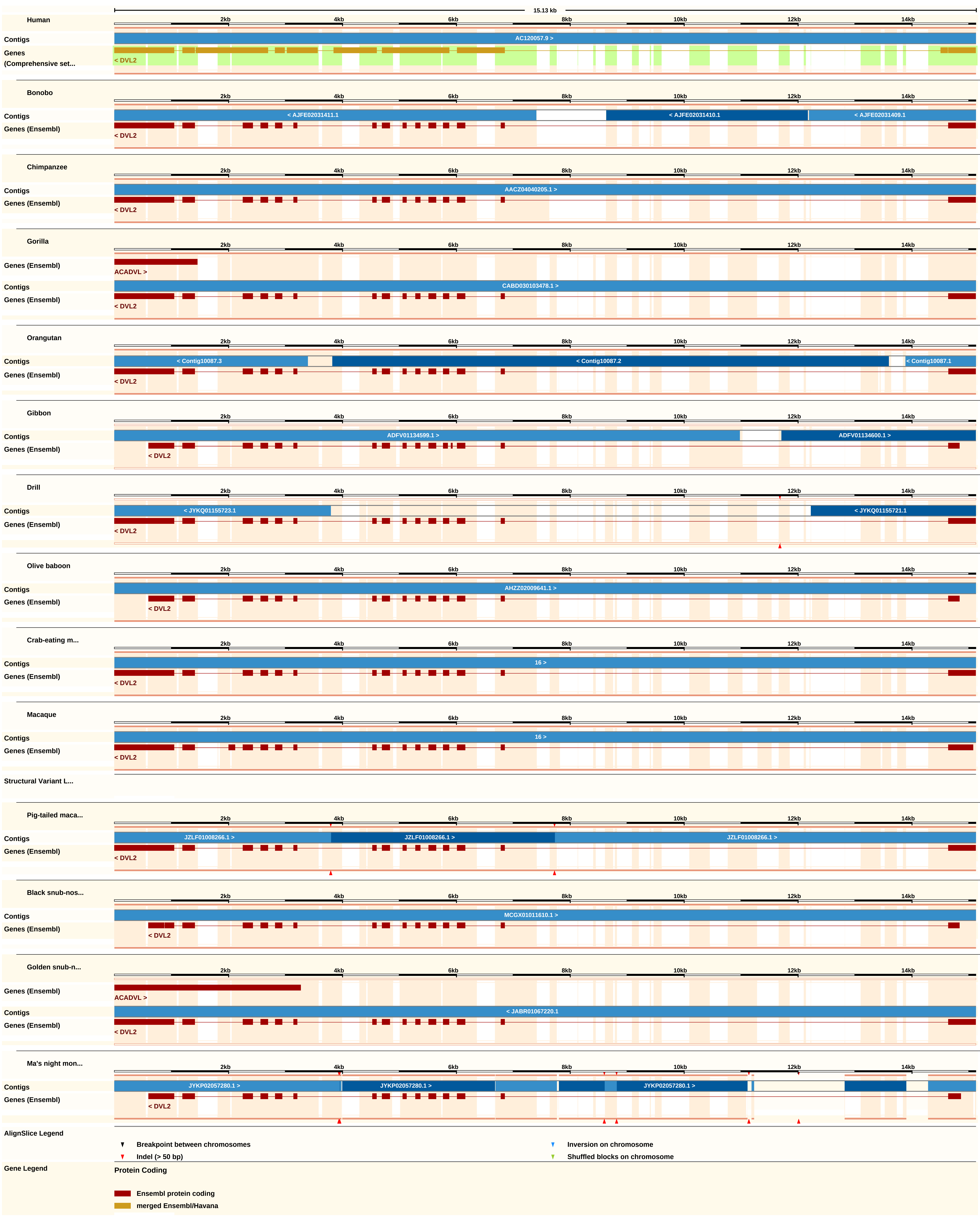
15.13 kb
Human
Contigs
AC120057.9 >
Genes (Comprehensive set...
< DVL2
Bonobo
Contigs
< AJFE02031411.1
< AJFE02031410.1
< AJFE02031409.1
Genes (Ensembl)
< DVL2
Chimpanzee
Contigs
AACZ04040205.1 >
Genes (Ensembl)
< DVL2
Gorilla
Genes (Ensembl)
ACADVL >
Contigs
CABD030103478.1 >
Genes (Ensembl)
< DVL2
Orangutan
Contigs
< Contig10087.3
< Contig10087.2
< Contig10087.1
Genes (Ensembl)
< DVL2
Gibbon
Contigs
ADFV01134599.1 >
ADFV01134600.1 >
Genes (Ensembl)
< DVL2
Drill
Contigs
< JYKQ01155723.1
< JYKQ01155721.1
Genes (Ensembl)
< DVL2
Olive baboon
Contigs
AHZZ02009641.1 >
Genes (Ensembl)
< DVL2
Crab-eating m...
Contigs
16 >
Genes (Ensembl)
< DVL2
Macaque
Contigs
16 >
Genes (Ensembl)
< DVL2
Structural Variant L...
Pig-tailed maca...
Contigs
JZLF01008266.1 >
JZLF01008266.1 >
JZLF01008266.1 >
Genes (Ensembl)
< DVL2
Black snub-nos...
Contigs
MCGX01011610.1 >
Genes (Ensembl)
< DVL2
Golden snub-n...
Genes (Ensembl)
ACADVL >
Contigs
< JABR01067220.1
Genes (Ensembl)
< DVL2
Ma's night mon...
Contigs
JYKP02057280.1 >
JYKP02057280.1 >
JYKP02057280.1 >
Genes (Ensembl)
< DVL2
2kb
4kb
6kb
8kb
10kb
12kb
14kb
AlignSlice Legend
Breakpoint between chromosomes
Indel (> 50 bp)
Inversion on chromosome
Shuffled blocks on chromosome
Gene Legend
Protein Coding
Ensembl protein coding
merged Ensembl/Havana

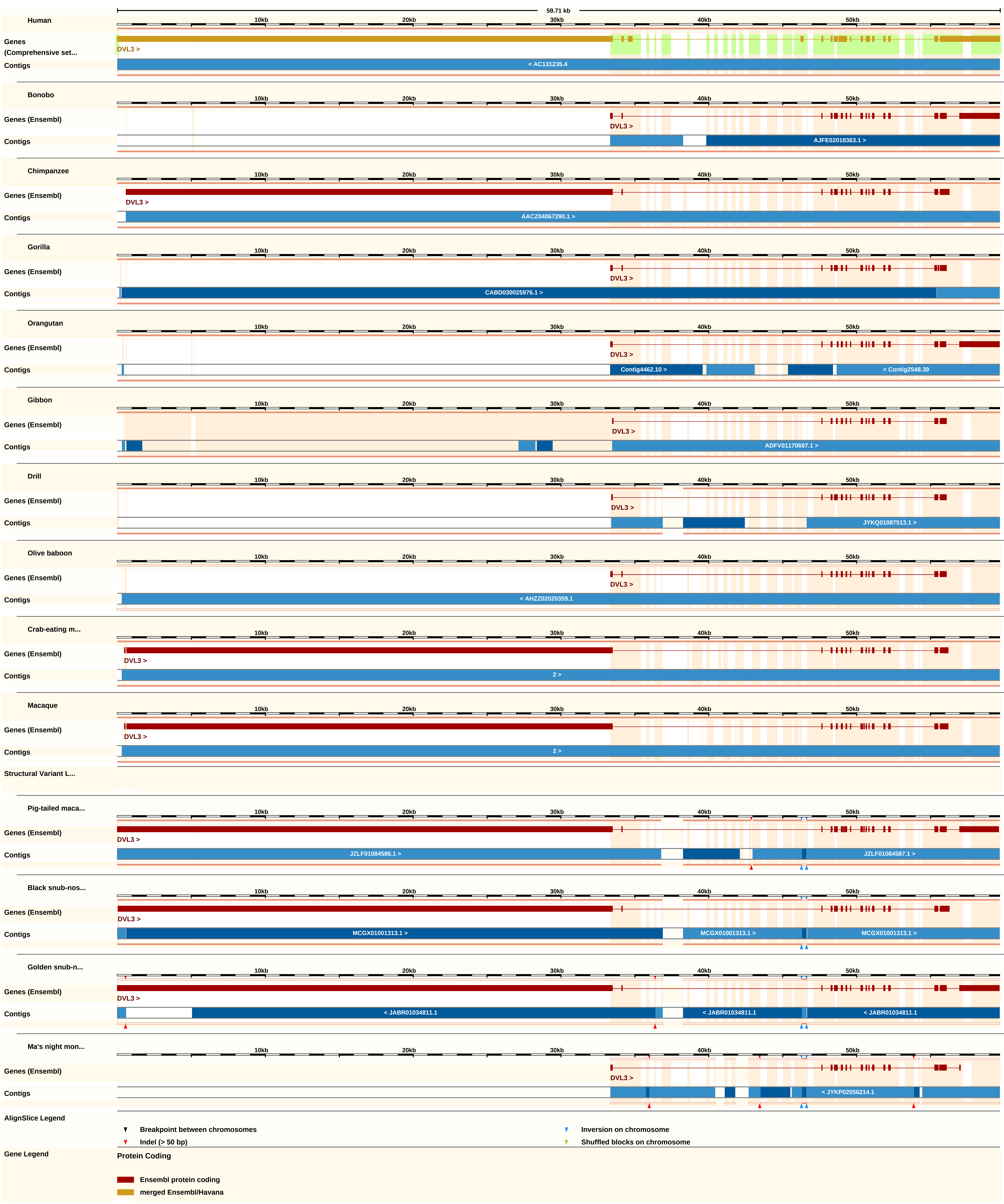

59.71 kb
10kb
20kb
30kb
40kb
50kb
Human
Genes (Comprehensive set...
DVL3 >
Contigs
< AC131235.4
Bonobo
Genes (Ensembl)
DVL3 >
Contigs
AJFE02018363.1 >
Chimpanzee
Genes (Ensembl)
DVL3 >
Contigs
AACZ04067290.1 >
Gorilla
Genes (Ensembl)
DVL3 >
Contigs
CABD030025976.1 >
Orangutan
Genes (Ensembl)
DVL3 >
Contigs
Contig4462.10 >
< Contig2548.39
Gibbon
Genes (Ensembl)
DVL3 >
Contigs
ADFV01170697.1 >
Drill
Genes (Ensembl)
DVL3 >
Contigs
JYKQ01087513.1 >
Olive baboon
Genes (Ensembl)
DVL3 >
Contigs
< AHZZ02020359.1
Crab-eating m...
Genes (Ensembl)
DVL3 >
Contigs
2 >
Macaque
Genes (Ensembl)
DVL3 >
Contigs
2 >
Structural Variant L...
Pig-tailed maca...
Genes (Ensembl)
DVL3 >
Contigs
JZLF01084586.1 >
JZLF01084587.1 >
Black snub-nos...
Genes (Ensembl)
DVL3 >
Contigs
MCGX01001313.1 >
MCGX01001313.1 >
MCGX01001313.1 >
Golden snub-n...
Genes (Ensembl)
DVL3 >
Contigs
< JABR01034811.1
< JABR01034811.1
< JABR01034811.1
Ma's night mon...
Genes (Ensembl)
DVL3 >
Contigs
< JYKP02056214.1
AlignSlice Legend
Breakpoint between chromosomes
Inversion on chromosome
Indel (> 50 bp)
Shuffled blocks on chromosome
Gene Legend
Protein Coding
Ensembl protein coding
merged Ensembl/Havana

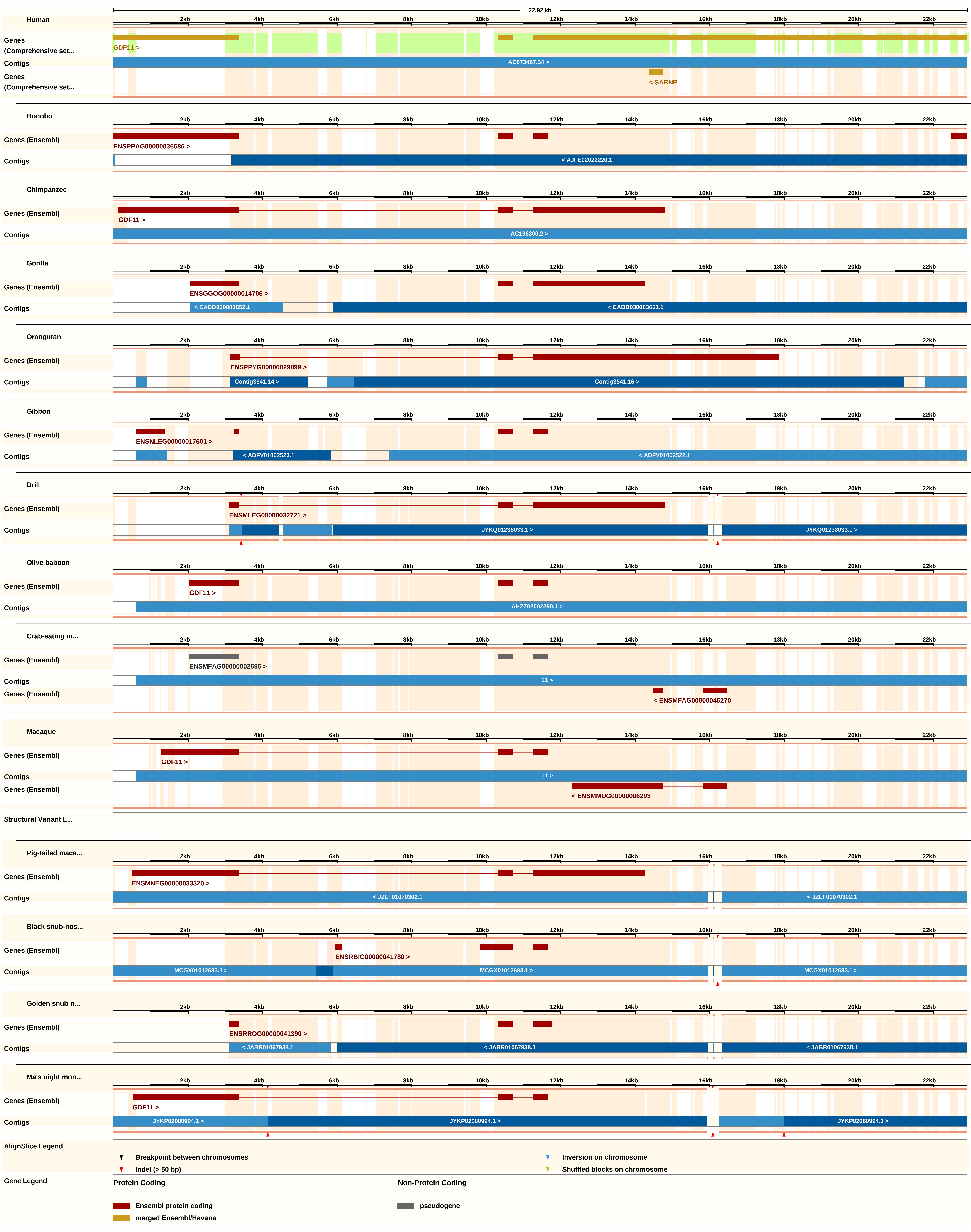
22.92 kb
Human
Genes (Comprehensive set...
GDF11 >
Contigs
AC073487.34 >
< SARNP
Bonobo
Genes (Ensembl)
ENSPPAG00000036686 >
< AJFE02022220.1
Chimpanzee
GDF11 >
AC196300.2 >
Gorilla
ENSGGOG00000014706 >
< CABD030083652.1
< CABD030083651.1
Orangutan
ENSPPYG00000029899 >
Contig3541.14 >
Contig3541.16 >
Gibbon
ENSNLEG00000017601 >
< ADFV01002523.1
< ADFV01002522.1
Drill
ENSMLEG00000032721 >
JYKQ01238033.1 >
Olive baboon
AHZZ02002250.1 >
Crab-eating m...
ENSMFAG00000002695 >
11 >
< ENSMFAG00000045270
Macaque
< ENSMMUG00000006293
Structural Variant L...
Pig-tailed maca...
ENSMNEG00000033320 >
< JZLF01070302.1
Black snub-nos...
ENSRBIG00000041780 >
MCGX01012683.1 >
Golden snub-n...
ENSRROG00000041390 >
< JABR01067938.1
Ma's night mon...
JYKP02080994.1 >
AlignSlice Legend
Breakpoint between chromosomes
Indel (> 50 bp)
Inversion on chromosome
Shuffled blocks on chromosome
Gene Legend
Protein Coding
Ensembl protein coding
merged Ensembl/Havana
Non-Protein Coding
pseudogene

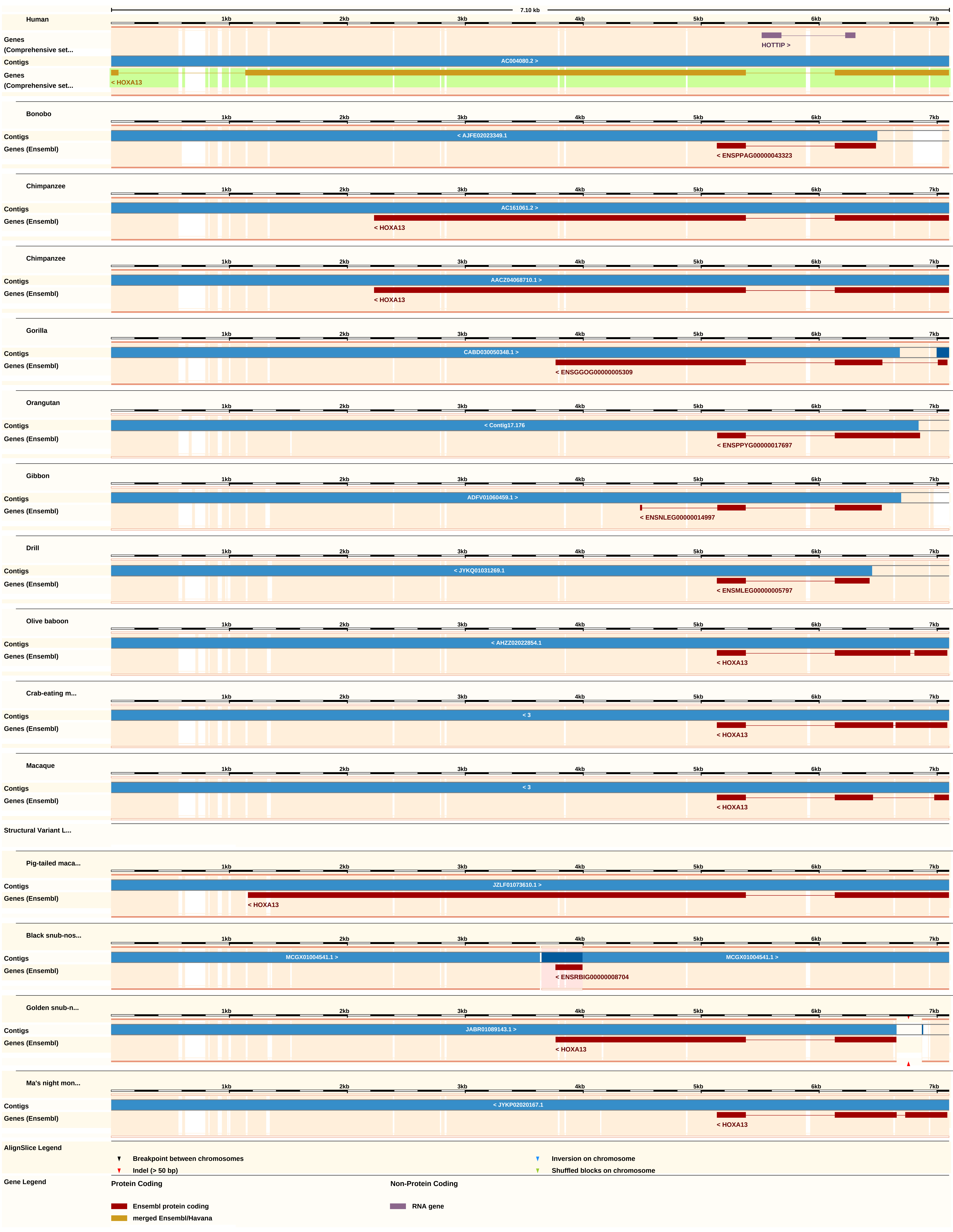
7.10 kb
1kb
2kb
3kb
4kb
5kb
6kb
7kb
Human
Genes (Comprehensive set...
HOTTIP >
Contigs
AC004080.2 >
Genes (Comprehensive set...
< HOXA13
Bonobo
Contigs
< AJFE02023349.1
Genes (Ensembl)
< ENSPPAG00000043323
Chimpanzee
Contigs
AC161061.2 >
Genes (Ensembl)
< HOXA13
Chimpanzee
Contigs
AACZ04068710.1 >
Genes (Ensembl)
< HOXA13
Gorilla
Contigs
CABD030050348.1 >
Genes (Ensembl)
< ENSGGOG00000005309
Orangutan
Contigs
< Contig17.176
Genes (Ensembl)
< ENSPPYG00000017697
Gibbon
Contigs
ADFV01060459.1 >
Genes (Ensembl)
< ENSNLEG00000014997
Drill
Contigs
< JYKQ01031269.1
Genes (Ensembl)
< ENSMLEG00000005797
Olive baboon
Contigs
< AHZZ02022854.1
Genes (Ensembl)
< HOXA13
Crab-eating m...
Contigs
< 3
Genes (Ensembl)
< HOXA13
Macaque
Contigs
< 3
Genes (Ensembl)
< HOXA13
Structural Variant L...
Pig-tailed maca...
Contigs
JZLF01073610.1 >
Genes (Ensembl)
< HOXA13
Black snub-nos...
Contigs
MCGX01004541.1 >
MCGX01004541.1 >
Genes (Ensembl)
< ENSRBIG00000008704
Golden snub-n...
Contigs
JABR01089143.1 >
Genes (Ensembl)
< HOXA13
Ma's night mon...
Contigs
< JYKP02020167.1
Genes (Ensembl)
< HOXA13
AlignSlice Legend
Breakpoint between chromosomes
Inversion on chromosome
Indel (> 50 bp)
Shuffled blocks on chromosome
Gene Legend
Protein Coding
Non-Protein Coding
Ensembl protein coding
RNA gene
merged Ensembl/Havana

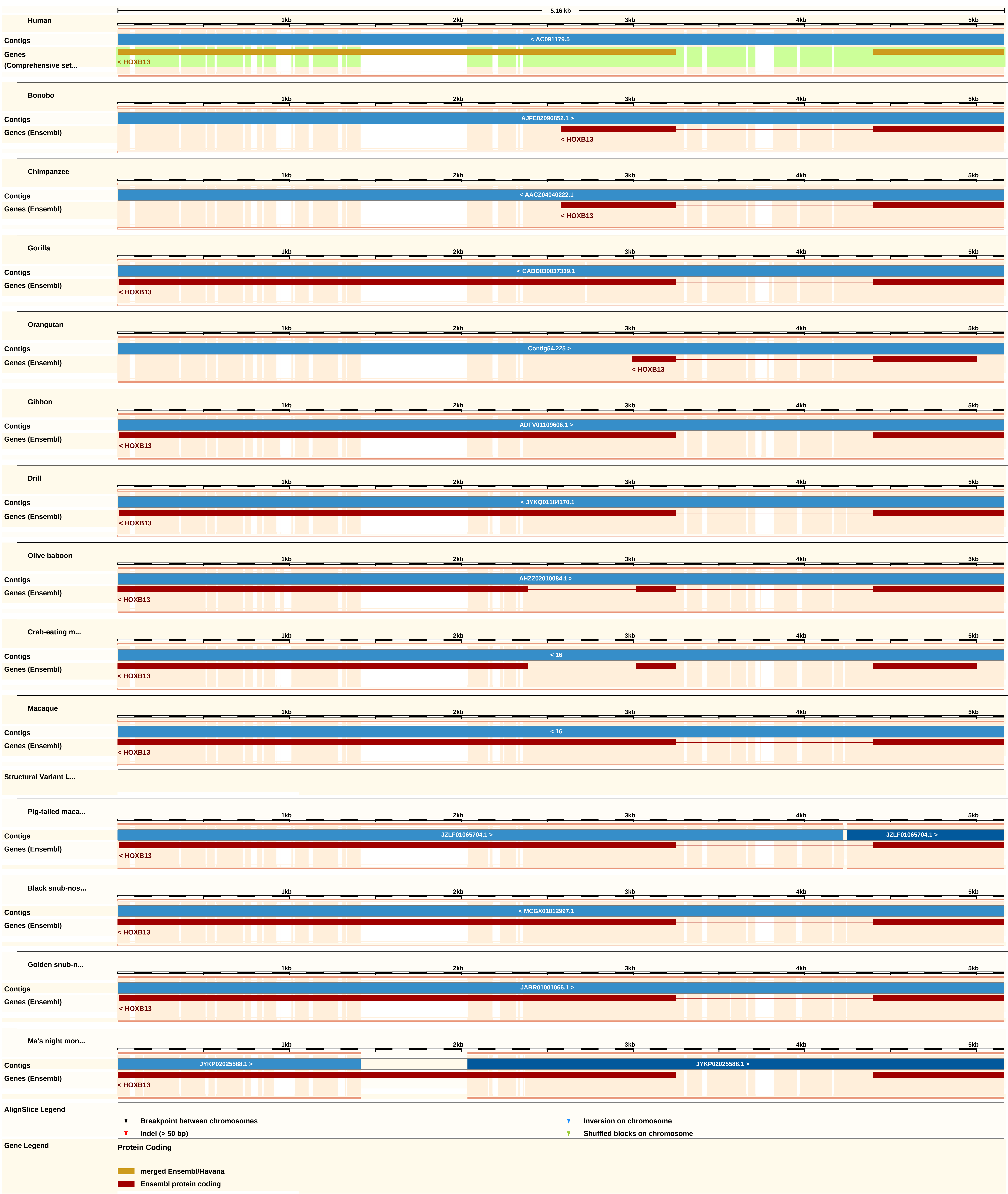
5.16 kb
1kb
2kb
3kb
4kb
5kb
Human
Contigs
< AC091179.5
Genes (Comprehensive set...
< HOXB13
Bonobo
Contigs
AJFE02096852.1 >
Genes (Ensembl)
< HOXB13
Chimpanzee
Contigs
< AACZ04040222.1
Genes (Ensembl)
< HOXB13
Gorilla
Contigs
< CABD030037339.1
Genes (Ensembl)
< HOXB13
Orangutan
Contigs
Contig54.225 >
Genes (Ensembl)
< HOXB13
Gibbon
Contigs
ADFV01109606.1 >
Genes (Ensembl)
< HOXB13
Drill
Contigs
< JYKQ01184170.1
Genes (Ensembl)
< HOXB13
Olive baboon
Contigs
AHZZ02010084.1 >
Genes (Ensembl)
< HOXB13
Crab-eating m...
Contigs
< 16
Genes (Ensembl)
< HOXB13
Macaque
Contigs
< 16
Genes (Ensembl)
< HOXB13
Structural Variant L...
Pig-tailed maca...
Contigs
JZLF01065704.1 >
JZLF01065704.1 >
Genes (Ensembl)
< HOXB13
Black snub-nos...
Contigs
< MCGX01012997.1
Genes (Ensembl)
< HOXB13
Golden snub-n...
Contigs
JABR01001066.1 >
Genes (Ensembl)
< HOXB13
Ma's night mon...
Contigs
JYKP02025588.1 >
JYKP02025588.1 >
Genes (Ensembl)
< HOXB13
AlignSlice Legend
Breakpoint between chromosomes
Inversion on chromosome
Indel (> 50 bp)
Shuffled blocks on chromosome
Gene Legend
Protein Coding
merged Ensembl/Havana
Ensembl protein coding

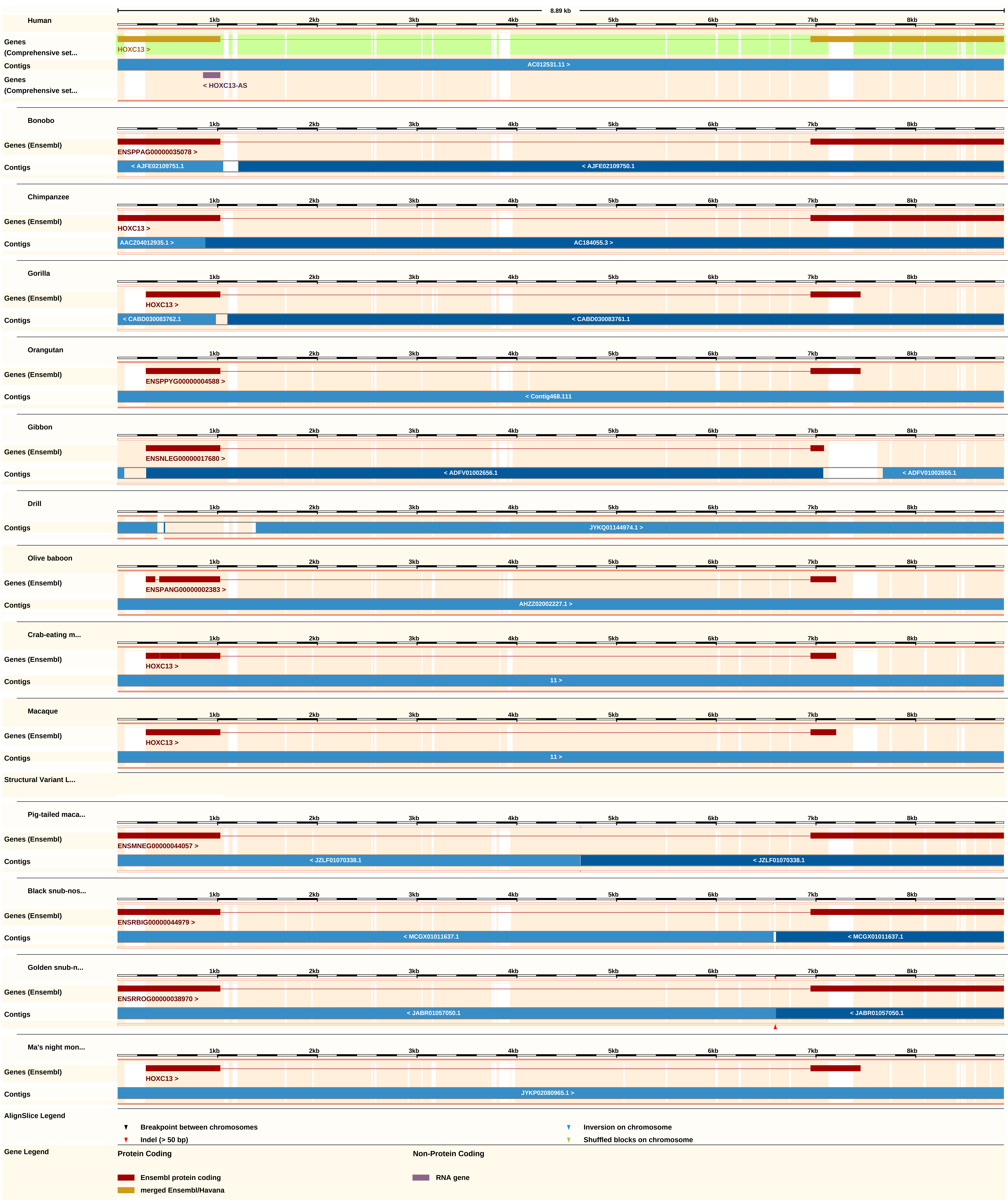
8.89 kb
Human
1kb
2kb
3kb
4kb
5kb
6kb
7kb
8kb
Genes (Comprehensive set...
HOXC13 >
Contigs
AC012531.11 >
Genes (Comprehensive set...
< HOXC13-AS
Bonobo
Genes (Ensembl)
ENSPPAG00000035078 >
Contigs
< AJFE02109751.1
< AJFE02109750.1
Chimpanzee
Genes (Ensembl)
HOXC13 >
Contigs
AACZ04012935.1 >
AC184055.3 >
Gorilla
Genes (Ensembl)
HOXC13 >
Contigs
< CABD030083762.1
< CABD030083761.1
Orangutan
Genes (Ensembl)
ENSPPYG00000004588 >
Contigs
< Contig468.111
Gibbon
Genes (Ensembl)
ENSNLEG00000017680 >
Contigs
< ADFV01002656.1
< ADFV01002655.1
Drill
Contigs
JYKQ01144974.1 >
Olive baboon
Genes (Ensembl)
ENSPANG00000002383 >
Contigs
AHZZ02002227.1 >
Crab-eating m...
Genes (Ensembl)
HOXC13 >
Contigs
11 >
Macaque
Genes (Ensembl)
HOXC13 >
Contigs
11 >
Structural Variant L...
Pig-tailed maca...
Genes (Ensembl)
ENSMNEG00000044057 >
Contigs
< JZLF01070338.1
< JZLF01070338.1
Black snub-nos...
Genes (Ensembl)
ENSRBIG00000044979 >
Contigs
< MCGX01011637.1
< MCGX01011637.1
Golden snub-n...
Genes (Ensembl)
ENSRROG00000038970 >
Contigs
< JABR01057050.1
< JABR01057050.1
Ma's night mon...
Genes (Ensembl)
HOXC13 >
Contigs
JYKP02080965.1 >
AlignSlice Legend
Breakpoint between chromosomes
Inversion on chromosome
Indel (> 50 bp)
Shuffled blocks on chromosome
Gene Legend
Protein Coding
Non-Protein Coding
Ensembl protein coding
RNA gene
merged Ensembl/Havana

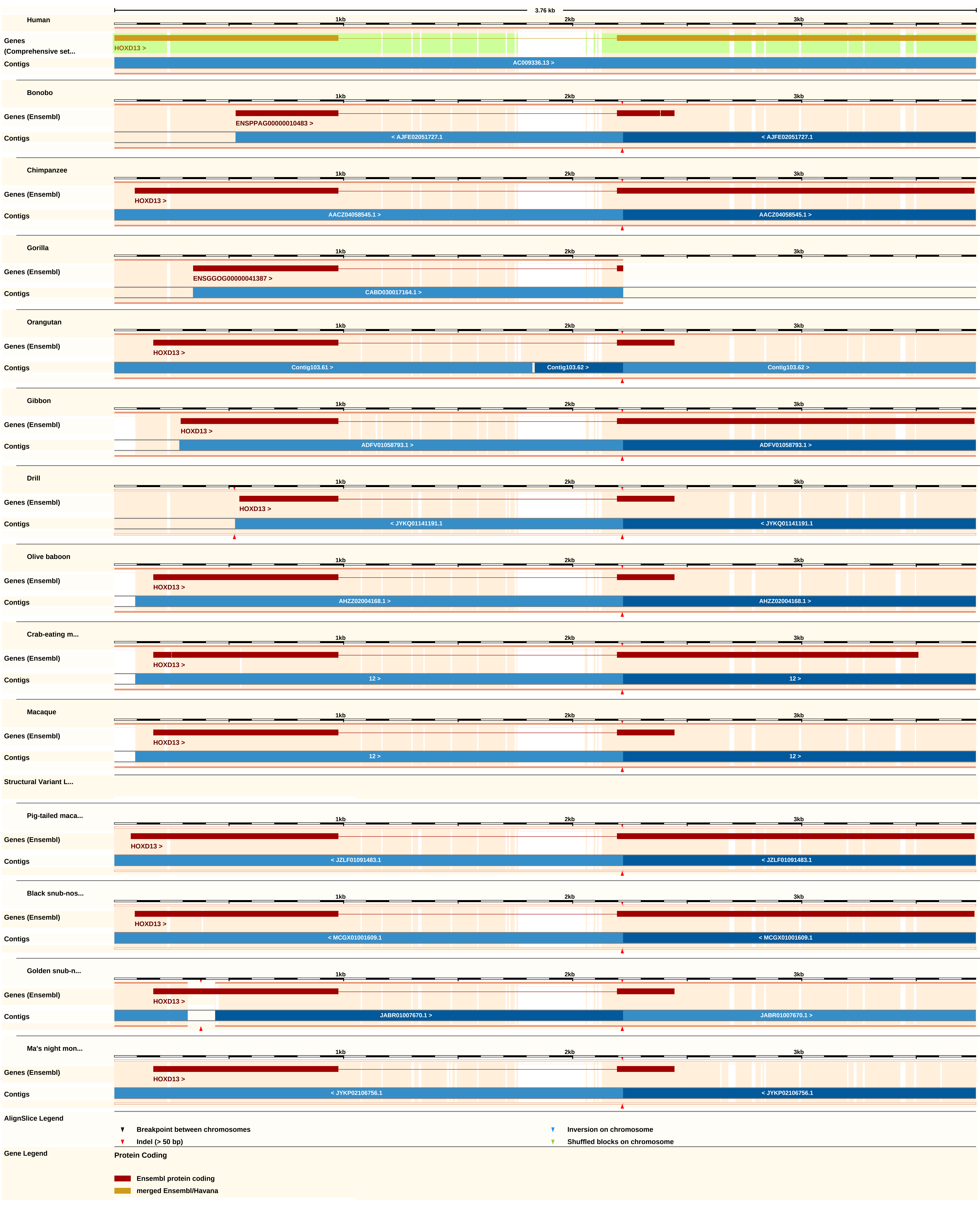
3.76 kb
Human
Genes (Comprehensive set...
HOXD13 >
Contigs
AC009336.13 >
Bonobo
Genes (Ensembl)
ENSPPAG00000010483 >
< AJFE02051727.1
Chimpanzee
HOXD13 >
AACZ04058545.1 >
Gorilla
ENSGGOG00000041387 >
CABD030017164.1 >
Orangutan
Contig103.61 >
Contig103.62 >
Gibbon
ADFV01058793.1 >
Drill
< JYKQ01141191.1
Olive baboon
AHZZ02004168.1 >
Crab-eating m...
12 >
Macaque
Structural Variant L...
Pig-tailed maca...
< JZLF01091483.1
Black snub-nos...
< MCGX01001609.1
Golden snub-n...
JABR01007670.1 >
Ma's night mon...
< JYKP02106756.1
1kb
2kb
3kb
AlignSlice Legend
Breakpoint between chromosomes
Indel (> 50 bp)
Inversion on chromosome
Shuffled blocks on chromosome
Gene Legend
Protein Coding
Ensembl protein coding
merged Ensembl/Havana

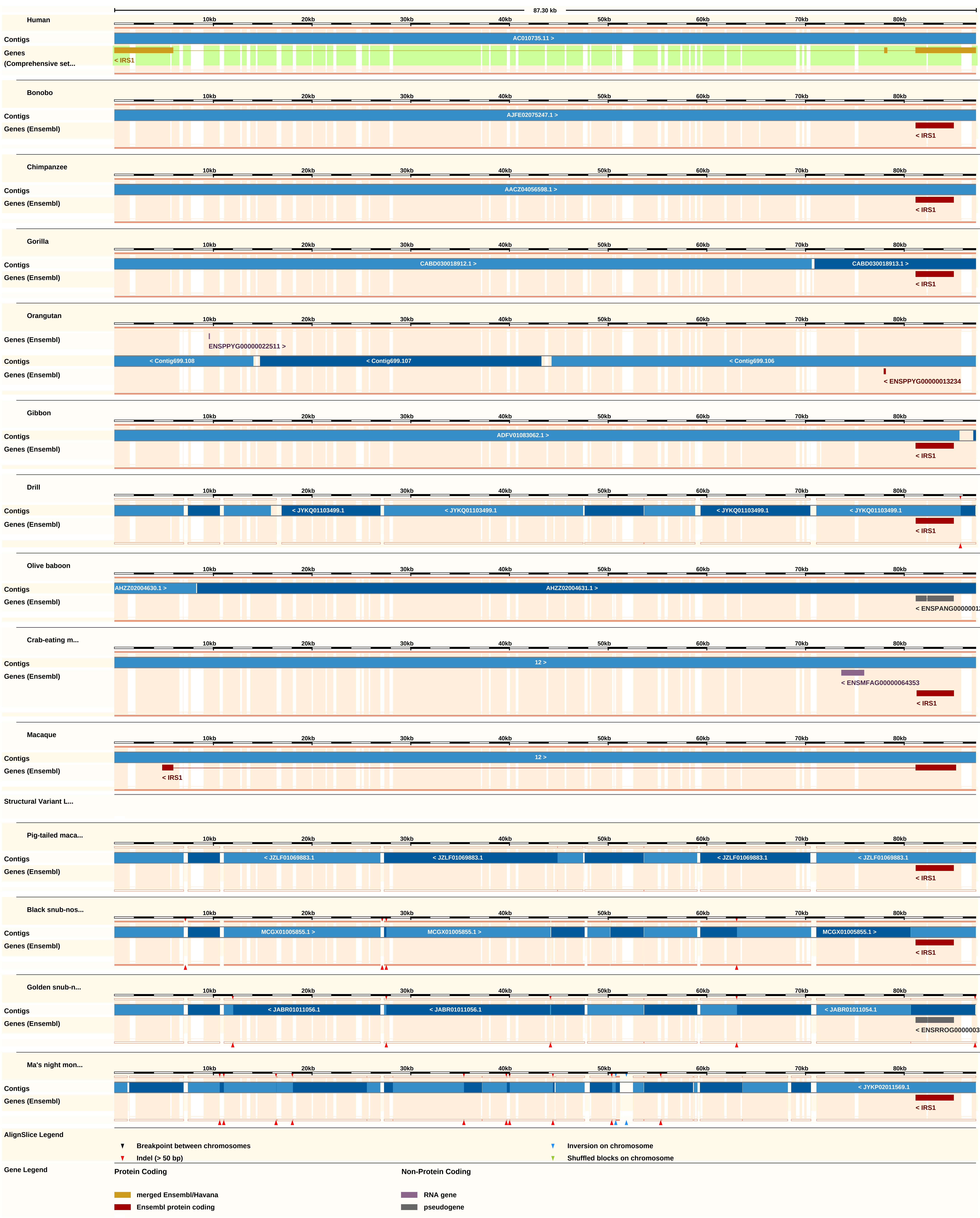
87.30 kb
10kb
20kb
30kb
40kb
50kb
60kb
70kb
80kb
Human
Contigs
AC010735.11 >
Genes
(Comprehensive set...
< IRS1
Bonobo
Contigs
AJFE02075247.1 >
Genes (Ensembl)
< IRS1
Chimpanzee
Contigs
AACZ04056598.1 >
Genes (Ensembl)
< IRS1
Gorilla
Contigs
CABD030018912.1 >
CABD030018913.1 >
Genes (Ensembl)
< IRS1
Orangutan
Genes (Ensembl)
ENSPPYG00000022511 >
Contigs
< Contig699.108
< Contig699.107
< Contig699.106
Genes (Ensembl)
< ENSPPYG00000013234
Gibbon
Contigs
ADFV01083062.1 >
Genes (Ensembl)
< IRS1
Drill
Contigs
< JYKQ01103499.1
< JYKQ01103499.1
< JYKQ01103499.1
< JYKQ01103499.1
Genes (Ensembl)
< IRS1
Olive baboon
Contigs
AHZZ02004630.1 >
AHZZ02004631.1 >
Genes (Ensembl)
< ENSPANG0000001
Crab-eating m...
Contigs
12 >
Genes (Ensembl)
< ENSMFAG00000064353
< IRS1
Macaque
Contigs
12 >
Genes (Ensembl)
< IRS1
Structural Variant L...
Pig-tailed maca...
Contigs
< JZLF01069883.1
< JZLF01069883.1
< JZLF01069883.1
< JZLF01069883.1
Genes (Ensembl)
< IRS1
Black snub-nos...
Contigs
MCGX01005855.1 >
MCGX01005855.1 >
MCGX01005855.1 >
Genes (Ensembl)
< IRS1
Golden snub-n...
Contigs
< JABR01011056.1
< JABR01011056.1
< JABR01011054.1
Genes (Ensembl)
< ENSRROG0000003
Ma's night mon...
Contigs
< JYKP02011569.1
Genes (Ensembl)
< IRS1
AlignSlice Legend
Breakpoint between chromosomes
Indel (> 50 bp)
Inversion on chromosome
Shuffled blocks on chromosome
Gene Legend
Protein Coding
merged Ensembl/Havana
Ensembl protein coding
Non-Protein Coding
RNA gene
pseudogene

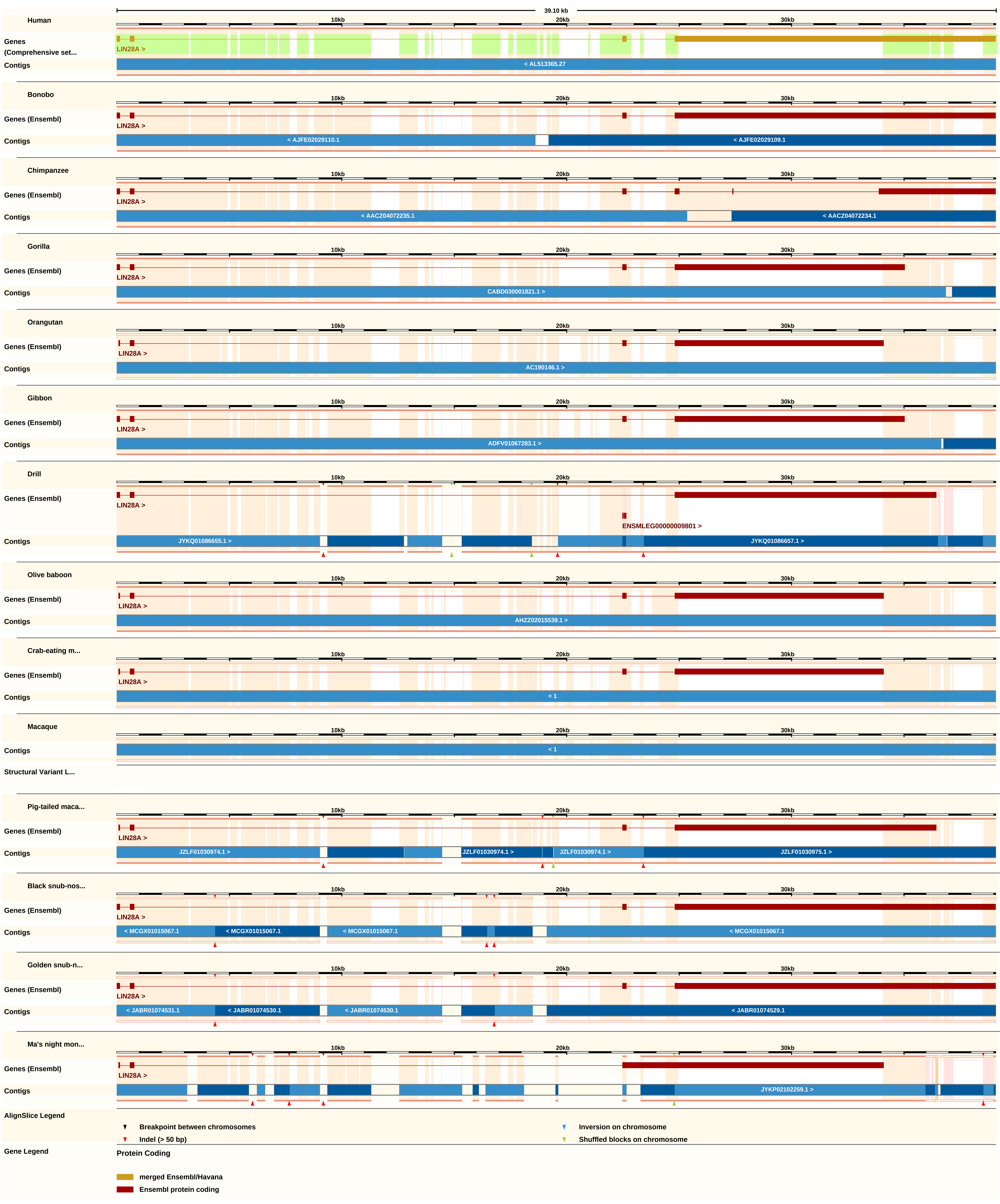
39.10 kb
10kb
20kb
30kb
Human
Genes (Comprehensive set...
LIN28A >
Contigs
< AL513365.27
Bonobo
Genes (Ensembl)
LIN28A >
Contigs
< AJFE02029110.1
< AJFE02029109.1
Chimpanzee
Genes (Ensembl)
LIN28A >
Contigs
< AACZ04072235.1
< AACZ04072234.1
Gorilla
Genes (Ensembl)
LIN28A >
Contigs
CABD030001821.1 >
Orangutan
Genes (Ensembl)
LIN28A >
Contigs
AC190146.1 >
Gibbon
Genes (Ensembl)
LIN28A >
Contigs
ADFV01067283.1 >
Drill
Genes (Ensembl)
LIN28A >
ENSMLEG00000009801 >
Contigs
JYKQ01086655.1 >
JYKQ01086657.1 >
Olive baboon
Genes (Ensembl)
LIN28A >
Contigs
AHZZ02015539.1 >
Crab-eating m...
Genes (Ensembl)
LIN28A >
Contigs
< 1
Macaque
Contigs
< 1
Structural Variant L...
Pig-tailed maca...
Genes (Ensembl)
LIN28A >
Contigs
JZLF01030974.1 >
JZLF01030974.1 >
JZLF01030974.1 >
JZLF01030975.1 >
Black snub-nos...
Genes (Ensembl)
LIN28A >
Contigs
< MCGX01015067.1
< MCGX01015067.1
< MCGX01015067.1
< MCGX01015067.1
Golden snub-n...
Genes (Ensembl)
LIN28A >
Contigs
< JABR01074531.1
< JABR01074530.1
< JABR01074530.1
< JABR01074529.1
Ma's night mon...
Genes (Ensembl)
LIN28A >
Contigs
JYKP02102259.1 >
AlignSlice Legend
Breakpoint between chromosomes
Indel (> 50 bp)
Inversion on chromosome
Shuffled blocks on chromosome
Gene Legend
Protein Coding
merged Ensembl/Havana
Ensembl protein coding

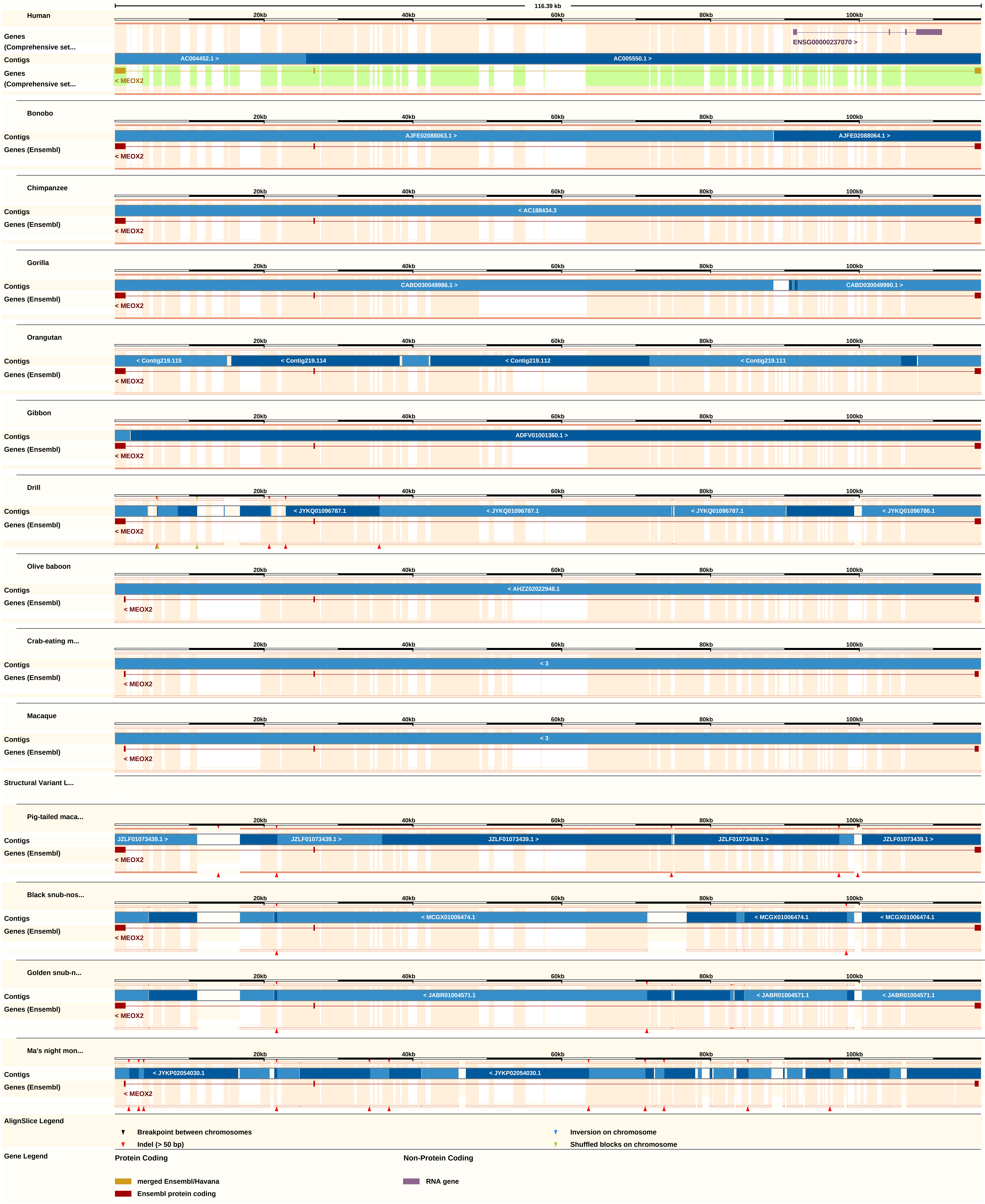
116.39 kb
Human
20kb
40kb
60kb
80kb
100kb
Genes (Comprehensive set...
ENSG00000237070 >
Contigs
AC004452.1 >
AC005550.1 >
Genes (Comprehensive set...
< MEOX2
Bonobo
Contigs
AJFE02088063.1 >
AJFE02088064.1 >
Genes (Ensembl)
< MEOX2
Chimpanzee
Contigs
< AC188434.3
Genes (Ensembl)
< MEOX2
Gorilla
Contigs
CABD030049986.1 >
CABD030049990.1 >
Genes (Ensembl)
< MEOX2
Orangutan
Contigs
< Contig219.115
< Contig219.114
< Contig219.112
< Contig219.111
Genes (Ensembl)
< MEOX2
Gibbon
Contigs
ADFV01001360.1 >
Genes (Ensembl)
< MEOX2
Drill
Contigs
< JYKQ01096787.1
< JYKQ01096787.1
< JYKQ01096787.1
< JYKQ01096786.1
Genes (Ensembl)
< MEOX2
Olive baboon
Contigs
< AHZZ02022948.1
Genes (Ensembl)
< MEOX2
Crab-eating m...
Contigs
< 3
Genes (Ensembl)
< MEOX2
Macaque
Contigs
< 3
Genes (Ensembl)
< MEOX2
Structural Variant L...
Pig-tailed maca...
Contigs
JZLF01073439.1 >
JZLF01073439.1 >
JZLF01073439.1 >
JZLF01073439.1 >
JZLF01073439.1 >
Genes (Ensembl)
< MEOX2
Black snub-nos...
Contigs
< MCGX01006474.1
< MCGX01006474.1
< MCGX01006474.1
Genes (Ensembl)
< MEOX2
Golden snub-n...
Contigs
< JABR01004571.1
< JABR01004571.1
< JABR01004571.1
Genes (Ensembl)
< MEOX2
Ma's night mon...
Contigs
< JYKP02054030.1
< JYKP02054030.1
Genes (Ensembl)
< MEOX2
AlignSlice Legend
Breakpoint between chromosomes
Indel (> 50 bp)
Inversion on chromosome
Shuffled blocks on chromosome
Gene Legend
Protein Coding
merged Ensembl/Havana
Ensembl protein coding
Non-Protein Coding
RNA gene

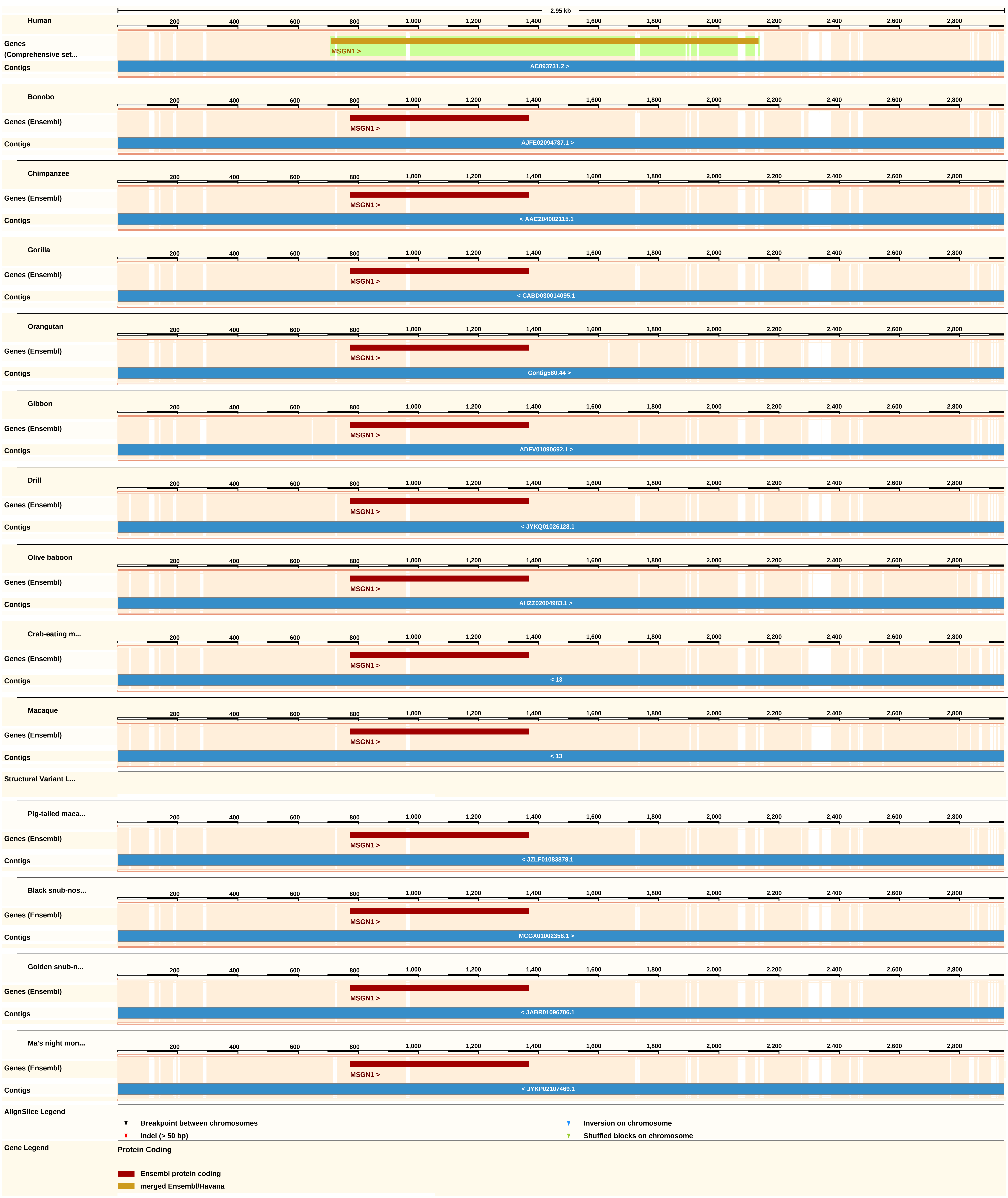
2.95 kb
200
400
600
800
1,000
1,200
1,400
1,600
1,800
2,000
2,200
2,400
2,600
2,800
Human
Genes (Comprehensive set...
MSGN1 >
Contigs
AC093731.2 >
Bonobo
Genes (Ensembl)
MSGN1 >
Contigs
AJFE02094787.1 >
Chimpanzee
Genes (Ensembl)
MSGN1 >
Contigs
< AACZ04002115.1
Gorilla
Genes (Ensembl)
MSGN1 >
Contigs
< CABD030014095.1
Orangutan
Genes (Ensembl)
MSGN1 >
Contigs
Contig580.44 >
Gibbon
Genes (Ensembl)
MSGN1 >
Contigs
ADFV01090692.1 >
Drill
Genes (Ensembl)
MSGN1 >
Contigs
< JYKQ01026128.1
Olive baboon
Genes (Ensembl)
MSGN1 >
Contigs
AHZZ02004983.1 >
Crab-eating m...
Genes (Ensembl)
MSGN1 >
Contigs
< 13
Macaque
Genes (Ensembl)
MSGN1 >
Contigs
< 13
Structural Variant L...
Pig-tailed maca...
Genes (Ensembl)
MSGN1 >
Contigs
< JZLF01083878.1
Black snub-nos...
Genes (Ensembl)
MSGN1 >
Contigs
MCGX01002358.1 >
Golden snub-n...
Genes (Ensembl)
MSGN1 >
Contigs
< JABR01096706.1
Ma's night mon...
Genes (Ensembl)
MSGN1 >
Contigs
< JYKP02107469.1
AlignSlice Legend
Breakpoint between chromosomes
Inversion on chromosome
Indel (> 50 bp)
Shuffled blocks on chromosome
Gene Legend
Protein Coding
Ensembl protein coding
merged Ensembl/Havana

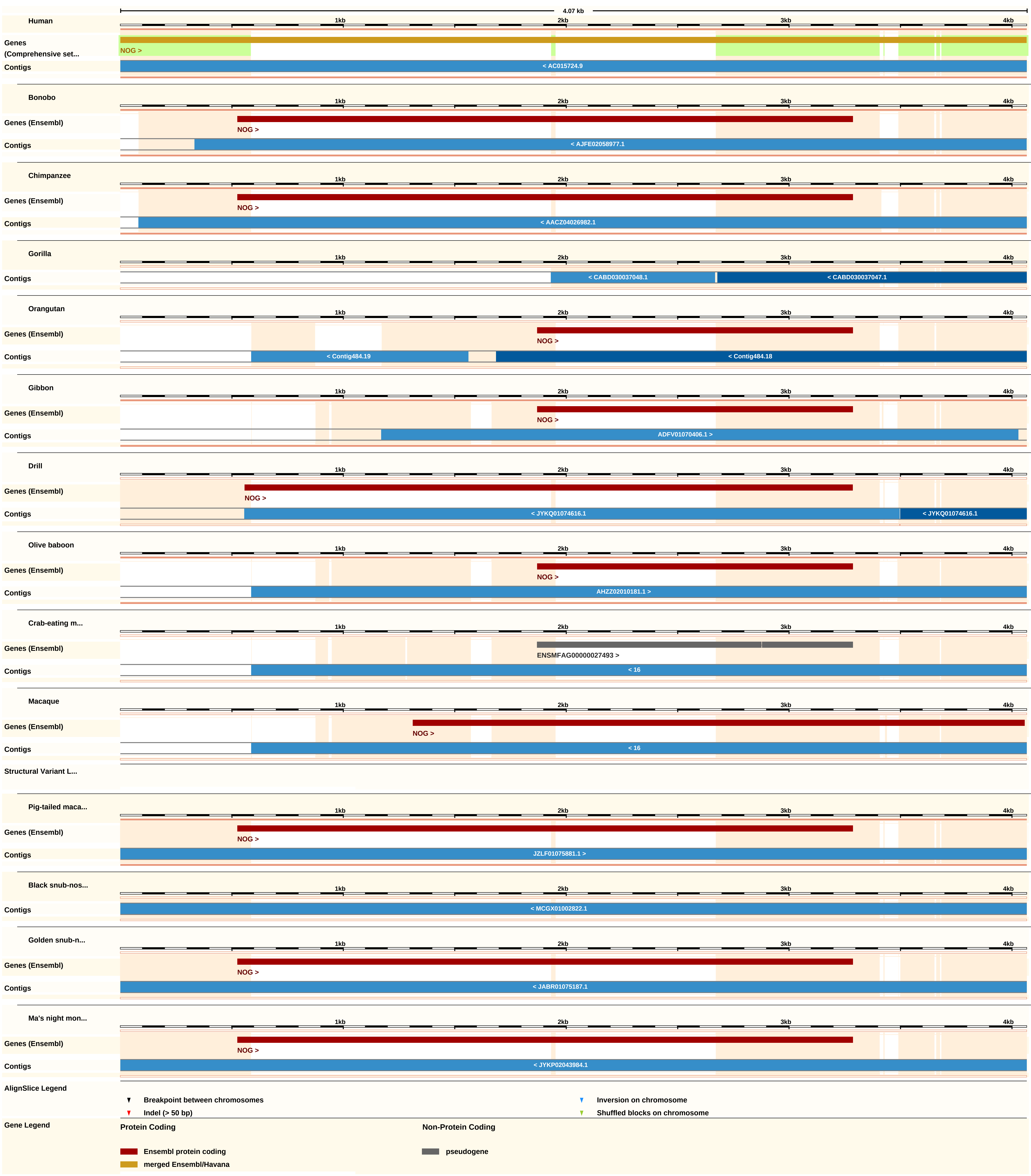
4.07 kb
Human
1kb
2kb
3kb
4kb
Genes (Comprehensive set...
NOG >
Contigs
< AC015724.9
Bonobo
Genes (Ensembl)
NOG >
Contigs
< AJFE02058977.1
Chimpanzee
Genes (Ensembl)
NOG >
Contigs
< AACZ04026982.1
Gorilla
Contigs
< CABD030037048.1
< CABD030037047.1
Orangutan
Genes (Ensembl)
NOG >
Contigs
< Contig484.19
< Contig484.18
Gibbon
Genes (Ensembl)
NOG >
Contigs
ADFV01070406.1 >
Drill
Genes (Ensembl)
NOG >
Contigs
< JYKQ01074616.1
< JYKQ01074616.1
Olive baboon
Genes (Ensembl)
NOG >
Contigs
AHZZ02010181.1 >
Crab-eating m...
Genes (Ensembl)
ENSMFAG00000027493 >
Contigs
< 16
Macaque
Genes (Ensembl)
NOG >
Contigs
< 16
Structural Variant L...
Pig-tailed maca...
Genes (Ensembl)
NOG >
Contigs
JZLF01075881.1 >
Black snub-nos...
Contigs
< MCGX01002822.1
Golden snub-n...
Genes (Ensembl)
NOG >
Contigs
< JABR01075187.1
Ma's night mon...
Genes (Ensembl)
NOG >
Contigs
< JYKP02043984.1
AlignSlice Legend
Breakpoint between chromosomes
Indel (> 50 bp)
Inversion on chromosome
Shuffled blocks on chromosome
Gene Legend
Protein Coding
Ensembl protein coding
merged Ensembl/Havana
Non-Protein Coding
pseudogene

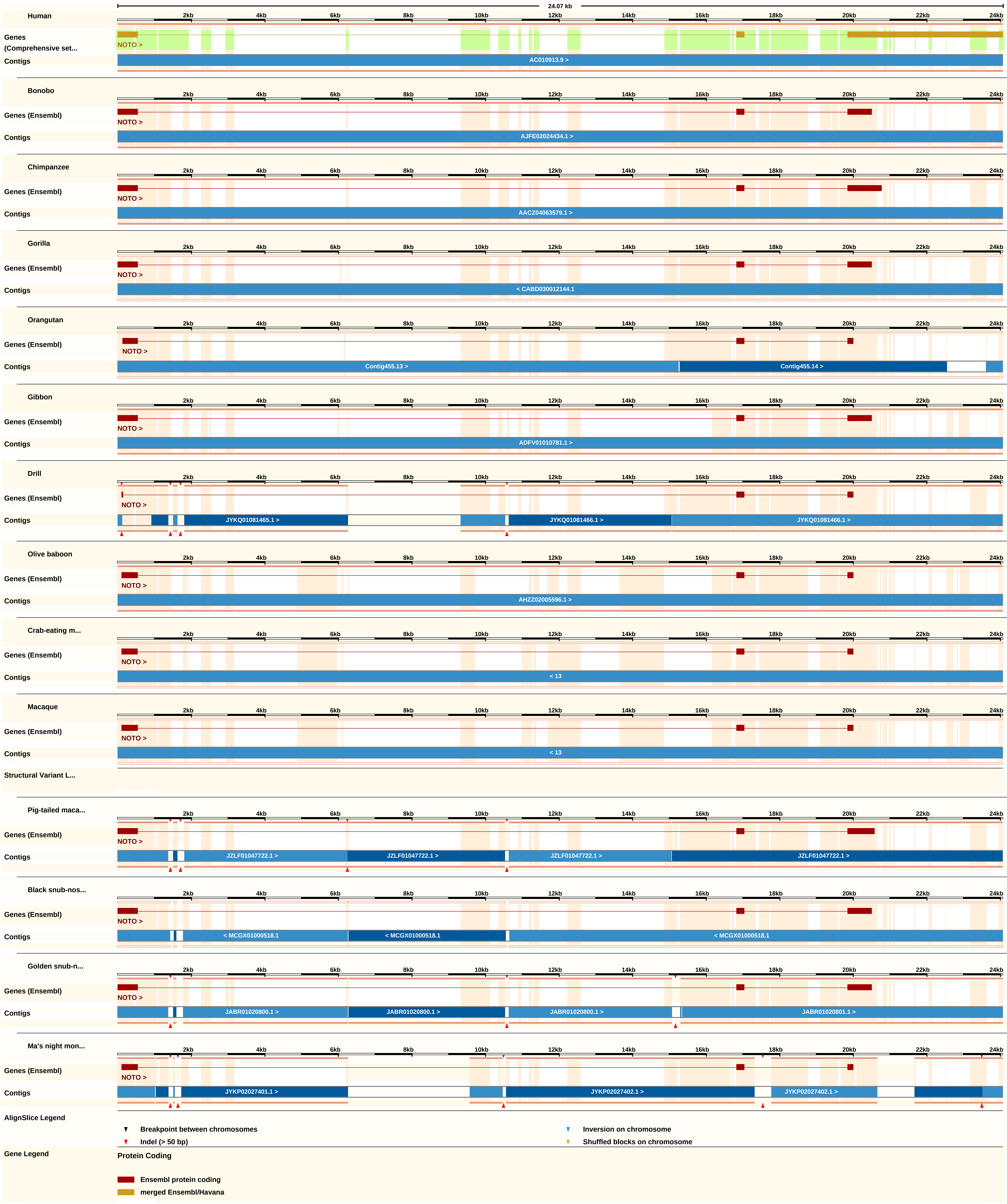

24.07 kb
2kb 4kb 6kb 8kb 10kb 12kb 14kb 16kb 18kb 20kb 22kb 24kb
Human
Genes (Comprehensive set...
NOTO >
Contigs
AC010913.9 >
Bonobo
Genes (Ensembl)
NOTO >
Contigs
AJFE02024434.1 >
Chimpanzee
Genes (Ensembl)
NOTO >
Contigs
AACZ04063579.1 >
Gorilla
Genes (Ensembl)
NOTO >
Contigs
< CABD030012144.1
Orangutan
Genes (Ensembl)
NOTO >
Contigs
Contig455.13 >
Contig455.14 >
Gibbon
Genes (Ensembl)
NOTO >
Contigs
ADFV01010781.1 >
Drill
Genes (Ensembl)
NOTO >
Contigs
JYKQ01081465.1 >
JYKQ01081466.1 >
JYKQ01081466.1 >
Olive baboon
Genes (Ensembl)
NOTO >
Contigs
AHZZ02005596.1 >
Crab-eating m...
Genes (Ensembl)
NOTO >
Contigs
< 13
Macaque
Genes (Ensembl)
NOTO >
Contigs
< 13
Structural Variant L...
Pig-tailed maca...
Genes (Ensembl)
NOTO >
Contigs
JZLF01047722.1 >
JZLF01047722.1 >
JZLF01047722.1 >
JZLF01047722.1 >
Black snub-nos...
Genes (Ensembl)
NOTO >
Contigs
< MCGX01000518.1
< MCGX01000518.1
< MCGX01000518.1
Golden snub-n...
Genes (Ensembl)
NOTO >
Contigs
JABR01020800.1 >
JABR01020800.1 >
JABR01020800.1 >
JABR01020801.1 >
Ma's night mon...
Genes (Ensembl)
NOTO >
Contigs
JYKP02027401.1 >
JYKP02027402.1 >
JYKP02027402.1 >
AlignSlice Legend
Breakpoint between chromosomes
Inversion on chromosome
Indel (> 50 bp)
Shuffled blocks on chromosome
Gene Legend
Protein Coding
Ensembl protein coding
merged Ensembl/Havana

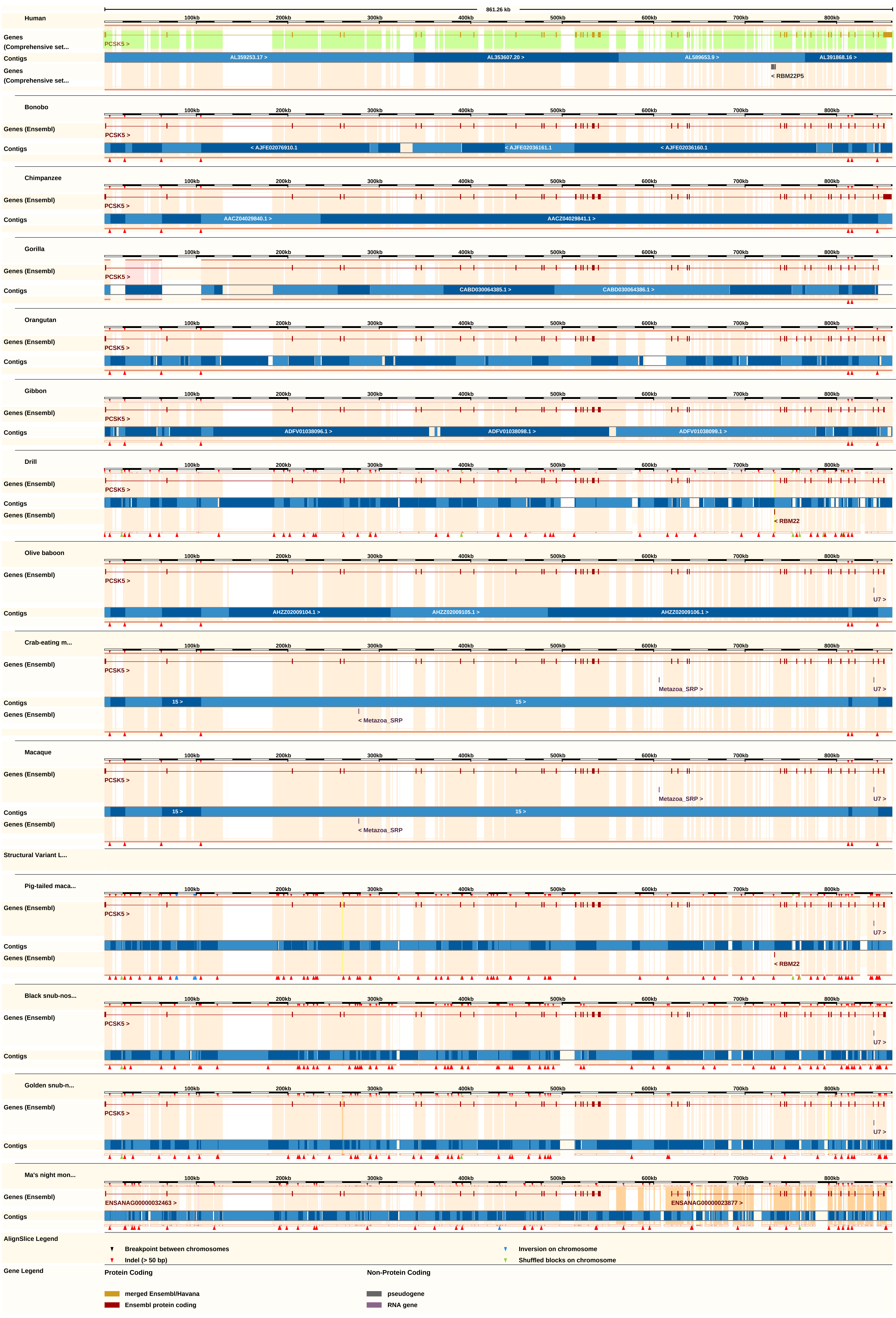

861.26 kb
100kb
200kb
300kb
400kb
500kb
600kb
700kb
800kb
Human
Genes
(Comprehensive set...
PCSK5 >
Contigs
AL359253.17 >
AL353607.20 >
AL589653.9 >
AL391868.16 >
< RBM22P5
Bonobo
Genes (Ensembl)
< AJFE02076910.1
< AJFE02036161.1
< AJFE02036160.1
Chimpanzee
AACZ04029840.1 >
AACZ04029841.1 >
Gorilla
CABD030064385.1 >
CABD030064386.1 >
Orangutan
Gibbon
ADFV01038096.1 >
ADFV01038098.1 >
ADFV01038099.1 >
Drill
< RBM22
Olive baboon
U7 >
AHZZ02009104.1 >
AHZZ02009105.1 >
AHZZ02009106.1 >
Crab-eating m...
Metazoa_SRP >
15 >
< Metazoa_SRP
Macaque
Structural Variant L...
Pig-tailed maca...
Black snub-nos...
Golden snub-n...
Ma's night mon...
ENSANAG00000032463 >
ENSANAG00000023877 >
AlignSlice Legend
Breakpoint between chromosomes
Indel (> 50 bp)
Inversion on chromosome
Shuffled blocks on chromosome
Gene Legend
Protein Coding
merged Ensembl/Havana
Ensembl protein coding
Non-Protein Coding
pseudogene
RNA gene

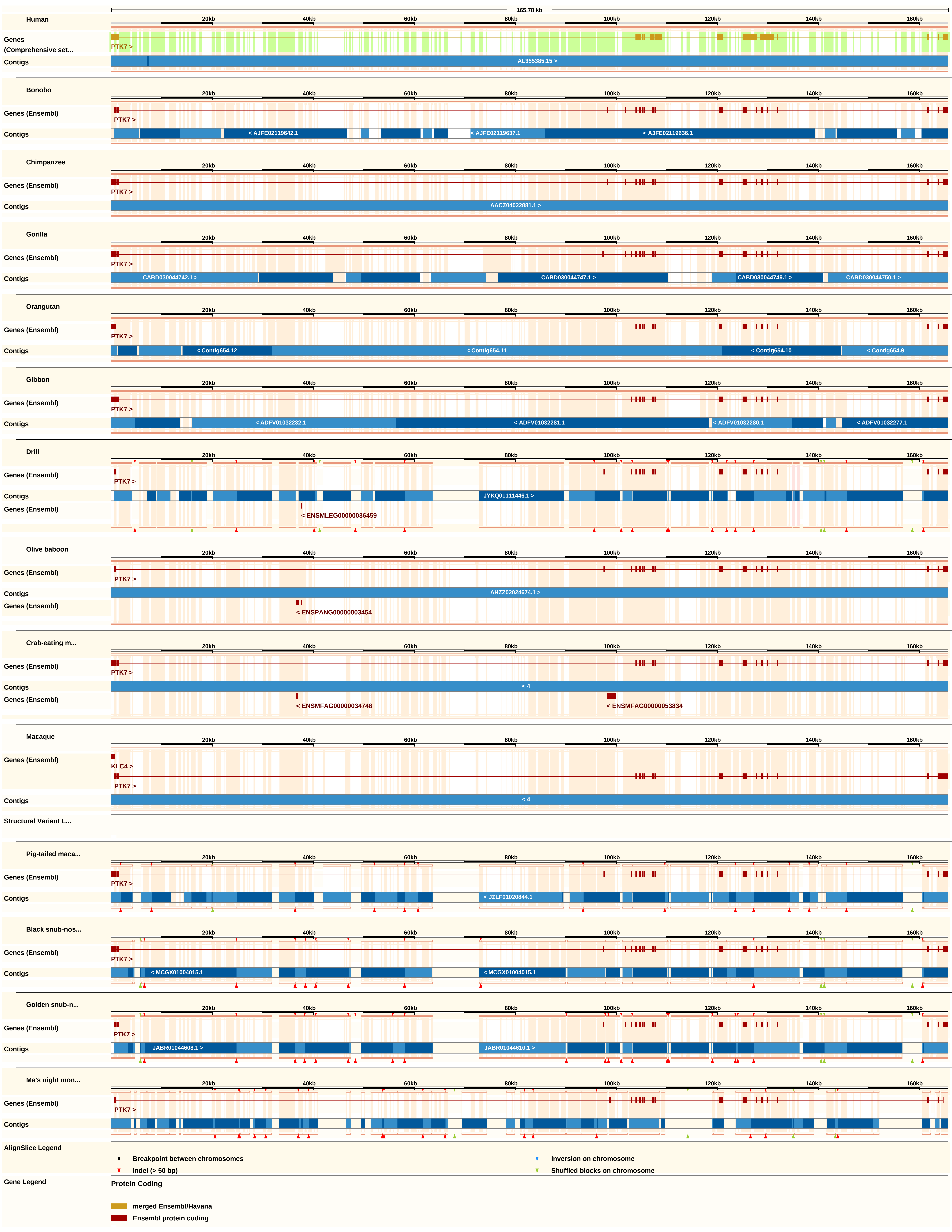
165.78 kb
20kb
40kb
60kb
80kb
100kb
120kb
140kb
160kb
Human
Genes
(Comprehensive set...
PTK7 >
Contigs
AL355385.15 >
Bonobo
Genes (Ensembl)
PTK7 >
Contigs
< AJFE02119642.1
< AJFE02119637.1
< AJFE02119636.1
Chimpanzee
Genes (Ensembl)
PTK7 >
Contigs
AACZ04022881.1 >
Gorilla
Genes (Ensembl)
PTK7 >
Contigs
CABD030044742.1 >
CABD030044747.1 >
CABD030044749.1 >
CABD030044750.1 >
Orangutan
Genes (Ensembl)
PTK7 >
Contigs
< Contig654.12
< Contig654.11
< Contig654.10
< Contig654.9
Gibbon
Genes (Ensembl)
PTK7 >
Contigs
< ADFV01032282.1
< ADFV01032281.1
< ADFV01032280.1
< ADFV01032277.1
Drill
Genes (Ensembl)
PTK7 >
Contigs
JYKQ01111446.1 >
Genes (Ensembl)
< ENSMLEG00000036459
Olive baboon
Genes (Ensembl)
PTK7 >
Contigs
AHZZ02024674.1 >
Genes (Ensembl)
< ENSPANG00000003454
Crab-eating m...
Genes (Ensembl)
PTK7 >
Contigs
< 4
Genes (Ensembl)
< ENSMFAG00000034748
< ENSMFAG00000053834
Macaque
Genes (Ensembl)
KLC4 >
PTK7 >
Contigs
< 4
Structural Variant L...
Pig-tailed maca...
Genes (Ensembl)
PTK7 >
Contigs
< JZLF01020844.1
Black snub-nos...
Genes (Ensembl)
PTK7 >
Contigs
< MCGX01004015.1
< MCGX01004015.1
Golden snub-n...
Genes (Ensembl)
PTK7 >
Contigs
JABR01044608.1 >
JABR01044610.1 >
Ma's night mon...
Genes (Ensembl)
PTK7 >
Contigs
AlignSlice Legend
Breakpoint between chromosomes
Inversion on chromosome
Indel (> 50 bp)
Shuffled blocks on chromosome
Gene Legend
Protein Coding
merged Ensembl/Havana
Ensembl protein coding

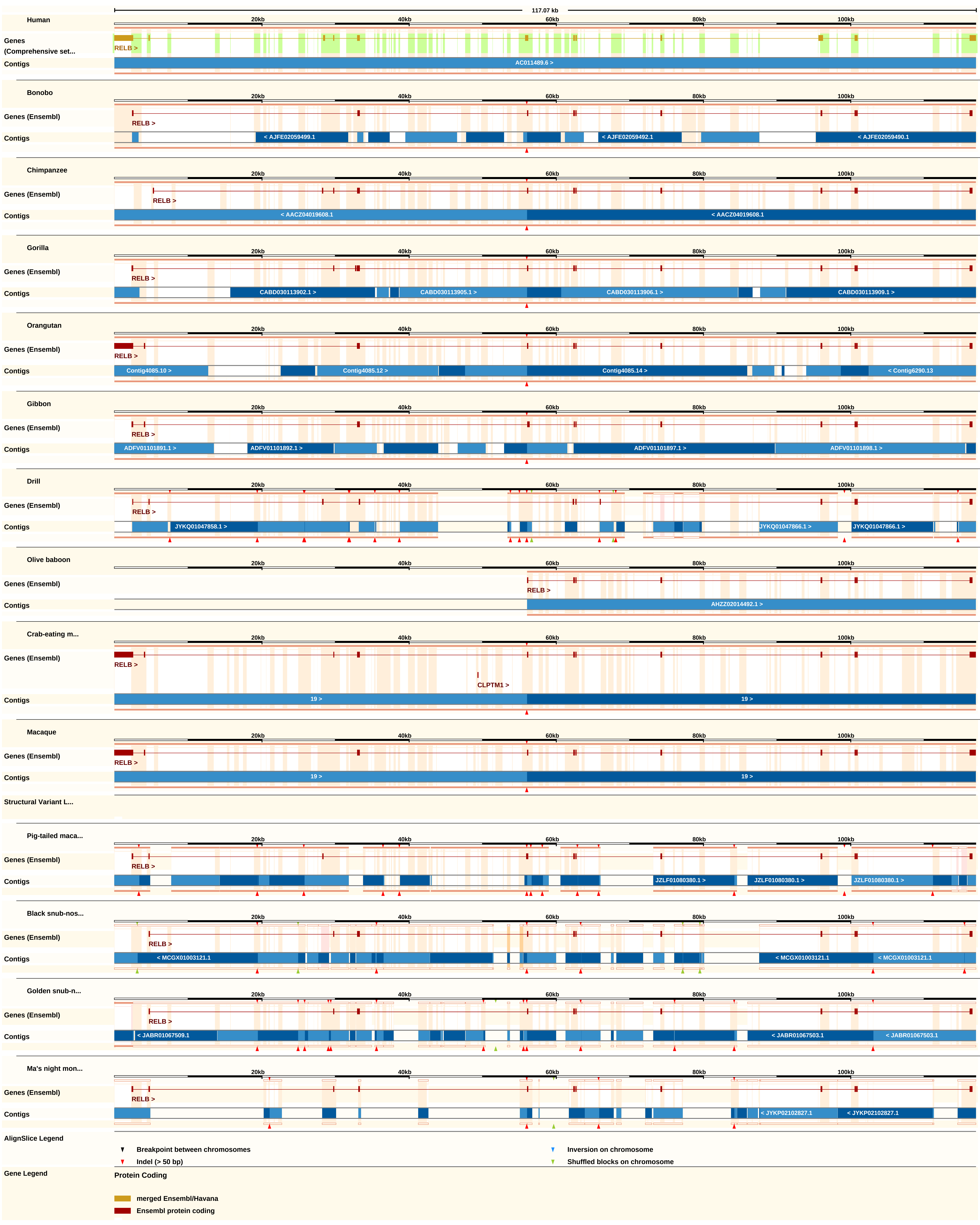

117.07 kb
Human
Genes (Comprehensive set...
RELB >
Contigs
AC011489.6 >
Bonobo
Genes (Ensembl)
RELB >
< AJFE02059499.1
< AJFE02059492.1
< AJFE02059490.1
Chimpanzee
< AACZ04019608.1
Gorilla
CABD030113902.1 >
CABD030113905.1 >
CABD030113906.1 >
CABD030113909.1 >
Orangutan
Contig4085.10 >
Contig4085.12 >
Contig4085.14 >
< Contig6290.13
Gibbon
ADFV01101891.1 >
ADFV01101892.1 >
ADFV01101897.1 >
ADFV01101898.1 >
Drill
JYKQ01047858.1 >
JYKQ01047866.1 >
Olive baboon
AHZZ02014492.1 >
Crab-eating m...
CLPTM1 >
19 >
Macaque
Structural Variant L...
Pig-tailed maca...
JZLF01080380.1 >
Black snub-nos...
< MCGX01003121.1
Golden snub-n...
< JABR01067509.1
< JABR01067503.1
Ma's night mon...
< JYKP02102827.1
20kb
40kb
60kb
80kb
100kb
AlignSlice Legend
Breakpoint between chromosomes
Indel (> 50 bp)
Inversion on chromosome
Shuffled blocks on chromosome
Gene Legend
Protein Coding
merged Ensembl/Havana
Ensembl protein coding

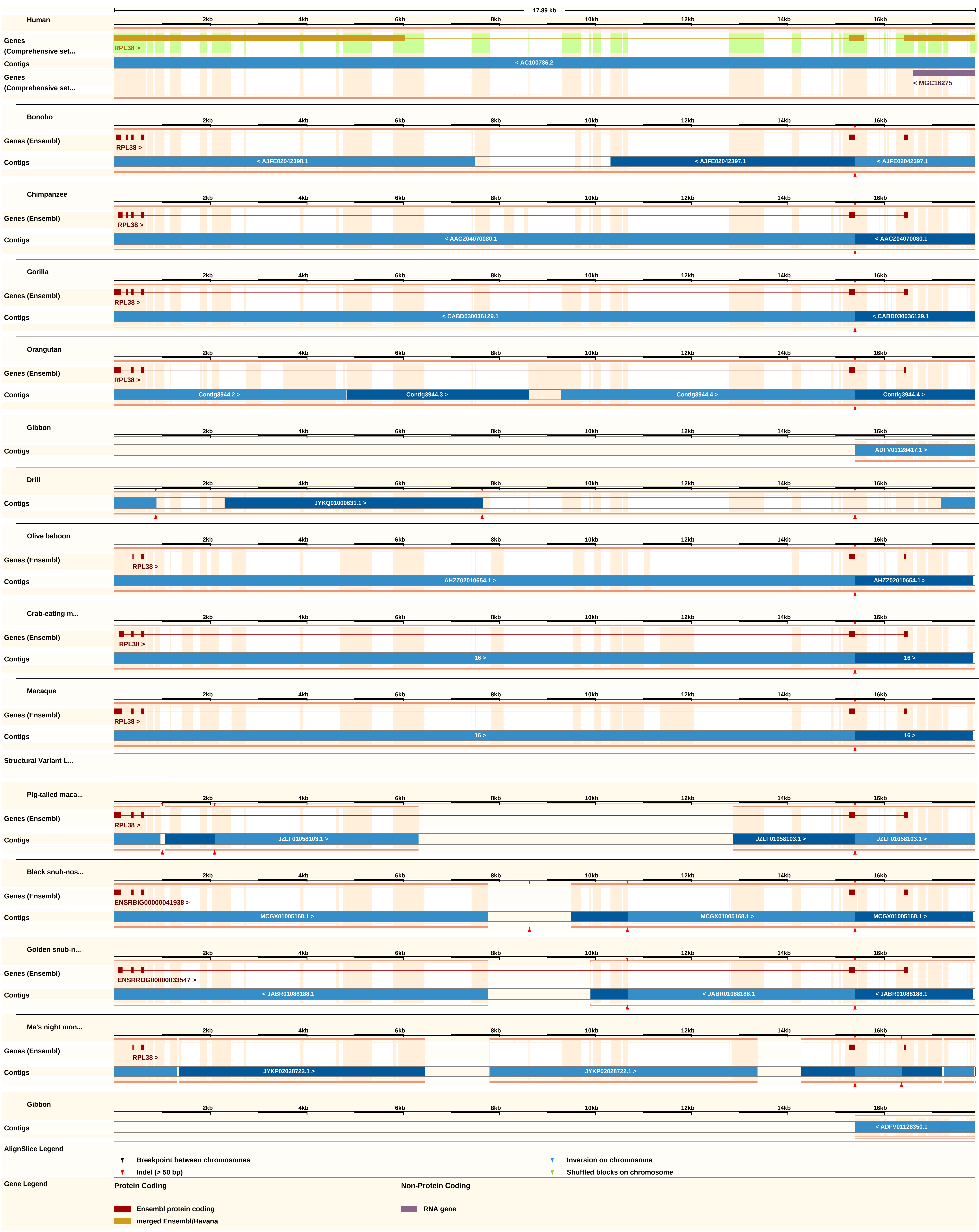
17.89 kb
2kb
4kb
6kb
8kb
10kb
12kb
14kb
16kb
Human
Genes (Comprehensive set...
RPL38 >
Contigs
< AC100786.2
Genes (Comprehensive set...
< MGC16275
Bonobo
Genes (Ensembl)
RPL38 >
Contigs
< AJFE02042398.1
< AJFE02042397.1
< AJFE02042397.1
Chimpanzee
Genes (Ensembl)
RPL38 >
Contigs
< AACZ04070080.1
< AACZ04070080.1
Gorilla
Genes (Ensembl)
RPL38 >
Contigs
< CABD030036129.1
< CABD030036129.1
Orangutan
Genes (Ensembl)
RPL38 >
Contigs
Contig3944.2 >
Contig3944.3 >
Contig3944.4 >
Contig3944.4 >
Gibbon
Contigs
ADFV01128417.1 >
Drill
Contigs
JYKQ01000631.1 >
Olive baboon
Genes (Ensembl)
RPL38 >
Contigs
AHZZ02010654.1 >
AHZZ02010654.1 >
Crab-eating m...
Genes (Ensembl)
RPL38 >
Contigs
16 >
16 >
Macaque
Genes (Ensembl)
RPL38 >
Contigs
16 >
16 >
Structural Variant L...
Pig-tailed maca...
Genes (Ensembl)
RPL38 >
Contigs
JZLF01058103.1 >
JZLF01058103.1 >
JZLF01058103.1 >
Black snub-nos...
Genes (Ensembl)
ENSRBIG00000041938 >
Contigs
MCGX01005168.1 >
MCGX01005168.1 >
MCGX01005168.1 >
Golden snub-n...
Genes (Ensembl)
ENSRROG00000033547 >
Contigs
< JABR01088188.1
< JABR01088188.1
< JABR01088188.1
Ma's night mon...
Genes (Ensembl)
RPL38 >
Contigs
JYKP02028722.1 >
JYKP02028722.1 >
Gibbon
Contigs
< ADFV01128350.1
AlignSlice Legend
Breakpoint between chromosomes
Indel (> 50 bp)
Inversion on chromosome
Shuffled blocks on chromosome
Gene Legend
Protein Coding
Ensembl protein coding
merged Ensembl/Havana
Non-Protein Coding
RNA gene

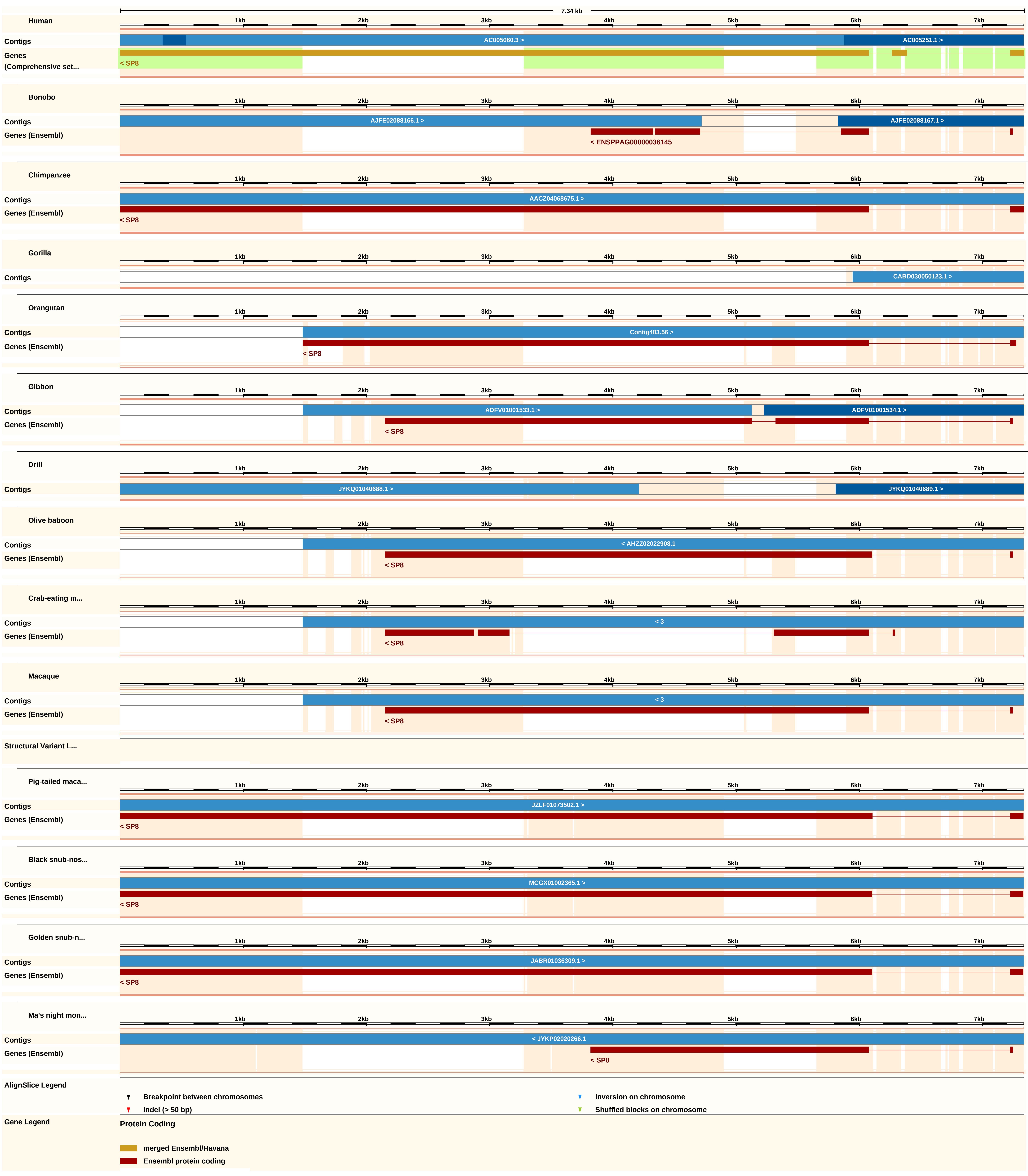

7.34 kb
Human
Contigs
AC005060.3 >
AC005251.1 >
Genes (Comprehensive set...
< SP8
Bonobo
Contigs
AJFE02088166.1 >
AJFE02088167.1 >
Genes (Ensembl)
< ENSPPAG00000036145
Chimpanzee
Contigs
AACZ04068675.1 >
Genes (Ensembl)
< SP8
Gorilla
Contigs
CABD030050123.1 >
Orangutan
Contigs
Contig483.56 >
Genes (Ensembl)
< SP8
Gibbon
Contigs
ADFV01001533.1 >
ADFV01001534.1 >
Genes (Ensembl)
< SP8
Drill
Contigs
JYKQ01040688.1 >
JYKQ01040689.1 >
Olive baboon
Contigs
< AHZZ02022908.1
Genes (Ensembl)
< SP8
Crab-eating m...
Contigs
< 3
Genes (Ensembl)
< SP8
Macaque
Contigs
< 3
Genes (Ensembl)
< SP8
Structural Variant L...
Pig-tailed maca...
Contigs
JZLF01073502.1 >
Genes (Ensembl)
< SP8
Black snub-nos...
Contigs
MCGX01002365.1 >
Genes (Ensembl)
< SP8
Golden snub-n...
Contigs
JABR01036309.1 >
Genes (Ensembl)
< SP8
Ma's night mon...
Contigs
< JYKP02020266.1
Genes (Ensembl)
< SP8
1kb 2kb 3kb 4kb 5kb 6kb 7kb
AlignSlice Legend
Breakpoint between chromosomes
Inversion on chromosome
Indel (> 50 bp)
Shuffled blocks on chromosome
Gene Legend
Protein Coding
merged Ensembl/Havana
Ensembl protein coding

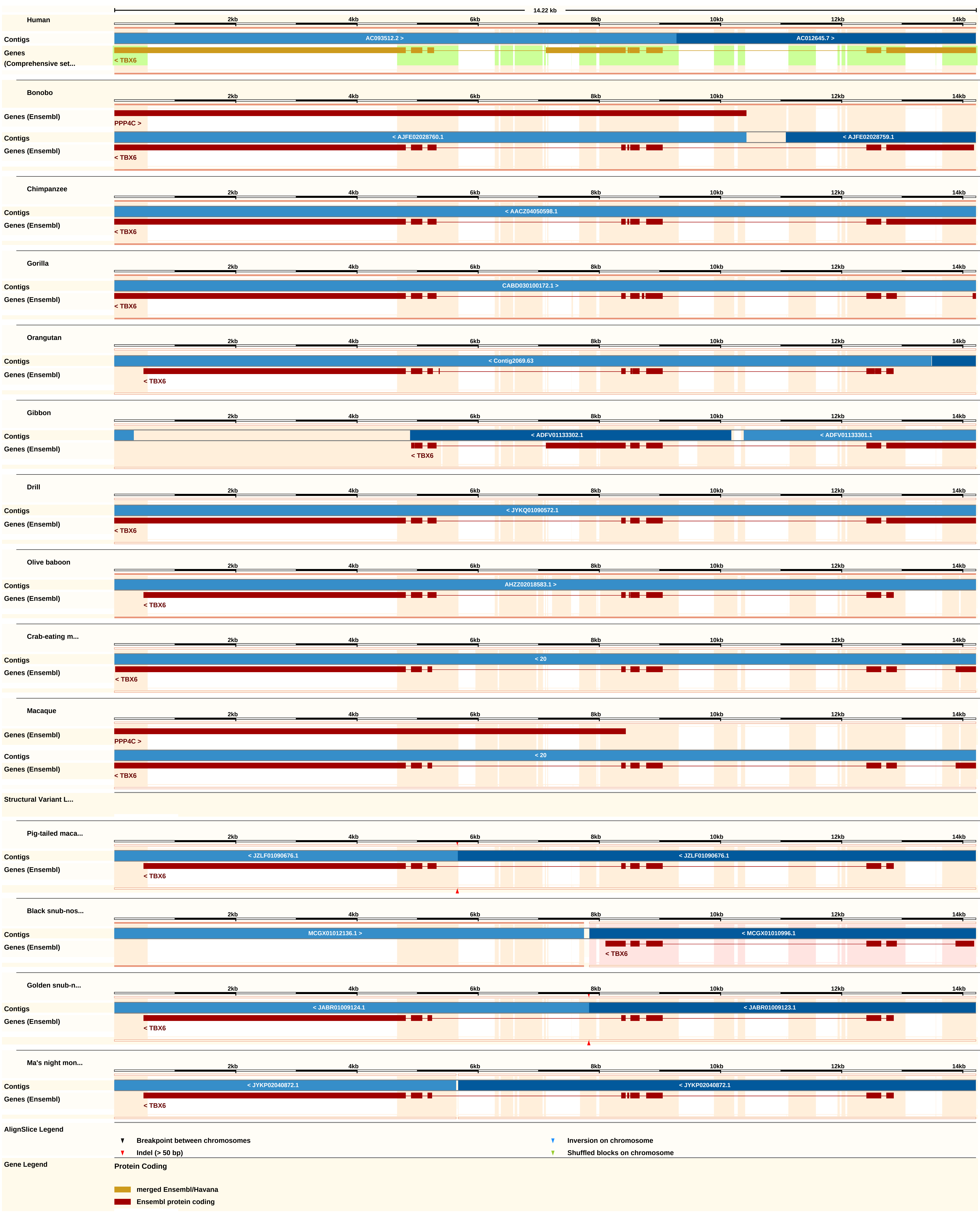
14.22 kb
2kb 4kb 6kb 8kb 10kb 12kb 14kb
Human
Contigs
AC093512.2 >
AC012645.7 >
Genes (Comprehensive set...
< TBX6
Bonobo
Genes (Ensembl)
PPP4C >
Contigs
< AJFE02028760.1
< AJFE02028759.1
Genes (Ensembl)
< TBX6
Chimpanzee
Contigs
< AACZ04050598.1
Genes (Ensembl)
< TBX6
Gorilla
Contigs
CABD030100172.1 >
Genes (Ensembl)
< TBX6
Orangutan
Contigs
< Contig2069.63
Genes (Ensembl)
< TBX6
Gibbon
Contigs
< ADFV01133302.1
< ADFV01133301.1
Genes (Ensembl)
< TBX6
Drill
Contigs
< JYKQ01090572.1
Genes (Ensembl)
< TBX6
Olive baboon
Contigs
AHZZ02018583.1 >
Genes (Ensembl)
< TBX6
Crab-eating m...
Contigs
< 20
Genes (Ensembl)
< TBX6
Macaque
Genes (Ensembl)
PPP4C >
Contigs
< 20
Genes (Ensembl)
< TBX6
Structural Variant L...
Pig-tailed maca...
Contigs
< JZLF01090676.1
< JZLF01090676.1
Genes (Ensembl)
< TBX6
Black snub-nos...
Contigs
MCGX01012136.1 >
< MCGX01010996.1
Genes (Ensembl)
< TBX6
Golden snub-n...
Contigs
< JABR01009124.1
< JABR01009123.1
Genes (Ensembl)
< TBX6
Ma's night mon...
Contigs
< JYKP02040872.1
< JYKP02040872.1
Genes (Ensembl)
< TBX6
AlignSlice Legend
Breakpoint between chromosomes
Inversion on chromosome
Indel (> 50 bp)
Shuffled blocks on chromosome
Gene Legend
Protein Coding
merged Ensembl/Havana
Ensembl protein coding

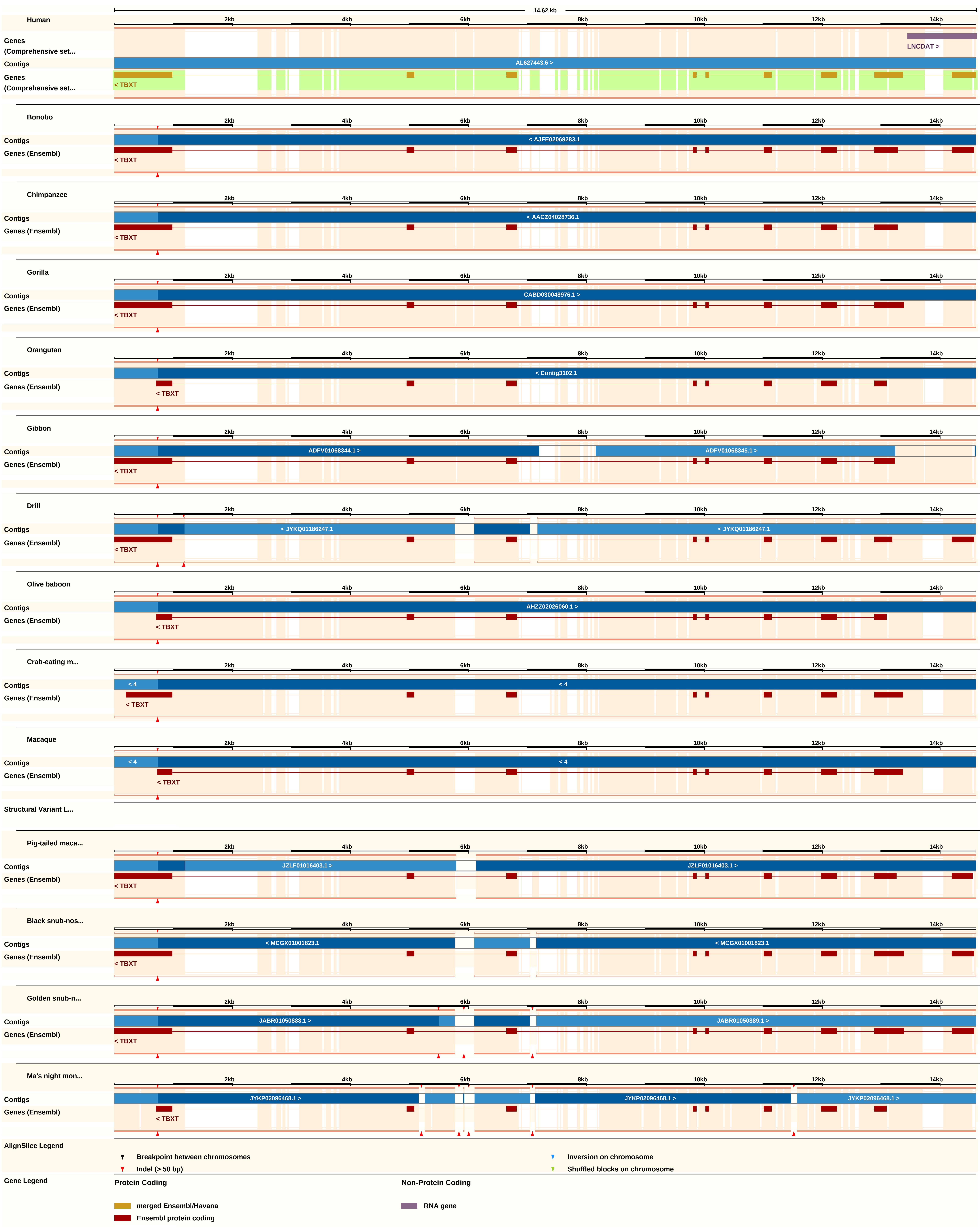
14.62 kb
Human
2kb
4kb
6kb
8kb
10kb
12kb
14kb
Genes (Comprehensive set...
LNCDAT >
Contigs
AL627443.6 >
Genes (Comprehensive set...
< TBXT
Bonobo
Contigs
< AJFE02069283.1
Genes (Ensembl)
< TBXT
Chimpanzee
Contigs
< AACZ04028736.1
Genes (Ensembl)
< TBXT
Gorilla
Contigs
CABD030048976.1 >
Genes (Ensembl)
< TBXT
Orangutan
Contigs
< Contig3102.1
Genes (Ensembl)
< TBXT
Gibbon
Contigs
ADFV01068344.1 >
ADFV01068345.1 >
Genes (Ensembl)
< TBXT
Drill
Contigs
< JYKQ01186247.1
< JYKQ01186247.1
Genes (Ensembl)
< TBXT
Olive baboon
Contigs
AHZZ02026060.1 >
Genes (Ensembl)
< TBXT
Crab-eating m...
Contigs
< 4
< 4
Genes (Ensembl)
< TBXT
Macaque
Contigs
< 4
< 4
Genes (Ensembl)
< TBXT
Structural Variant L...
Pig-tailed maca...
Contigs
JZLF01016403.1 >
JZLF01016403.1 >
Genes (Ensembl)
< TBXT
Black snub-nos...
Contigs
< MCGX01001823.1
< MCGX01001823.1
Genes (Ensembl)
< TBXT
Golden snub-n...
Contigs
JABR01050888.1 >
JABR01050889.1 >
Genes (Ensembl)
< TBXT
Ma's night mon...
Contigs
JYKP02096468.1 >
JYKP02096468.1 >
JYKP02096468.1 >
Genes (Ensembl)
< TBXT
AlignSlice Legend
Breakpoint between chromosomes
Indel (> 50 bp)
Inversion on chromosome
Shuffled blocks on chromosome
Gene Legend
Protein Coding
merged Ensembl/Havana
Ensembl protein coding
Non-Protein Coding
RNA gene

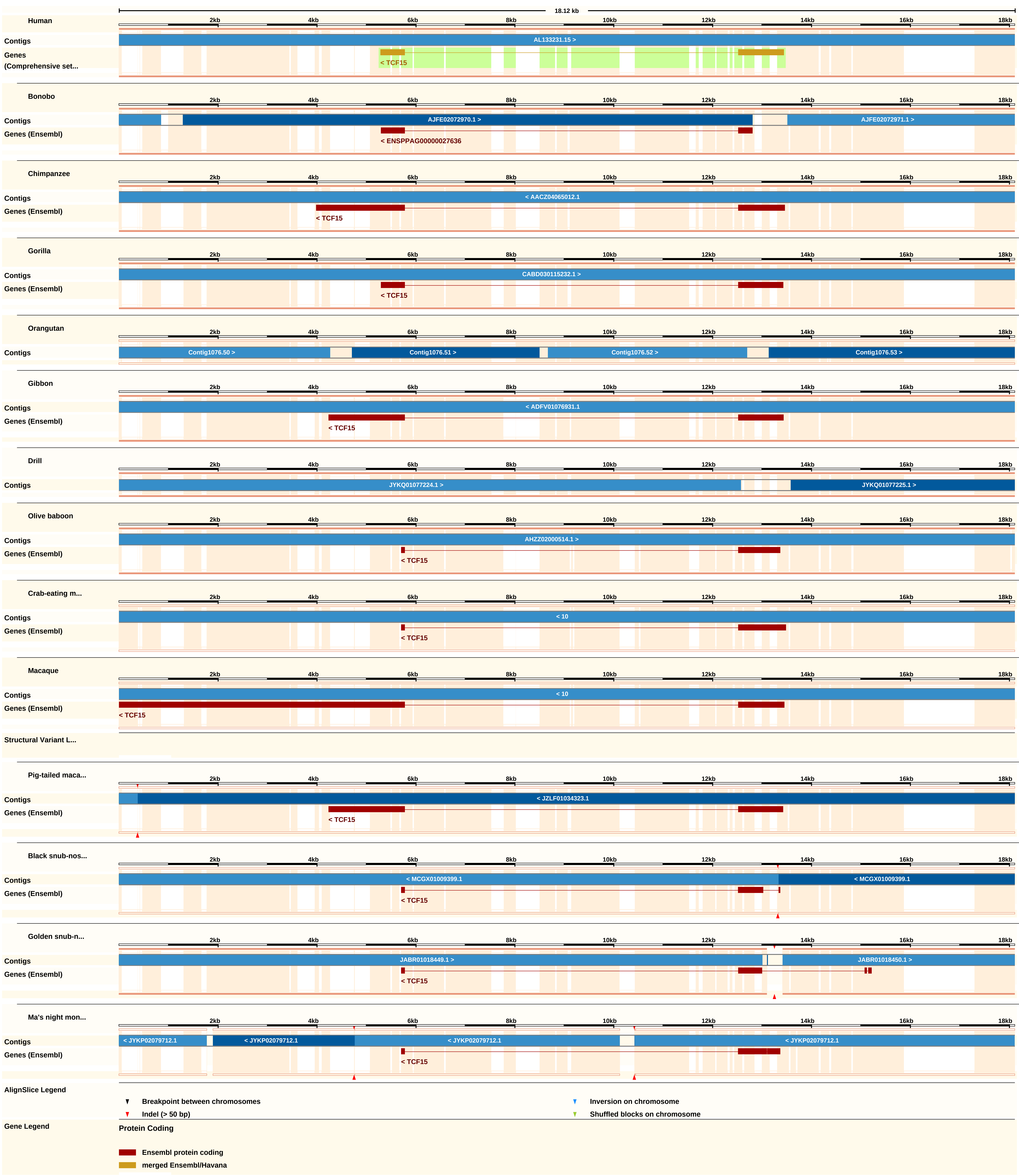
18.12 kb
2kb
4kb
6kb
8kb
10kb
12kb
14kb
16kb
18kb
Human
Contigs
AL133231.15 >
Genes
(Comprehensive set...
< TCF15
Bonobo
Contigs
AJFE02072970.1 >
AJFE02072971.1 >
Genes (Ensembl)
< ENSPPAG00000027636
Chimpanzee
Contigs
< AACZ04065012.1
Genes (Ensembl)
< TCF15
Gorilla
Contigs
CABD030115232.1 >
Genes (Ensembl)
< TCF15
Orangutan
Contigs
Contig1076.50 >
Contig1076.51 >
Contig1076.52 >
Contig1076.53 >
Gibbon
Contigs
< ADFV01076931.1
Genes (Ensembl)
< TCF15
Drill
Contigs
JYKQ01077224.1 >
JYKQ01077225.1 >
Olive baboon
Contigs
AHZZ02000514.1 >
Genes (Ensembl)
< TCF15
Crab-eating m...
Contigs
< 10
Genes (Ensembl)
< TCF15
Macaque
Contigs
< 10
Genes (Ensembl)
< TCF15
Structural Variant L...
Pig-tailed maca...
Contigs
< JZLF01034323.1
Genes (Ensembl)
< TCF15
Black snub-nos...
Contigs
< MCGX01009399.1
< MCGX01009399.1
Genes (Ensembl)
< TCF15
Golden snub-n...
Contigs
JABR01018449.1 >
JABR01018450.1 >
Genes (Ensembl)
< TCF15
Ma's night mon...
Contigs
< JYKP02079712.1
< JYKP02079712.1
< JYKP02079712.1
< JYKP02079712.1
Genes (Ensembl)
< TCF15
AlignSlice Legend
Breakpoint between chromosomes
Inversion on chromosome
Indel (> 50 bp)
Shuffled blocks on chromosome
Gene Legend
Protein Coding
Ensembl protein coding
merged Ensembl/Havana

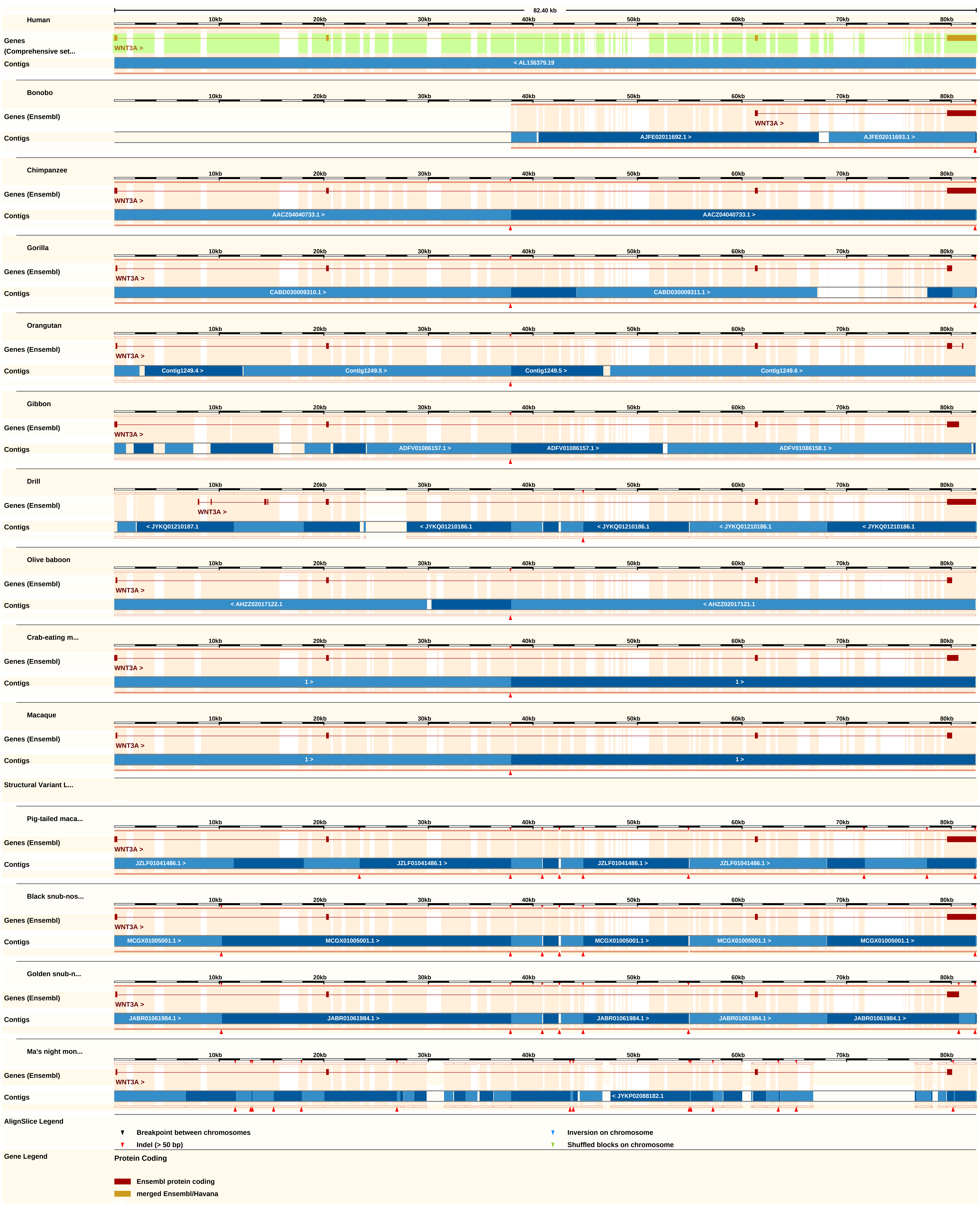

82.40 kb
10kb
20kb
30kb
40kb
50kb
60kb
70kb
80kb
Human
Genes (Comprehensive set...
WNT3A >
Contigs
< AL136379.19
Bonobo
Genes (Ensembl)
WNT3A >
Contigs
AJFE02011692.1 >
AJFE02011693.1 >
Chimpanzee
Genes (Ensembl)
WNT3A >
Contigs
AACZ04040733.1 >
AACZ04040733.1 >
Gorilla
Genes (Ensembl)
WNT3A >
Contigs
CABD030009310.1 >
CABD030009311.1 >
Orangutan
Genes (Ensembl)
WNT3A >
Contigs
Contig1249.4 >
Contig1249.5 >
Contig1249.5 >
Contig1249.6 >
Gibbon
Genes (Ensembl)
WNT3A >
Contigs
ADFV01086157.1 >
ADFV01086157.1 >
ADFV01086158.1 >
Drill
Genes (Ensembl)
WNT3A >
Contigs
< JYKQ01210187.1
< JYKQ01210186.1
< JYKQ01210186.1
< JYKQ01210186.1
< JYKQ01210186.1
Olive baboon
Genes (Ensembl)
WNT3A >
Contigs
< AHZZ02017122.1
< AHZZ02017121.1
Crab-eating m...
Genes (Ensembl)
WNT3A >
Contigs
1 >
1 >
Macaque
Genes (Ensembl)
WNT3A >
Contigs
1 >
1 >
Structural Variant L...
Pig-tailed maca...
Genes (Ensembl)
WNT3A >
Contigs
JZLF01041486.1 >
JZLF01041486.1 >
JZLF01041486.1 >
JZLF01041486.1 >
Black snub-nos...
Genes (Ensembl)
WNT3A >
Contigs
MCGX01005001.1 >
MCGX01005001.1 >
MCGX01005001.1 >
MCGX01005001.1 >
MCGX01005001.1 >
Golden snub-n...
Genes (Ensembl)
WNT3A >
Contigs
JABR01061984.1 >
JABR01061984.1 >
JABR01061984.1 >
JABR01061984.1 >
JABR01061984.1 >
Ma's night mon...
Genes (Ensembl)
WNT3A >
Contigs
< JYKP02088182.1
AlignSlice Legend
Breakpoint between chromosomes
Inversion on chromosome
Indel (> 50 bp)
Shuffled blocks on chromosome
Gene Legend
Protein Coding
Ensembl protein coding
merged Ensembl/Havana
